# Disentangled generative modeling of single-cell perturbation responses across dynamic cellular states

**DOI:** 10.64898/2026.11.07.765083

**Authors:** Yue Wang, Yueshan Zhao, Sihan Li, Da Yang

## Abstract

Single-cell perturbation studies are increasingly conducted across complex, multi-condition biological systems, offering a powerful lens into how interventions reshape cell states and molecular programs. However, standard computational models compress single-cell profiles into unified representations where treatment effects are obscured by cell division, lineage progression, donor variation, and cellular stress. Here, we present Disentangled Variational Inference for Perturbation Response (PertVI), a deep generative model that separates baseline mitotic phase, cellular state, and batch effects from treatment-induced shifts while isolating aberrant off-manifold responses in orthogonal anomaly space. This structured design tackles two inter-twined tasks: (1) quantifying treatment-induced changes in cell-state composition, and (2) inferring state-gated transcriptional changes in response to treatment. Across simulated bench-marks, PertVI accurately disentangles coupled biological axes and recovers weak treatment signatures. In conditional chemical and genetic screens of primary human CD8+ and CD4+ T cells, PertVI separates memory-precursor adjuvants from toxic phenocopies, resolves co-clus-tered regulators into directional kinetic trajectories and biphasic rebounders obscured by standard workflows, and reveals that treatment-induced phenotypic shifting is decoupled from broad transcriptional perturbation responses and instead governed by state-specific impulses. Altogether, PertVI brings unique mechanistic granularity to single-cell perturbation response modeling, serving as a proof of principle for biology-informed, structure-guided variance disentan-glement in dynamic cellular systems.

## INTRODUCTION

Over the past decade, single-cell perturbation screens have rapidly expanded from uniform cell lines^1–8^ into dynamic, multi-condition biological systems^9–15^, providing unprecedented opportunities to decode how gene regulatory networks, transcriptional regulons, and molecular interactomes rewire across physiological contexts. Realizing the full potential of these multilayered perturbation screens, however, requires rethinking how to leverage the expressive capacity of deep architectures to uncover the complex biological relationships these datasets are built to reveal. Early analytical frameworks relied on a static cellular baseline where treatment effects dominate, whereas in dynamic, multi-condition systems, localized perturbation shifts^15–17^ are overshadowed by macroscopic state transitions, periodic cell-cycle oscillations^18–23^, and donor or batch variation^17,24–28^. Moreover, these sources of variation are usually biologically intertwined: proliferation covaries with lineage progression and activation, while perturbations simultaneously alter division rates, shift cell-state composition, modulate state-dependent gene programs, and trigger cellular stress^2,6,7,13,15^. This pervasive variance entanglement poses fundamental challenges for perturbation analysis in modern multi-condition screens.

Interrogating dynamic, multi-condition perturbation screens centers on two fundamental analytical tasks. The first task is to determine whether an intervention pushes, arrests, or redirects cells along permissible biological trajectories, such as accelerating differentiation, halting mitotic progression, or shifting functional state occupancy across developing tissues and primary cohorts^9,10,12–14^. The second task is to resolve how a cell’s underlying baseline state or environmental context govern its transcriptional response to a perturbation, uncovering regulatory circuits that act selectively across developmental stages, co-cultures, or stimulation regimes^6,11,15,29^. Accomplishing both tasks amid confounding proliferation and non-physiological toxicity exposes three core challenges that monolithic latent embeddings^30–34^ and purely unsupervised factorization models^19,35–37^ struggle to resolve.

First, isolating perturbation effects in dynamic systems requires an unconfounded baseline representation that separates periodic cell division from progressive differentiation or activation states. Standard preprocessing pipelines score canonical cell-cycle marker sets^18,38^ and regress them out of expression matrices using linear or empirical Bayes models^24,39,40^. Whenever proliferation covaries with state transitions, orthogonal linear projection subtracts correlated developmental signals from pleiotropic metabolic and signaling genes, distorting the underlying state trajectory^21,22,27^. Conversely, specialized circular projection algorithms reconstruct continuous mitotic phase angles on two-dimensional ring manifolds^20,22^, but operate as standalone projections that neither infer a deconfounded cellular state manifold nor model perturbation effect. Meanwhile, deep generative frameworks based on monolithic variational autoencoders^30,41^, unsupervised factorized bottlenecks^37^, and contrastive or multi-resolution architectures^32,41,42^ encode the unperturbed baseline as a single undivided space, allowing mitotic oscillations to fragment developmental trajectories.

Second, relative to a cell’s baseline state, perturbations act through two distinct biological modes that current frameworks fail to disentangle. A perturbation can act as a state-driving intervention that accelerates, arrests, or redirects cellular occupancy along a baseline trajectory, or as a state-driven intervention whose transcriptional effect is conditioned on a cell’s local baseline state. Population-level differential expression (DE) models^43–45^ and replicate-level pseudobulk workflows^46,47^ average across heterogeneous cells, conflating compositional shifts in state occupancy with state-dependent transcriptional response. Conversely, when tracking compositional changes, trajectory estimators^48–52^ and graph-based differential abundance methods^53–55^ construct cell-similarity graphs on the total embeddings, naturally conflating genuine abundance shifts with perturbation-induced topological distortions. Deep generative perturbation models^31,33,34,56,57^, on the other hand, commonly lack a perturbation-agnostic baseline reference, inevitably entangling intervention-driven abundance shifts with state-driven transcriptional responses.

Third, genetic or chemical perturbations that disrupt essential survival, metabolic, or homeo-static pathways frequently trigger acute cellular stress and cytotoxicity that drive transcriptomes off the physiological baseline manifold. Rather than modulating homeostatic lineage or activation programs, such interventions activate conserved endoplasmic reticulum stress and the unfolded protein response^2^, proteotoxic or oxidative stress pathways^6,58^, or apoptotic and viability-linked cytotoxic programs^12,58^. Separating these non-physiological stress responses from homeostatic state modulation is critical for distinguishing selective biological regulators and therapeutic adjuvants from toxic interventions that merely arrest growth or mimic quiescence. However, existing compositional and additive latent-shift models^33^, contrastive and multi-resolution salient-space architectures^32,41,42^, and optimal-transport frameworks^29,59^ operate under a closed-world manifold assumption that enforces no geometric boundary between on-manifold physiological transitions and off-manifold toxicity. Absorbing both homeostatic transitions and acute cellular stress into unconstrained perturbation embeddings, current methods can hardly distinguish whether an intervention pushes the cells along permitted, on-mani-fold trajectory or induce aberrant, off-manifold toxicity.

To address these challenges as a proof of principle, we introduce Disentangled Variational Inference for Perturbation Response (PertVI), a structurally disentangled counterfactual generative framework that resolves variance entanglement by replacing monolithic latent compression with biology-informed, separated manifolds. PertVI uses three perturbation-blind encoders to factorize each cell’s transcriptome into a low-dimensional circular cell-cycle anchor^60^, a macroscopic cellular state anchor, and an origin-anchored out-of-distribution anomaly sub-space, while an isolated linear decoder head absorbs donor and batch variation. Structural separation is enforced by decoding the cell-cycle subspace through a hard gene-marker mask^18^, penalizing cell-cycle weights in the main nonlinear decoder, and applying minibatch cross-covariance repulsion across subspaces. Perturbation identity enters exclusively after inference through a Split Feature-wise Linear Modulation module^61^ that applies perturbation-specific affine transformations to the cell-cycle and state anchors, whereas non-linear off-manifold pathology bypasses this transition and routes into the anomaly subspace. This architecture supports both unbiased differential abundance analysis on the perturbation-blind baseline manifolds and single-cell counterfactual differential expression across modulated states. Together, these components provide a unique multi-layered representation for single-cell perturbation data while delivering state-of-the-art performance in sample integration specifically designed for perturbation experiments. On the software side, PertVI leverages the optimization procedures and probabilistic modeling ecosystem of scvi-tools^62^, allowing it to scale to large-scale perturbation screens.

In this work, we first validate PertVI on simulated multi-covariate benchmarks with known ground truth, demonstrating that it simultaneously harmonizes batches, reconstructs circular cell-cycle topology and continuous differentiation trajectories, and recovers weak and targetoverlapping perturbation effects where existing methods fail. We then apply PertVI to two primary human T-cell screens in which all three entanglement regimes converge. In a chemical screen of differentiating CD8+ T cells^63^, PertVI distinguishes T-cell-sparing oncology drugs from conserved cell-cycle blockers, uncovers stage-restricted differentiation catalysts whose fate shifts are decoupled from global transcriptional magnitude, and separates genuine memory-precursor adjuvants from toxic phenocopies. In a multi-condition CRISPR interference screen of CD4+ T cells across resting and stimulated states^15^, PertVI resolves unsigned perturbation clusters into signed kinetic trajectories, uncovering tonic checkpoint brakes and biphasic homeostatic regulators that delay early activation but rebound into late effector hyper-activation, while preventing proliferative arrest from confounding T-cell activation phenotypes.

## RESULTS

### Disentangled variational inference for perturbation response

Conventional deep generative integration approaches and dimensionality reduction techniques usually compress whole-transcriptomic variance into a single, monolithic latent space. In highly proliferative systems like cancer cells and immune cells, however, this monolithic compression encounters a fundamental analytical bottleneck: high-amplitude transcriptional oscillations driven by the cell cycle routinely dominate the latent embedding, obscuring other subtle intrinsic cellular processes like metabolism and differentiation. Furthermore, in perturbation genomics, an entangled baseline mathematically conflates expression changes governed by intrinsic cellular state shift with state-driving perturbation momentum, making it difficult to determine whether a candidate perturbagen actively accelerates a permitted developmental transition or induces an aberrant stress response (**Fig. 1a**).

**Fig. 1.**
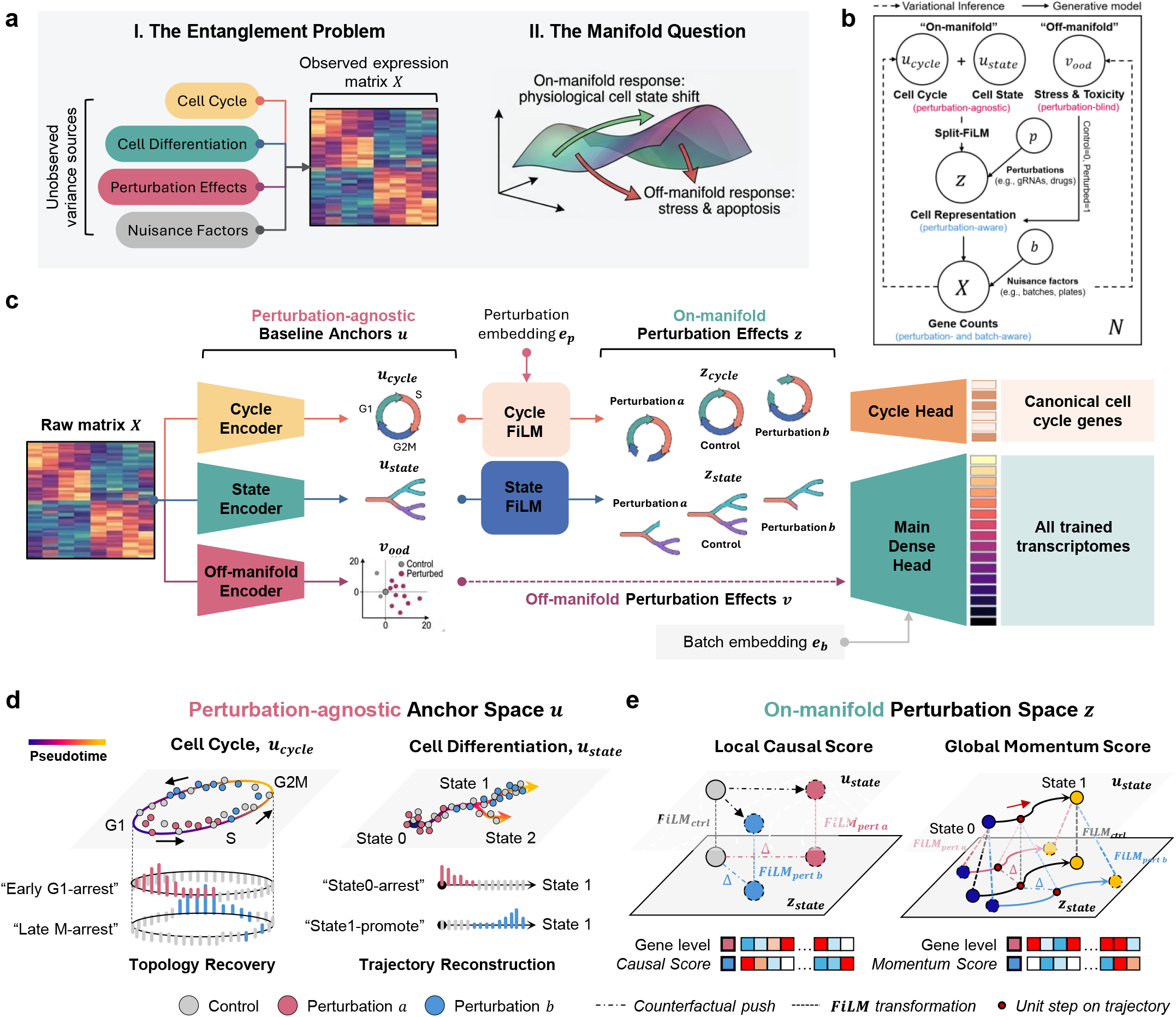
Architecture overview of PertVI. **a**, Conceptual motivation illustrating the variance entanglement problem and on-/off-manifold conflation in dynamic cellular systems. **b**, Probabilistic graphical model of PertVI comparing variational inference (dashed arrows) and the generative process (solid arrows). Latent variation is partitioned into on-manifold perturbation-agnostic baseline anchors (*u*_cycle_ for cell cycle and *u*_state_ for cell state) and an off-manifold perturbation-blind anomaly space (*v*_ood_ for stress and toxicity). Observed perturbations *p* modulate the baseline anchors via a Split-FiLM transition module to yield the perturbation-aware cell representation *z*, which combines with *v*_ood_ and technical nuisance factors *b* to generate the observed gene counts *X*. **c**, End-to-end neural network architecture of PertVI. From the input raw count matrix *X*, three parallel variational encoders operating without access to perturbation or batch labels (Cycle Encoder, State Encoder, and Off-manifold Encoder) infer the perturbation-agnostic baseline anchors *u*: a circular cell-cycle ring *u*_cycle_, a continuous cellular state trajectory *u*_state_, and an origin-anchored anomaly space *v*_ood_ (where control cells cluster at the origin (0, 0) and perturbed cells disperse outward). A learned perturbation embedding *e*_*p*_ feeds into two parallel Feature-wise Linear Modulation blocks (Cycle FiLM and State FiLM) that apply condition-specific affine scaling and shifting (Control, Perturbation a, Perturbation b) to *u*_cycle_ and *u*_state_ to form the on-mani-fold perturbation representations *z*_cycle_ and *z*_state_, whereas off-manifold perturbation effects *v*_ood_ bypass FiLM conditioning. In the bifurcated generative decoder, *z*_cycle_ and the batch embedding *e*_*b*_ decode through a masked linear Cycle Head restricted to canonical cell-cycle genes, while *z*_state_, *v*_ood_, and *e*_*b*_ decode through a nonlinear Main Dense Head to reconstruct all trained transcriptomes. **d**, Downstream analysis in the perturbation-agnostic anchor space *u* across Control (grey), Perturbation a (red), and Perturbation b (blue) populations. Left (Cell Cycle, *u*_cycle_): topology recovery along continuous circular pseudotime (G1 → S → G2M), where unrolled one-dimensional phase-density profiles resolve phase-specific occupancy shifts such as Early G1-arrest (Perturbation a) and Late M-arrest (Perturbation b). Right (Cell Differentiation, *u*_state_): trajectory reconstruction along ordered developmental states (State 0 → State 1 → State 2), resolving compositional shifts such as State0-arrest (Perturbation a) and State1-promote (Perturbation b). **e**, Counterfactual differential expression analysis in the on-manifold perturbation space *z*. Left (Local Causal Score): at a fixed local anchor on *u*_state_, condition-specific FiLM transformations (FiLM_ctrl_, FiLM_pert *a*_, and FiLM_pert *b*_) project identical baseline cells onto *z*_state_ to quantify the local counterfactual push Δ(dashed arrows), which decodes into a state-specific gene-level Causal Score vector. Right (Global Momentum Score): evaluating and integrating the counterfactual push Δacross discretized unit steps along the State 0 → State 1 trajectory on *u*_state_ and decoding to gene space yields the trajectory-integrated gene-level Momentum Score vector for each perturbation.

PertVI addresses this dual challenge by enforcing two levels of mathematical disentanglement (**Fig. 1b and c**). First, at the covariate level, PertVI learns a universal, perturbation-agnostic baseline (*u*_*base*_) that represents where a cell resides on the natural biological landscape, serving as baseline anchors. Within this baseline, a semantic filtration system physically quarantines high-amplitude periodic cell-cycle variance (*u*_*cycle*_) away from the macroscopic cellular state backbone (*u*_*state*_). Second, at the causal perturbation level, PertVI utilizes Feature-wise Linear Modulation (FiLM)^61^ to restrict “on-manifold” perturbation effect to an affine transcriptomic momentum (*z*_*cycle*_, *z*_*state*_) conditioned on the baseline anchor (*u*_*cycle*_, *u*_*state*_) (see **Methods**). Because FiLM is mathematically constrained to linear scaling and shifting along permitted biological axes, it cannot accommodate topological ruptures; consequently, PertVI routes any residual, non-linear perturbation variance, such as synthetic stress and toxicity, into a dedicated out-of-distribution novelty space (*v*_*ood*_) that the natural baseline anchor cannot explain.

By decoupling the perturbation-agnostic baseline manifold *u*_*base*_ from the perturbation-aware affine shift *z*_*base*_, a trained PertVI model enables two complementary analytical regimes that overcome the current limitations of conventional single-cell perturbation pipelines (**Fig. 1d and e**).

In the first regime (**Fig. 1d**), the baseline manifolds (*u*_*cycle*_ and *u*_*state*_) serve as an unbiased coordinate canvas for investigating cell-cycle progression, constructing developmental trajectories, and performing Differential Abundance (DA) analysis. Conventional differential abundance methods rely on clustering or integrating treated and control cells into a shared latent space, which is prone to algorithmic over-correction that erases genuine treatment-induced biological shifts, or under-correction that leaves residual batch artifacts. Because PertVI learns a universal baseline manifold using probabilistic encoders that operate strictly blind to perturbation labels during inference, mapping treated and control populations onto *u*_*cycle*_ and *u*_*state*_ allows for the rigorous quantification of shifts in phenotypic occupancy, such as drug-induced lineage selection or cell-cycle arrest, without confounding from supervised label leakage or integration artifacts.

In the second regime (**Fig. 1e**), the perturbation-aware latent spaces *z*_*cycle*_ and *z*_*state*_ enable two kinds of counterfactual Differential Expression (cDE) analyses: state-specific responses and state-shifting momentum. In standard observational DE analyses, comparing treated and control populations inevitably conflates perturbation-induced expression changes with baseline covariate imbalances and perturbation-induced population shifts. PertVI resolves this confounding by holding an identical basal coordinate anchor (*u*_*base*_) constant while substituting alternative treatment conditions into the FiLM engine. By generating counterfactual trajectories, PertVI allows the investigation on whether a perturbagen induces static, state-specific transcriptional signatures within an isolated sub-population or exerts active, state-shifting perturbation momentum that propels cells along permitted Waddington trajectories.

A comprehensive description of mathematical formulation of the PertVI architecture, objective function governance, and the post-training counterfactual analytical framework is provided in the **Methods**.

### Disentangling perturbation response from cellular baseline on a synthetic dataset

To validate the structural disentanglement capabilities of PertVI, we benchmarked the architecture against conventional analytical tools (Linear Discriminant Analysis, LDA; COMBAT^64^) and the deep generative models for data integration and variance disentanglement (LinearSCVI^35^, MrVI^41^, DrVI^37^, contrastiveVI^32^, and MoCAVI^42^) using a synthetic simulation with ground-truth sources of variation (**Fig. 2a, b; Extended Data Fig. 1**; See **Methods**). The synthetic dataset comprised 10,000 cells and 100 genes encoding four concurrent sources of transcriptomic variance in a hierarchical structure, quantified as mean per-gene variance fractions on log-normalized counts: cell differentiation state (19.6%), cell-cycle progression (10.9%), sequencing batch (4.6%), and transcriptional perturbation (3.6%). Cell-cycle expression was modeled as a continuous half-sinusoidal oscillation across three phases (G1-S-G2M), with phase-specific gene expression profiles designed to mimic the ring topology of the cell cycle (**Extended Data Fig. 1a-b**). Differentiation state was encoded as a continuous trajectory through three ordered states (State1-State2-State3) using tent-function basis functions, with a peak log-foldchange (LFC) of 5.0 relative to a cell-cycle peak of 3.5 (**Extended Data Fig. 1c and d**). Four perturbations were constructed with a defined similarity structure (**Fig. 2b; Extended Data Fig. 1e and f**): Pert-A and Pert-B were strong perturbations each targeting 50 genes, with 90% gene-level overlap between them; Pert-C was a weak perturbation targeting 18 genes drawn exclusively from loci not targeted by Pert-A or Pert-B; Pert-D introduced negligible genome-wide noise as a near-null control. Counts were sampled from a Zero-Inflated Negative Binomial model, yielding a dataset with 41.2% sparsity and a mean of 6,263 UMIs per cell (**Extended Data Fig. 1g**). In addition, a bidirectional cross-covariate coupling term between state and cell-cycle marker genes was included to reflect the partial non-orthogonality inherent to biological data (**Extended Data Fig. 1h and i**). In this design, perturbation is the weakest and most confounded variance component, representing a benchmark condition in which monolithic latent representations are expected to fail. To benchmark, all model-based approaches were trained on identical train/test splits with matched batch sizes and learning rates; methods requiring hyperparameter optimization underwent grid search prior to evaluation. A detailed description of the model training, the parameter settings, and the performance is provided in the **Methods** and supplementary materials (**Extended Data Fig. 2a**).

**Fig. 2.**
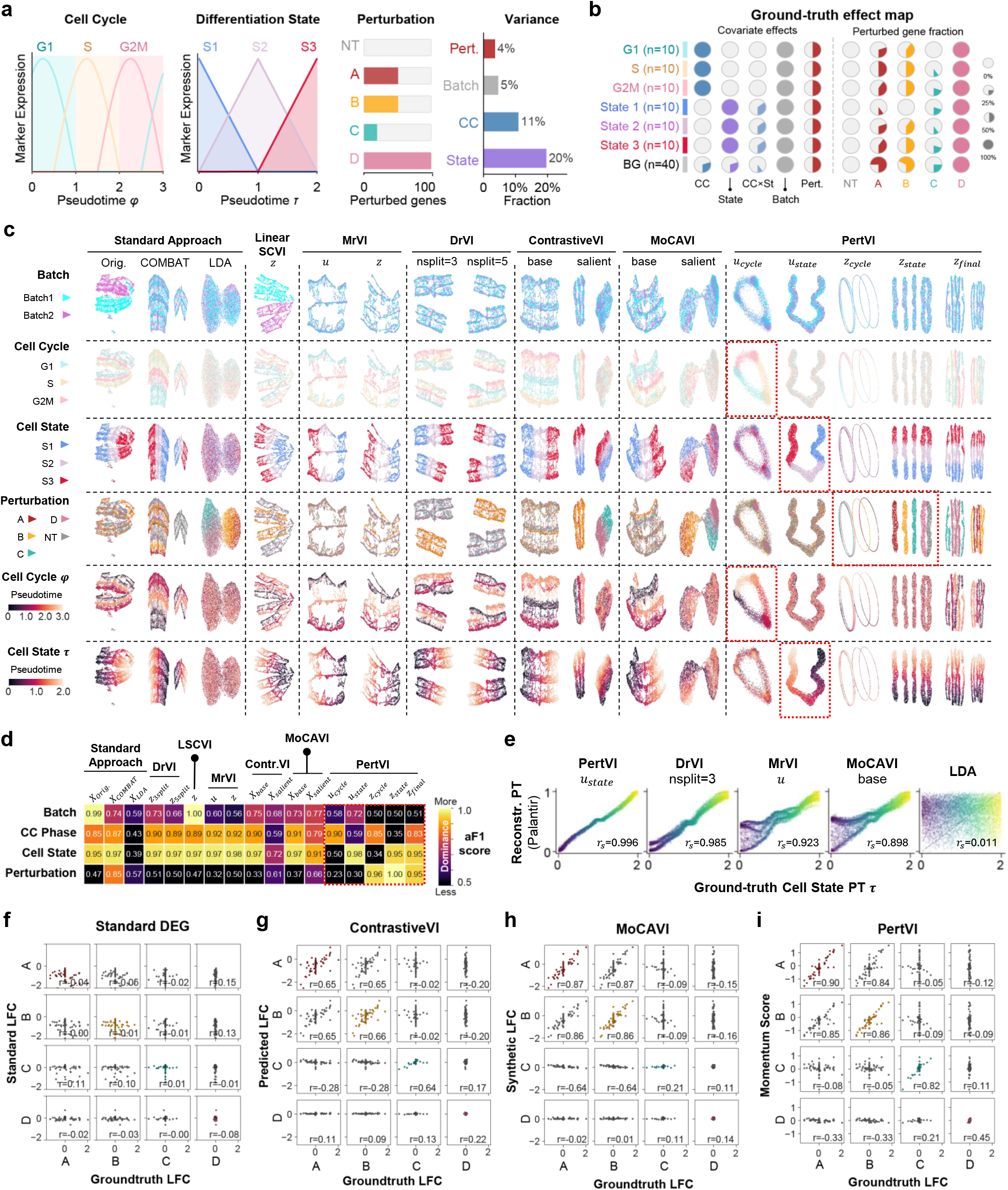
Simulation benchmark of PertVI against existing disentanglement models. **a**, Schematic of the synthetic multi-covariate Perturb-seq dataset (*N* = 10,000 cells across *G* = 100 genes). Left: cell-cycle marker expression curves along circular pseudotime *φ* ∈ [0, 3) across G1 (*φ* ∈ [0, 1), peak at 0.25), S (*φ* ∈ [1, 2), peak at 1.25), and G2M (*φ* ∈ [2, 3), peak at 2.25). Middle left: differentiation state marker expression curves along continuous pseudotime *τ* ∈ [0, 2] across S1 (State1, *τ* ∈ [0, 2/3)), S2 (State2, *τ* ∈ [2,/34/3)), and S3 (State3, *τ* ∈ [4,/32]). Middle right: perturbation target gene counts across the *G* = 100genes (*n* = 2,000 cells per condition) for non-targeting control NT (0 perturbed genes), strong co-functional perturbations A (Pert-A, 50 genes) and B (Pert-B, 50 genes sharing 45 targets [90% overlap] with A), weak orthogonal perturbation C (Pert-C, 18 non-overlapping genes), and near-null negative control D (Pert-D, diffuse sub-threshold noise *σ* = 0.05 across all 100genes). Right: bar chart of mean per-gene one-way ANOVA variance fractions (*η*^2^ on ln(CP10K + 1)-normalized counts) showing the four-tier hier-archy: State (20%; exact 19.6%) >CC (11%; exact 10.9%) >Batch (5%; exact 4.6%) >Pert. (4%; exact 3.6%). **b**, Ground-truth effect map across the seven simulated gene groups: G1 (*n* = 10), S (*n* = 10), G2M (*n* = 10), State 1 (*n* = 10), State 2 (*n* = 10), State 3 (*n* = 10), and background BG (*n* = 40). Left (Covariate effects): pie symbols (0%, 25%, 50%, 100%) indicate direct and cross-coupled variance contributions from cell cycle (CC), differentiation state (State), bidirectional cell-cycle and state coupling (CC+St), technical batch (Batch), and perturbation (Pert.). Right (Perturbed gene fraction): pie symbols indicate the fraction of genes targeted within each gene group across conditions NT, A, B, C, and D. **c**, two-dimensional UMAP embeddings (*N* = 10,000 cells per panel) comparing 17 latent representations across Standard Approach (uncorrected Orig., COMBAT, and LDA), LinearSCVI (*z*), MrVI (*u* and *z*), DrVI (nsplit = 3 and nsplit = 5), ContrastiveVI (base and salient), MoCAVI (base and salient), and PertVI (*u*_cycle_, *u*_state_, *z*_cycle_, *z*_state_, and *z*_final_). Rows color cells by sequencing Batch, discrete Cell Cycle phase, discrete Cell State, Perturbation condition, continuous Cell Cycle *φ* Pseudotime, and continuous Cell State *τ* Pseudotime. Red dashed boxes mark PertVI’s selective recovery of the closed circular cell-cycle ring in *u*_cycle_, the ordered differentiation trajectory in *u*_state_, and perturbation separation in *z*_cycle_, *z*_state_, and *z*_final_. **d**, heatmap of 5-fold stratified cross-validation *k*-nearest-neighbor (*k* = 10) macro-averaged adversarial F1 across the 17 representations. Scores are evaluated across Batch, CC Phase, Cell State, and Perturbation. **e**, Scatter plots comparing reconstructed Palantir differentiation pseudotime (normalized onto [0, 1], *y*-axis) against groundtruth Cell State pseudotime (*τ* ∈ [0, 2], *x*-axis, colored by *τ*) across all *N* = 10,000 cells for PertVI *u*_state_, DrVI nsplit = 3, MrVI *u*, MoCAVI base, and LDA. **f–i**, Scatter-plot matrices across all *G* = 100 genes comparing inferred gene-level perturbation effect vectors (*y*-axis, rows A, B, C, D) against ground-truth perturbation log-fold changes (*x*-axis, columns A, B, C, D; dashed lines mark *x* = 0and *y* = 0). **f**, Standard DEG (empirical LFC from two-sided Wilcoxon rank-sum testing vs. NT). **g**, ContrastiveVI (predicted LFC from salient posterior decoding). **h**, MoCAVI (synthetic LFC from synthetic differential abundance). **i**, PertVI (path-integrated Momentum Score).

We first assessed whether each method could recover the baseline biological manifold from the four concurrent variance sources (**Fig. 2c**). Disentanglement performance was quantified using adversarial F1 (aF1) and LISI scores computed independently for each of the four ground-truth variance sources (**Fig. 2d; Extended Data Fig. 2b**; see **Methods**). Batch integration was assessed first as a prerequisite filter: all methods except LinearSCVI achieved adequate batch mixing (aF1 < 0.80; LISI > 0.20), compared to the uncorrected embedding (aF1 = 0.99; LISI = 0.01); LinearSCVI was therefore excluded from further disentanglement evaluation. Among the remaining methods, only PertVI resolved the cell-cycle and cell-state manifolds into separate, interpretable subspaces (**Fig. 2e; Extended Fig. 2c**). The perturbationagnostic cell-cycle encoder (*u*_*cycle*_), constrained to a low-dimensional topology with a relaxed prior, reconstructed the continuous ring structure of the cell cycle, with G1, S, and G2M populations forming smooth, overlapping arcs fully aligned with the ground-truth cell-cycle pseudotime gradient (**Fig. 2c**, “Cell Cycle” and “Cell Cycle *φ*”, PertVI *u*_*cycle*_; aF1 = 0.90, LISI = 0.04 for the cell-cycle variable in *u*_*cycle*_). The perturbation-agnostic macroscopic state encoder (*u*_*state*_), operating without any prior knowledge of differentiation state, autonomously resolved the three ground-truth states as an ordered trajectory (aF1 = 0.98, LISI = 0.00 for the cell-state variable in *u*_*state*_), with the cell-cycle pseudotime gradient absent from the state axis (**Fig. 2c**, “Cell State” and “Cell State *τ*”, PertVI *u*_*state*_; aF1 = 0.59, LISI = 0.36 for the cellcycle variable in *u*_*state*_). No competing method achieved this dual resolution.

COMBAT, applied with perturbation as covariate, left cell-cycle and cell-state signals entangled within a single corrected space (**Fig. 2c and d**, “Cell Cycle” and “Cell State”, COMBAT). LDA (Linear Discriminant Analysis) separated perturbation clusters but conflated cell-cycle and cell-state variance within the same projection, with perturbation effects dominating the embedding (**Fig. 2c and d**, “Cell Cycle” and “Cell State”, LDA). MrVI’s sample-unaware *u* space removed perturbation-driven variance from the biological background but could not partition cell-cycle from cell-state variance within a single continuous representation; its sampleaware *z* space further failed to isolate perturbation from the cellular background (**Fig. 2c and d**, “Cell Cycle” and “Cell State”, MrVI *u* and *z*). Both contrastive learning approaches, ContrastiveVI and MoCAVI, partitioned perturbation-induced variance into a salient space and biology into a background space (**Fig. 2c and d**, ContrastiveVI and MoCAVI); however, the background space itself remained a joint, undifferentiated mixture of cell-cycle and cell-state variance with no further factorization. The unsupervised approach DrVI, on the other hand, distributed cell-state variance across all factored subspaces, such that no individual split represented a unique variance source (**Fig. 2c and d**, DrVI with n_split=3 and n_split=5; **Extended Data Fig. 2d and 2e**). Importantly, visualizing PertVI’s full concatenated representation (*z*_*final*_) produced the same entangled topology observed in all competing methods (**Fig. 2c**, PertVI *z*_*final*_, compared to the salient spaces of ContrastiveVI and MoCAVI and the total latent space of DrVI), confirming that signal conflation is an intrinsic consequence of joint compression and that upstream semantic filtration is necessary to recover interpretable biological subspaces.

We next evaluated perturbation recovery across two complementary levels: the separation of perturbation conditions in latent embedding space, and the accuracy of gene-level perturbation effect estimation against the simulation ground truth (see **Methods**). At the embedding level, five methods produced representations in which the four perturbation conditions separated from the non-targeting control: LDA, ContrastiveVI (salient space), MoCAVI (salient space), DrVI (total *z*), and PertVI (perturbation-aware *z*_*cycle*_, *z*_*state*_, *z*_*final*_) (**Fig. 2c and d**). Among these, only PertVI’s perturbation-aware spaces achieved aF1 scores above 0.90 and LISI scores near zero for the perturbation variable (*z*_*cycle*_: aF1 = 0.96, LISI = 0.04; *z*_*state*_: aF1 = 1.00, LISI = 0.00; *z*_*final*_: aF1 = 0.95, LISI = 0.09), indicating a substantially higher signal-to-noise separation than any competing representation (**Fig. 2d; Extended Data Fig. 2b)**. At the gene-level, we compared the inferred perturbation effects from methods with a generative or analytical decoder capable of producing gene-level output: LinearSCVI, ContrastiveVI, MoCAVI, and PertVI (see **Methods**). Although LinearSCVI failed batch correction and was excluded from the embedding-level comparison, it provides gene-level perturbation effect estimates and was therefore included here for completeness. Overall, each method’s inferred gene-level effects were compared to the known LFCs for each of the four perturbations using Pearson correlation, alongside a standard differential expression (DE) baseline (**Fig. 2f-i**). Among them, the standard DE analysis failed to recover ground-truth LFC for all four perturbations (**Fig. 2f**).

LinearSCVI recovered weak signal for the two strong perturbations (r = 0.31, Pert-A; r = 0.28, Pert-B) but showed negligible correlation for the weak and near-null perturbations (r = 0.11, Pert-C; r = 0.21, Pert-D) (**Extended Data Fig.2f**). ContrastiveVI, on the other hand, produced more uniform recovery across perturbation strengths (r = 0.65, 0.66, 0.64, 0.22 for Pert-A, -B, -C, -D) (**Fig.2g**), while MoCAVI achieved high correlations for the two strong perturbations but failed on the weak and near-null ones (r = 0.87, 0.86, 0.21, 0.14 for Pert-A, -B, -C, -D) (**Fig.2h**). In contrast, PertVI’s perturbation momentum achieved the highest correlation across all four perturbations (r = 0.90, 0.86, 0.82, 0.45 for Pert-A, -B, -C, -D) and was the only method to recover meaningful signal for Pert-C, the weak perturbation targeting only 18 genes (**Fig.2i**). This advantage likely stems from PertVI’s Split-FiLM mechanism on the disentangled biological axes: because a subset of perturbation target genes is designed to overlap with cell-cycle and cell-state marker genes (**Fig. 2b**) (a common scenario in real perturbation studies, where drug targets frequently participate in cell-cycle regulation or differentiation programs), modeling perturbation effects independently along each biological axis conferred sensitivity to weak signals that are otherwise masked by the dominant cellular variance. Of note, across all methods that recovered perturbation signal, the within-benchmark similarity structure of the four perturbations (Pert-A ≈ Pert-B, Pert-C orthogonal, Pert-D near-null) was recapitulated in all the respective inferred gene-level perturbation effects (**Extended Data Fig. 2g**). Taken together, both MoCAVI and ContrastiveVI represent capable approaches for gene-level perturbation effect inference under strong perturbation regimes; PertVI extends this capacity to weak perturbations and the scenario where target gene overlaps with endogenous biological programs require axis-separated modeling to achieve reliable signal recovery.

Aggregating across all benchmarked criteria, i.e., batch correction, cell-cycle topology reconstruction, differentiation state recovery, and perturbation signal recovery, PertVI was the only method to achieve adequate performance on all four axes simultaneously. Each competing method was limited by architectural constraints that produced a specific failure mode: LinearSCVI failed batch correction; COMBAT and LDA could not factorize cell-cycle from cell-state variance; MrVI, ContrastiveVI, and MoCAVI lacked dedicated encoders for the two baseline biological processes; DrVI distributed dominant variance sources across all split spaces without specificity; and no method other than PertVI recovered the weak perturbation signal against the confounded cellular background. These failure modes collectively reflect a fundamental limitation of monolithic and undifferentiated latent representations when applied to data in which multiple biological processes partially share gene targets and span a range of effect sizes.

### PertVI reveals heterogenous regulation by anti-cancer agents on CD8+ effector T cell differentiation

Next, we applied PertVI to a high-throughput *in vitro* pharmacological screen in primary human CD8+ T cells^63^ (**Fig. 3a**; see **Methods**). This dataset profiles 62,207 peripheral blood mononuclear cell-derived CD8+ T cells from two healthy blood donors (Donor 1: 34,548 cells; Donor 2: 27,659 cells) across 62 small-molecule compounds (predominantly anti-cancer agents; 55,735 treated cells) and dimethyl sulfoxide (DMSO) vehicle controls (6,472 cells). Naive CD8+ T cells were pre-incubated with each compound (1*µM*) across two biological replicates for 24 h, followed by 72 h of anti-CD3/CD28 co-stimulation and IL-2 treatment to induce clonal proliferation and effector differentiation across five developmental states: quiescent naive T cells (n = 16,060), early proliferating blasts (n = 18,796), intermediate memory precursors (n = 5,076), cytolytic effectors (n = 15,505), and terminal effectors (n = 6,770). Primary singlecell perturbation screens of this scale pose a fundamental analytical challenge: donor and replicate variation, periodic cell-cycle progression, continuous lineage differentiation, and pleiotropic drug responses are nonlinearly entangled within each cell’s transcriptome. To resolve these sources of variation within a unified causal framework (**Fig. 1**; see **Methods**), we assigned the biological replicate identifier as the technical batch covariate (*b*_*n*_) and compound identity as the target perturbation covariate (*p*_*n*_). PertVI strips technical batch artifacts during initial encoding to infer two perturbation-agnostic single-cell baseline anchors: a 2D circular cell-cycle manifold (*u*_*cycle*_) and a macroscopic developmental state manifold (*u*_*state*_). On-manifold pharmacological effects are modeled via a Split-FiLM transition that applies condition-specific affine shifts independently to each baseline anchor (*z*_*cycle*_ and *z*_*state*_). Conversely, non-homeostatic cellular stress and toxicity that depart from physiological trajectories are routed into an orthogonal out-of-distribution anomaly subspace (*v*_*ood*_) before decoding through a bifurcated gene-expression architecture. Across 600 training and validation epochs, PertVI converged smoothly without overfitting in evidence lower bound (ELBO), reconstruction loss, and total Kullback–Leibler (KL) divergence (**Extended Data Fig. 33**).

**Fig. 3.**
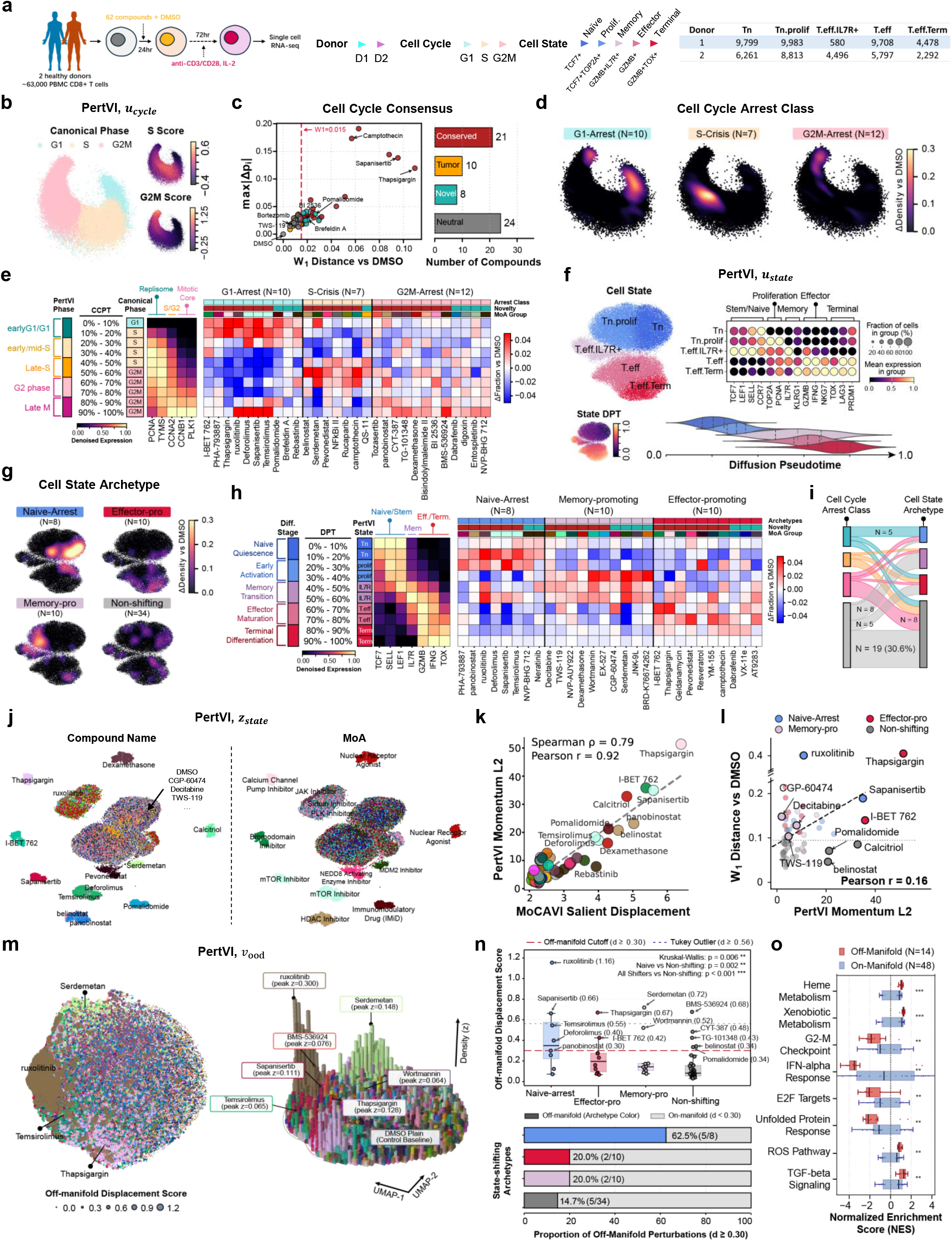
PertVI reveals pharmacological interaction between anti-cancer drugs and CD8+ T cell effector differentiation. **a**, Experimental schematic of the primary human CD8+ T-cell pharmacological screen (GSE306429). **b**, UMAP embeddings of PertVI’s circular cell-cycle baseline manifold (*u*_cycle_) colored by canonical discrete cell-cycle phase (G1, S, G2M), continuous S Score, and continuous G2M Score. **c**, Compound-level Cell Cycle Consensus scatter plot (1D Wasserstein Distance vs. DMSO *W*_1_ against Max Absolute Bin Shift max ∣ Δ*p*_*i*_ ∣, empirical replicate threshold *W*_1_ ≥ 0.015) and top summary bar chart partitioning all 63 conditions (62 compounds plus DMSO) into four immuno-oncology classes: Conserved (*n* = 21), Tumor-activeonly / T-cell-sparing (*n* = 10), Novel in T cells (*n* = 8), and Neutral (*n* = 24). **d**, Three 2D kernel density estimation (KDE) differential density contour maps (ΔDensity vs. DMSO on *u*_cycle_) for the 29 active cellcycle modulators (21 Conserved plus 8 Novel): G1-Arrest (*N* = 10), S-Crisis (*N* = 7), and G2M-Arrest (*N* = 12). **e**, Ten-bin continuous cell-cycle progression (CCPT, deciles 0% to 10% through 90% to 100%) differential abundance heatmap (ΔFraction vs. DMSO) across the 29 active compounds, aligned with PertVI’s five continuous sub-phases, canonical phases, and five denoised marker waves. **f**, UMAP embeddings of PertVI’s developmental state baseline manifold (*u*_state_) colored by the five CD8+ T-cell differentiation states and continuous Diffusion Pseudotime, alongside the 14-marker expression dotplot. **g**, Four KDE differential density contour maps (ΔDensity vs. DMSO on *u*_state_) for the four differentiation archetypes: C1 Naive-Arrest (*N* = 8), C2 Effector-promoting (*N* = 10), C3 Memory-shifting (*N* = 10), and Non-shifting (*N* = 34). **h**, Ten-bin DPT differential abundance heatmap across the 28 active state-shifting compounds, aligned with developmental stage and seven denoised marker trajectories. **i**, Cross-manifold alluvial ribbon diagram linking Cell Cycle Arrest Class (*u*_cycle_) to Cell State Archetype (*u*_state_) across all 62 compounds, highlighting significant coupling between G1 arrest and C1 Naive-arrest (5/10 compounds, standardized residual *z* = +3.27, *p* < 0.001) and cell-cycle sparing among C3 Memory-shifting compounds (8/10 compounds, *z* = +1.16). **j**, UMAP embeddings of PertVI’s perturbation-aware state space (*z*_state_) colored by Compound Name (left) and MoA (right), illustrating co-clustering of shared-MoA drugs while low-Momentum catalysts localize near DMSO. **k**, Scatter plot of MoCAVI Salient Displacement versus PertVI Momentum L2 across all 62 compounds. **l**, Broken y-axis scatter plot of phenotypic state shift (1D Wasserstein Distance vs. DMSO on *u*_state_) against PertVI Momentum L2 across all 62 compounds. **m**, UMAP embedding of PertVI’s 5-dimensional anomaly subspace (*v*_ood_) colored by Euclidean off-manifold displacement score *d*_ood_(left) and 3D continuous KDE density landscape spires. **n**, Boxplot of off-manifold displacement *d*_ood_ across the four differentiation archetypes and bar chart of the proportion of off-manifold compounds. **o**, Preranked MSigDB Hallmark normalized enrichment score (NES) boxplots derived from the state-invariant Marginalized Counterfactual Operator (MCO) comparing Off-Manifold (*N* = 14, red) versus On-Manifold (*N* = 48, blue) compounds across eight differentially regulated pathways.

#### Benchmarking baseline disentanglement

Across all latent subspaces, PertVI harmonized donor and replicate variation so that the perturbation-agnostic baseline anchors (*u*_*cycle*_, *u*_*state*_) and perturbation-aware spaces (*z*_*cycle*_, *z*_*state*_, *z*_*final*_) reflected biological and pharmacological variation rather than technical batch structure (**Extended Data Fig. 33, c**, “Donor”; adversarial F1 (*aF*1) =0.76 and Local Inverse Simpson’s Index (LISI) =0.17 in *z*_*final*_, with aF1=0.53 to 0.72 in *u*_*cycle*_, *z*_*cycle*_, and *z*_*state*_). In actively differentiating lymphocytes, periodic cell-cycle signals strongly covary with lineage progression. PertVI resolves this entanglement by routing proliferative variation through its dedicated circular encoder, reconstructing the continuous topology of the cell cycle on *u*_*cycle*_ (cell-cycle phase aF1=0.87, LISI=0.00; cell state aF1=0.32, LISI=0.44; **Extended Data Fig. 33, c**). Quarantining proliferative variance within *u*_*cycle*_ freed the macroscopic state encoder to recover an unconfounded CD8+ T-cell differentiation trajectory on *u*_*state*_ (cell state aF1=0.97, LISI=0.02; cell-cycle phase aF1=0.53, LISI=0.29; **Extended Data Fig. 33, c**). By contrast, linear empirical Bayes batch regression in COMBAT overcorrected for covariate variation and shattered the developmental continuum, whereas baseline embeddings from deep generative models including MoCAVI and MrVI remained dominated by cell-cycle variation (**Extended Data Fig. 33, c**). Consistent with our simulation benchmarks (**Fig. 2c**), PertVI’s concatenated representation (*z*_*final*_) reproduced the similar entangled topology of MoCAVI *base* and MrVI *u* (**Extended Data Fig. 33**), confirming that upstream semantic filtration is required to recover unconfounded developmental trajectories.

PertVI’s Split-FiLM transition also overcame a major failure mode of multi-condition perturbation modeling when all 62 drugs were supplied as target covariates (**Extended Data Fig. 33**, “Perturbation”). Whereas supervised LDA and COMBAT conflated compound responses with baseline state identity, MrVI failed to resolve individual compound effects except Thapsigargin (perturbation aF1=0.04, LISI=1.09), likely because its high-magnitude response monopolized the variance of shared sample-aware latent space. MoCAVI, on the other hand, effectively captures dominant perturbation effects upon centroid aggregation (perturbation aF1=0.21, LISI=0.93); however, it suffers from a critical control-memorization artifact in imbalanced screens. Compared to all the benchmarked approaches, PertVI successfully isolated individual pharmacological effects in *z*_*cycle*_ (perturbation aF1=0.45, LISI=0.54) and *z*_*state*_ (perturbation aF1=0.53, LISI=0.67) while keeping the baseline manifolds free of drug-specific clustering and control-memorization artifacts (**Extended Data Fig. 33, c**). We hence used this disentangled causal structure to address three central biological questions across the screen: (1) how anti-cancer agents differentially remodel primary T-cell proliferation versus tumor division on *u*_*cycle*_; (2) how compounds redirect physiological CD8+ T-cell differentiation along *u*_*state*_ and what transcriptional principles govern these transitions in *z*_*state*_; and (3) which compounds induce aberrant, off-manifold cellular toxicity in *v*_*ood*_.

#### Continuous cell-cycle reconstruction and lineage-selective proliferation remodeling on u_cycle_

On the 3D circular manifold *u*_*cycle*_, continuous angular pseudotime (*t* ∈ [0, 1]) smoothly ordered canonical G1, S, and G2/M phase scores across ten progression deciles (**Fig. 3b; Extended Data Fig. 33**). To compare pharmacological cell-cycle regulation in primary human CD8+ T lymphocytes against canonical arrest phenotypes in solid-tumor and leukemic cell lines (**Fig. 3c**), we implemented a two-stage unsupervised discovery pipeline that first filtered compounds against empirical DMSO replicate variation (1D Wasserstein distance *W*_1_ ≥ 0.015 versus maximum bin shift) and then clustered row-normalized sub-phase profiles, separating 29 active cell-cycle modulators from 34 T-cell-sparing conditions (33 compounds plus DMSO; **Fig. 3c**; see **Methods**). Cross-referencing these primary T-cell phenotypes against published tumor arrest profiles partitioned the 63 conditions into four immuno-oncology consensus classes (**Fig. 3c**). The Conserved class (*n* = 21) comprised lineage-invariant antiproliferative agents that arrested both tumor cells and primary CD8+ T cells, including the JAK1/2 inhibitor ruxolitinib, mTOR inhibitors Sapanisertib, Temsirolimus, and Deforolimus, the interphase CDK2/4/5 inhibitor PHA-793887^65^, the topoisomerase I inhibitor Camptothecin, MDM2-p53 modulator Serdemetan^66^, pan-HDAC inhibitors Panobinostat and Belinostat^67,68^. The Tumor-active-only/T-cell-sparing class (*n* = 10) arrested cancer cell lines but spared primary CD8+ T-cell division at 1*µM*. This T-cell-sparing class spanned the proteasome inhibitor bortezomib, HSP90 inhibitors geldanamycin and NVP-AUY922, MEK/ERK inhibitors selumetinib and VX-11e, the hypomethylating agent decitabine, the CDK1/2/9 inhibitor CGP-60474, the Aurora/JAK2 inhibitor AT9283, the class I HDAC inhibitor Apicidin, and solid-tumor receptor tyrosine kinase (RTK) inhibitors crizotinib, ponatinib, and sorafenib. Conversely, the Novel in T cells class (*n* = 8) uncovered *de novo* T-cell proliferation remodelers that are non-canonical in solid-tumor screens but altered lymphocyte division, including pomalidomide (early-G1 arrest via IKZF1/IKZF3 degradation^69^). The remaining conditions formed the universally cell-cycle-neutral class (Quadrant IV, *n* = 24, including DMSO, TWS-119, and resveratrol; **Fig. 3c**).

Across the 29 active cell-cycle modulators (21 Conserved plus 8 Novel), 2D kernel density estimation (*ΔDensity* versus DMSO on *u*_*cycle*_; **Fig. 3d**) and 10-bin circular progression profiling (**Fig. 3e**) resolved three arrest regimes aligned with five sequential sub-phases and five denoised marker waves (*PCNA, TYMS, CCNA2, CCNB1*, and *PLK1*): G1-arrest (*n* = 10, peaking in early-G1/G1 Bins 0 to 1, where ruxolitinib and mTOR inhibitors also exhibited secondary late-M cytokinesis retention), S-crisis (*n* = 7, peaking in early/mid-S Bins 2 to 3 and late-S Bins 4 to 5), and G2/M-arrest (*n* = 12, peaking in G2 Bins 6 to 7 and late-M Bins 8 to 9). Of note, the continuous angular coordinates on *u*_*cycle*_ recovered the Late-S Crisis Zone (*t* ∈ [0.45, 0.65]; **Extended Data Fig. 33–h**) that is commonly masked by conventional score-based phase calling. Because canonical S and G2/M gene-set scores crossed over prematurely at *t* ≈ 0.49 (**Extended Data Fig. 33**), denoised S-phase synthesis markers *TYMS* and *PCNA* peaked around Bin 5 (*t* ∈ [0.45, 0.65]) while *CCNA2* rose and mitotic regulators *CCNB1* and *PLK1* remained uninduced (**Extended Data Fig. 33**). Consequently, discrete phase scoring misclassified 87.4% of Bin 5 cells as G2/M and mislabeled the late-S replication crisis peaks (*t* ≈ 0.53, mitosis depleted ≤ 2.2) of camptothecin^70^ (60.7% canonical G2/M; Bin 5 *ΔFraction* = +0.173) and Serdemetan^71^ (54.7% canonical G2/M; Bin 5 *ΔFraction* = +0.191) as mitotic arrest (**Extended Data Fig. 33, g**). PertVI’s continuous sub-phase calling rescued both drugs as Late-S replication crisis (49.9% and 55.3% Late-S), cleanly distinguishing them from true mitotic blockers BI 2536^72^ (64.6 Early-G2/M) and Tozasertib^73^ (72.6% Late-M; **Extended Data Fig. 33**).

#### Developmental trajectory reconstruction and compound-induced phenotypic shift on u_state_

Having resolved how compounds remodel cell division on *u*_*cycle*_, we next examined how they redirect lineage differentiation along the unconfounded developmental manifold *u*_*state*_ (**Fig. 3f**; **Extended Data Fig. 44**). Across Diffusion Pseudotime (DPT, *τ* ∈ [0, 1]) deciles, denoised marker expression resolved five sequential CD8+ T-cell states (**Fig. 3f** and **Extended Data Fig. 44**): quiescent naive T cells (Tn, Bins 0 to 1; peak *TCF7, LEF1, SELL*, and *CCR7*), early proliferating blasts (Tn.prolif, Bins 2 to 3; peak *TOP2A* and *PCNA*), an intermediate memory-precursor window (T.eff.IL7R+, Bins 4 to 5; unimodal *IL7R* peak reaching 0.728 in Bin 4 and 1.000 in Bin 5), cytolytic effectors (T.eff, Bins 6 to 7; induction of *GZMB, IFNG, PRF1*, and *NKG7*), and terminal effectors (T.eff.Term, Bins 8 to 9; maximal *TOX, LAG3, GZMB, IFNG*, and *PRDM1*). To further characterize compound-induced phenotypic shift, we combined hierarchical clustering with dual-front statistical union gate, partitioning 62 compounds into 34 non-shifting compounds indistinguishable from DMSO and 28 active compounds showing three distinct differentiation regulation archetypes (**Fig. 3g, h; Extended Data Fig. 44, c**; see **Methods**). Specifically, archetype C1 (“Naïve arrest”, *n* = 8) blocked exit from naive and early proliferative states. This archetype includes compounds like mTOR inhibitors (Temsirolimus, Deforolimus^74^, and Sapanisertib^75^) and JAK1/2 inhibitor ruxolitinib^76^, consistent with the requirement for mTOR and JAK-STAT signaling in TCR-driven blastogenesis^77,78^. Conversely, archetype C2 (“Effector-promoting”, *n* = 10) depleted naïve states and accelerated effector accumulation. Compounds in this archetype include I-BET762, a Bromodomain and extra-terminal inhibitor that is reported to enhance cytotoxic T cell effector function^79^, and Thapsigargin, an endoplasmic reticulum calcium ATPase pump inhibitor^80^ that is known to promote T cell activation^81^. Finally, archetype C3 (“Memory-shifting”, *n* = 10) selectively expanded the intermediate *IL7R*+ memory-precursor pool (Deciles 4 to 5, *ΔPMF* = +5.2 to +7.1) while depleting terminal effectors (Deciles 7 to 9). Prominent C3 compounds include the DNA methyltransferase inhibitor Decitabine, which prevents *de novo* exhaustion methylation and enhances CAR-T persistence^82,83^; the GSK-3*β* inhibitor TWS-119, which activates Wnt-*β*-catenin/TCF-1 signaling to generate stem-like memory CD8+ T cells^84,85^; the glucocorticoid receptor agonist dexamethasone^86,87^, which may enhance the binding of glucocorticoid to the enhancer near IL-7Rα locus and promote the expression of IL7R; and the CDK1/2/9 inhibitor CGP-60474^88–90^, which has been reported as a potent booster of immune checkpoint blockades (**Fig. 3g, h**).

Cross-tabulating the four cell cycle-arrest classes against the four differentiation archetypes across all 62 compounds rejected global independence (*χ*^2^ = 20.34,df = 9, *p* = 0.0159, Cramér’s *V* = 0.331; **Fig. 3i**), partitioning the library into 14 dual-active (22.6%), 15 cell-cycle-only (24.2%), 14 cell-state-only (22.6), and 19 universally neutral (30.6%) compounds. Half of all G1-arrest compounds (5/10: Deforolimus, Temsirolimus, Sapanisertib, PHA-793887, and ruxolitinib) arrested differentiation in C1-“Naive-arrest” (62.5% of C1, standardized residual *z* = +3.27, *p* < 0.001), confirming that early *G*1 → *S* progression and metabolic blas-togenesis are required to exit naive quiescence^78^. By contrast, 80.0% of C3-“Memory-shifting” compounds (8/10: decitabine, TWS-119, CGP-60474, Wortmannin, EX-527, NVP-AUY922, JNK-9L, and BRD-K76674262) spared cell-cycle progression (*z* = +1.16; **Fig. 3i**).

Despite these structured phenotypes, broad Mechanism of Action (MoA) categories failed to predict T-cell differentiation shifts at the global omnibus level across all eight high-level MoA classes (binary shifting *χ*^2^ = 10.81, df = 7, *p*_perm_ = 0.1505, Freeman-Halton *p* = 0.1528; **Extended Data Fig. 44** and **Extended Data Fig. 5a**). By contrast, multi-resolution enrichment across expressed target families revealed a sharp division of labor between kinase and non-kinase modulators (**Extended Data Fig. 5b**). In particular, canonical mTOR inhibitors routed 100% (3/3) to C1 Naive-arrest (Boschloo *p* = 0.000541, *q* = 0.0065), and kinase inhibitors accounted for 87.5% (7/8) of all C1 compounds (Boschloo *p* = 0.00755, *q* = 0.0151), whereas non-kinase modulators (*n* = 14) preferentially drove C2-“Effector-promoting” (6/14) and C3-”Memory-shifting” (7/14) differentiation (*OR* = 13.00, Boschloo *p* = 0.00711, *q* = 0.0142).

To interrogate why some compounds can induce cell-state shift while others fail, we quantified the mRNA expression of drug targets across the control CD8+ T cells (*N*_DMSO_ = 6,472; **Extended Data Fig. 5c, d**). We noticed most intracellular kinase targets were ubiquitously expressed across all five T-cell states (median DMSO detection 66.59%, range 49.1% to 96.5%), whereas solid-tumor RTK and B-cell/myeloid spleen tyrosine kinase (*SYK*) targets were nearly absent (median detection 1.64%; **Extended Data Fig. 5c, d**). Because mature human CD8+ T cells transduce TCR–CD3*ζ* signals through *ZAP70*^*91*^ (74.89% detection) rather than its B-cell and myeloid paralog *SYK* (0.99% detection), the selective SYK inhibitors entospletinib^92^ and fostamatinib^93^ remained silent on *u*_*state*_; as did six non-lymphoid RTK, Hedgehog, and dopamine modulators lacking target expression (≤ 5 detection). Conversely, the only two active compounds whose canonical primary targets fell below 5% detection, Neratinib (*ERBB2*, 4.67%) and NVP-BHG 712 (*EPHB4*, 0.79%), both routed to C1 Naive-arrest via reported offtarget effect against abundantly expressed T-cell kinases: Neratinib inhibits *LCK* (95.2%), *MAP4K1*/HPK1 (63.8%), and *ITK* (61.5%)^94,95^, while NVP-BHG 712 cross-inhibits *LCK* (95.2%) and *RAF1*/c-Raf (68.4%)^96^ (**Extended Data Fig. 5d**).

#### State-driving perturbation responses revealed by counterfactual analysis via z_state_

To further investigate the transcriptional driving forces underlying these compound-induced state shift, we looked into the perturbation-aware state space *z*_*state*_ that modeled the on-manifold pharmacological effects. We noted that some compounds sharing the same primary mechanism coclustered tightly (e.g., Panobinostat and Belinostat as HDACi; Deforolimus, Temsirolimus, and Sapanisertib as mTORi), confirming that PertVI’s Split-FiLM module recapitulates pharmacological transcriptional responses (**Fig. 3j**). Across all 62 compounds, PertVI’s global Momentum Score correlated strongly with contrastive MoCAVI salient displacement (Spearman *ρ* = 0.79, Pearson *r* = 0.92; **Fig. 3k**), confirming cross-model concordance of global transcriptional effect sizes. While conventional perturbation pipelines usually assume that large transcriptional shifts drive phenotypic changes, comparing the Momentum effect size against phenotypic state-shift magnitude on *u*_*state*_ (**Fig. 3l** and **Extended Data Fig. 6a**), however, revealed that global Momentum explained only 15.30% of state-shift variance across the 62 compounds, weakly correlating with state-shifting magnitude (outlier-excluded, *r* = 0.16, *p* = 0.2221 *NS*). Indeed, we found nearly half of active state shifters (13/28 = 46.4) fell into Quadrant Q2 as low-Momentum lineage-gating catalysts (**Fig. 3l**; **Extended Data Fig. 6a**), exemplified by CGP-60474 (*L*_2_ = 1.95, rank 62 of 62), dabrafenib (3.27), YM-155 (3.42), camptothecin (3.52), neratinib (3.57), TWS-119 (4.73), PHA-793887 (5.03), and decitabine (8.01). Conversely, seven non-shifting compounds fell into Quadrant Q3, including calcitriol (*L*_2_ = 32.81, rank 4 of 62), pomalidomide (21.23), and belinostat (20.76), alongside GW-843682X, apicidin, BMS-536924, and CYT-387 (*L*_2_ = 8.34 to 11.11), indicating potential induction of state-invariant transcriptional programs along orthogonal axes that remained home-ostatically buffered with respect to *u*_*state*_ differentiation (**Fig. 3l**; **Extended Data Fig. 6a**). By integrating pairwise functional correlations across all compound pairs (**Extended Data Fig. 6b-g;** see **Methods**), we further revealed that compounds sharing both MoA and archetype showed the strongest regulatory concordance (*r*_TF_ = +0.59, *r*_PW_ = +0.41), followed by compounds of different MoA but sharing the same archetype (within C3-”Memory-promoting”, *r*_TF_ = +0.42, *r*_PW_ = +0.27; within C1-”Naïve arrest”, *r*_TF_ = +0.38, *r*_PW_ = −0.02 NS; within C2-”Effector-promoting” were almost uncorrelated, *r*_TF_ = −0.04 NS, *r*_PW_ = +0.03 NS).

We therefore hypothesized that a compound’s state-driving response may be fundamentally decoupled from its global perturbation response and textbook MoAs, and that lineage transitions are governed by localized key regulatory events at specific developmental checkpoints rather than bulk, transcriptome-wide differential expression. To test this hypothesis, we used PertVI’s Split-FiLM decoder to evaluate state-conditioned counterfactual shifts (CSS_*s*_) within each developmental state (See **Methods**). Interestingly, we found that compounds in C1-”Na-ïve arrest” suppressed early immune activation and trafficking pathways in both Momentum score and all differentiation stages, including lymphocyte chemotaxis (net vote = −0.625, mean *NES* = −1.56, *q*_shared_ = 0.0045) and cellular response to chemokine (net = −0.50, mean NES = −0.59, *q*_shared_ = 0.0251), while activating T-cell tolerance and quiescence regulons NR4A2^97,98^ (net = +0.88, *t* = +2.80, *p*_sp_ = 0.0013) and ATF3^99^ (net = +0.625, *t* = +1.61, *p*_sp_ = 0.0061). Within C1, regulation further bifurcated into two sub-modules (**Extended Data Fig. 6e, f, g**). In sub-module C1a (*n* = 5), four mTOR and CDK2/4 inhibitors coactivated anti-proliferative TGF-*β* /SMAD3 signaling (Momentum *t* = +2.12 *to* + 2.33, reaching +2.67 in CSS_Tn.prolif_ for Temsirolimus^100^). In sub-module C1b (*n* = 3: ruxolitinib, Neratinib, and Panobinostat), compounds instead suppressed transcription factor SMAD3 (*t* = −1.16 *to* − 1.84) and blocked cytokine-driven STAT3/STAT6 and E2F4 activity^101^ (ruxolitinib STAT3/STAT6 *t* = −2.39 *to* − 3.13 in CSS_Tn_ and CSS_Tn.prolif_).

Conversely, C2-“Effector-promoting” compounds converged in mature effectors on proin-flammatory chemokine-mediated signaling (net = +0.70, mean NES = +1.30, *q*_shared_ = 2.09 × 10^−4^), cellular response to chemokine (net = +0.70, mean NES = +1.28, *p*_sp_ = 0.0067), and TNF-mediated signaling (net = +0.80, mean NES = +1.23, *q*_shared_ = 4.94 × 10^−5^), despite low global correlation across structurally diverse drug classes (**Extended Data Fig. 6e, f, h, i**). Unexpectedly, C3 (Memory-shifting) compounds showed zero cluster-wide consensus across all six canonical memory transcription factors (Block TF-3). Rather than directly activating memory transcription factors in naive cells, C3 compounds (including both hypomethylating Decitabine and Wnt-activating TWS-119 in sub-module C3a) appear to converge in late effector states on suppressing RNA modification (active-voter NES = −1.80, *p*_sp_ = 1.15 × 10^−5^) and SSU-rRNA maturation (active-voter NES = −2.01, *p*_sp_ = 2.88 × 10^−5^; **Extended Data Fig. 6e, f, h, i**). This late-stage biosynthetic and mitotic brake aligns with reports that restraining ribosomal biogenesis and translation during clonal expansion favors memory precursor formation in murine and human T cells^102,103^. At the transcription factor level, this program was coupled to coordinated repression of the E2F/DREAM cell-cycle regulons^104^ E2F4 (active-voter *t* = −2.24, *p*_sp_ = 0.00025,), E2F2 (active-voter *t* = −1.92, *p*_sp_ = 0.00060), and TFDP1 (*p*_sp_ = 0.0027), alongside suppression of the chromatin architectural factor CTCF in CSS_T.eff_ (active-voter *t* = −1.99, *p*_sp_ = 0.00060), which is required for terminal effector chromatin looping^105,106^.

As a specificity control (Block TF-6 and PW-6), ectopic non-T-cell lineage regulons (hepatocyte HNF4A, myogenic MYOD1, and neuronal NEUROD1) and pathways (cardiac muscle contraction, chemical synaptic transmission, and negative regulation of ossification) were completely null across all archetypes and states (*q*_shared_ = 1.0, *q*_sp_ = 1.0), and nominal signals from unexpressed non-T-cell factors (osteoblast SP7 at 0.0 single-cell mRNA detection, B-cell PAX5 and EBF1, and myeloid CEBPA) were identified and excised as CollecTRI target-overlap artifacts (see **Methods**).

#### Off-manifold cellular stress and toxicity disentanglement via v_ood_

Having mapped how compounds redirect T cells along physiological proliferation and differentiation trajectories, we next addressed our third question: which perturbations induce aberrant, off-manifold cellular stress or toxicity? To this end, we quantified each compound’s off-manifold displacement *d*_*ood*_ as the Euclidean distance of its treatment centroid from the DMSO control centroid within the anomaly subspace *v*_*ood*_ (**Fig. 3m** and **Extended Data Fig. 7a**; see **Methods**). A two-component Gaussian Mixture Model (*µ*_1_ = 0.0885 versus *µ*_2_ = 0.4082; operational cutoff *d*_*ood*_ ≥ 0.30) partitioned the screen into 14 off-manifold stress-inducing compounds (22.6) and 48 onmanifold compounds clustered near the physiological origin, with five extreme outliers exceeding the Tukey fence (*d*_*ood*_ ≥ 0.5623 : ruxolitinib 1.156, serdemetan 0.719, BMS-536924 0.677, thapsigargin 0.670, and sapanisertib 0.662; **Extended Data Fig. 7a**). Specifically, 62.5% (5/8) of C1 Naive-arrest drugs were routed off-manifold (library-wide *OR* = 8.33, *p* = 0.0113), compared with only 20.0% (2/10) of C2 (thapsigargin 0.670 and I-BET 762 0.425) and 20.0% (2/10) of C3 (serdemetan 0.719 and wortmannin 0.521) compounds (**Fig. 3n**). Across MoA classes, 100% of JAK inhibitors (3/3; Fisher’s exact *p* = 0.0096), 100% of mTOR inhibitors (3/3; *p* = 0.0096), 100% of combined PI3K/mTOR inhibitors including wortmannin (4/4; *p* = 0.0018), and 66.7% of HDAC inhibitors (2/3: belinostat and panobinostat; *p* = 0.13) triggered significant off-manifold deviation (**Extended Data Fig. 7b**). Because CD8+ T cells in this screen are driven by anti-CD3/CD28 co-stimulation and IL-2, they obligately depend on JAK-STAT and PI3K-mTOR signaling for TCR blastogenesis and metabolic survival^78^; pharmacological blockade of these essential pathways induces acute deprivation stress that PertVI routes into *v*_*ood*_. Likewise, although Thapsigargin accelerated effector differentiation on *u*_*state*_ (C2), PertVI isolated its physical endoplasmic reticulum calcium-depletion stress within *v*_*ood*_ (*d*_*ood*_ = 0.670).

Separating permitted on-manifold trajectory shifts from off-manifold toxicity is critical for discovering T-cell therapy adjuvants that redirect lineage fate without compromising cellular fitness. To characterize the off-manifold response across compounds, we derived the Marginalized Counterfactual Operator (MCO) to extract the off-manifold gene signatures (see **Methods**). Across the 14 off-manifold perturbations and three on-manifold controls (ponatinib, sorafenib, and BI 2536), MCO isolated concerted induction of 9 canonical oxidative, xenobiotic, and endoplasmic reticulum stress genes paired with profound suppression of 9 interferon-stimulated and cytolytic immune genes (**Extended Data Fig. 7c, d**). Further pathway analysis on MCO signatures revealed eight core pathways that significantly distinguished the 14 off-manifold compounds from the rest 48 on-manifold compounds (**Fig. 3o**). Unsupervised clustering further stratified the 14 off-manifold drugs into six mechanism-specific toxicity classes across five functional pathway programs (**Extended Data Fig. 7e, f**). Crucially, this decomposition resolved a major blind spot within the C3 Memory-shifting archetype (**Fig. 3m, o** and **Extended Data Fig. 7e, f**). Genuine epigenetic, Wnt/GSK-3*β*, and transcriptional-pause memory adjuvants such as decitabine (*d*_*ood*_ = 0.175), TWS-119 (*d*_*ood*_ = 0.172), and CGP-60474 (*d*_*ood*_ = 0.076) expanded *IL7R*+ memory precursors cleanly on-manifold without inducing *v*_*ood*_ stress. By contrast, Serdemetan (*d*_*ood*_ = 0.719) and wortmannin (*d*_*ood*_ = 0.521) phenocopied *IL7R*+ accumulation on *u*_*state*_ while triggering severe off-manifold pathology in *v*_*ood*_. Specifically, the MDM2 inhibitor Serdemetan induced acute p53 checkpoint arrest and bioenergetic collapse (p53 pathway: *NES* = +1.36, MYC targets: *NES* = −3.56, and oxidative phosphorylation: *NES* = −1.58), whereas the covalent PI3K inhibitor wortmannin triggered apoptosis (*NES* = +1.63, *p* = 0.0097), unfolded protein response (+1.68), and an atypical interferon-*α*/*γ* storm (+3.05 and +3.26). Without an explicit anomaly subspace, conventional trajectory and differential abundance analyses would likely conflate these toxic, phenocopying agents with genuine, developmental adjuvants.

Together, by structurally decoupling permitted, on-manifold biological momentum from aberrant, off-manifold toxicity, PertVI provides a unique level of mechanistic granularity that extends and complements the current analytical capabilities of conventional single-cell perturbseq workflows.

### PertVI disentangles cell-cycle dynamics, activation states, and context-specific genetic regulators in primary human CD4+ T cells

To demonstrate that PertVI generalizes to multi-condition screens, we applied it to a pilot dataset extracted from a recent genome-scale CRISPRi (ZIM3-KRAB-dCas9) Perturb-seq atlas of primary human CD4+ T cells^15^ (**Fig. 4a**). The parent atlas targeted 12,779 expressed genes (26,504 sgRNAs including 5% non-targeting controls) across 21.99 million single-guide primary human CD4+ T cells across four healthy donors (D1–D4) and three T-cell receptor (TCR; anti-CD3/CD28/CD2) stimulation conditions: resting baseline (Rest, unstimulated), acute activation (Stim 8hr, 8h post-stimulation), and late effector maturation (Stim 48hr, 48h post-stimulation), defining 111 co-functional perturbation clusters across conditions. Across this multicondition screen, single-cell transcriptomes reflect the combined influence of inter-donor biological variation (D1–D4), chronological activation-state transitions (Rest to Stim 8hr and Stim 48hr), rapid mitotic cell-cycle entry during 48h blastogenesis, and stage-dependent genetic perturbations. To benchmark PertVI across these coexisting sources of variation as a proof of principle, we constructed a stratified pilot dataset of 120,545 high-quality single-guide cells (Rest: 42,344 cells; Stim 8hr: 40,272 cells; Stim 48hr: 37,929 cells across D1–D4) profiled across 10,004 expressed genes (the top 10,000 donor-aware highly variable genes unioned with 94 canonical cell-cycle markers^18^; **Fig. 4a** and **Extended Data Fig. 8a**). Specifically, the pilot dataset includes 61 curated T-cell activation regulators (77,003 cells) drawn from the original study’s T-cell activation clusters (Clusters 21, 22, and 99) and stratified by their active timepoints into early (active at 8h in Cluster 21, *n* = 12; 19,248 cells), general (shared across 8h in Clusters 21 and 22 and 48h in Cluster 99, *n* = 29; 34,081 cells), and late (active at 48h in Cluster 99, *n* = 20; 23,674 cells) stages across seven functional categories (See **Methods 6.1**). Alongside the annotated active regulators, we also designed two sets of controls: 20 non-essential control genes (NCGs; 37,518 cells) randomly sampled from perturbed genes strictly outside all co-functional clusters to serve as empirical null, and 6,024 uniformly sampled non-targeting control (NTC) cells (5.00% of the dataset; 502 per donor-condition pool, 2,008 per condition, and 1,506 per donor) matching the parent library’s 5% NTC proportion to anchor the unperturbed baseline. Training PertVI over 400 epochs with donor batch conditioning and perturbation Feature-wise Linear Modulation converged smoothly across ELBO, reconstruction loss, and KL divergence (**Extended Data Fig. 8b**; **Methods**). Comparing uncorrected expression with PertVI’s six latent subspaces (*u*_*cycle*_, *u*_*state*_, *z*_*cycle*_, *z*_*state*_, *v*_*ood*_, *z*_*final*_) confirmed clean architectural disentanglement (**Fig. 4b** and **Extended Data Fig. 8c**).

**Fig. 4.**
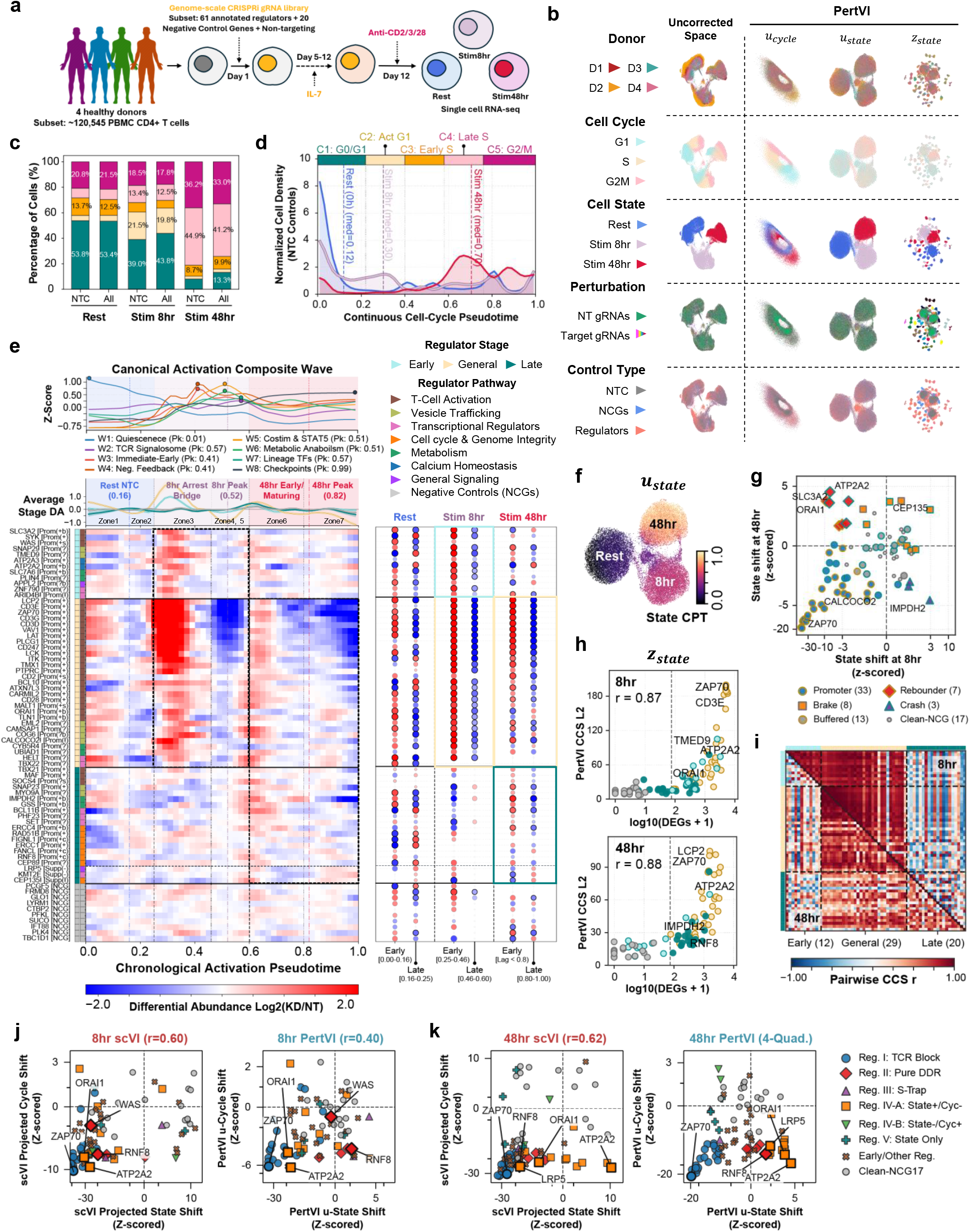
PertVI reveals context-specific regulators governing early- and late-stage CD4+ T cell activation. **a**, Experimental workflow schematic of the multi-condition primary human CD4+ T-cell CRISPRi (ZIM3-KRAB-dCas9) Perturb-seq screen (GSE314342). **b**, UMAP embeddings (*n* = 120,545 cells) comparing Uncorrected Space (column 1) against PertVI’s 3D basal cell-cycle subspace (*u*_cycle_, column 2), 30D basal cell-state subspace (*u*_state_, column 3), and 30D perturbation-modulated state subspace (*z*_state_, column 4), colored by Donor (D1–D4, row 1), canonical Cell Cycle phase (G1, S, G2M, row 2), Cell State condition (Rest, Stim 8hr, Stim 48hr, row 3), Perturbation status (NT gRNAs versus Target gRNAs, row 4), and Control Type (NTC, NCGs, Regulators, row 5). **c**, Stacked bar chart of cell percentages (Percentage of Cells (%)) across the five continuous cell-cycle phase zones along *τ*_cycle_ comparing unperturbed NTC controls (*n* = 6,024) and All cells (*n* = 120,545) across Rest, Stim 8hr, and Stim 48hr. **d**, Kernel density distributions of Normalized Cell Density (NTC Controls) (*n* = 6,024) along Continuous Cell-Cycle Pseudotime (*τ*_cycle_ ∈ [0.0, 1.0]) on *u*_cycle_. Top color bar and vertical grey lines mark boundaries of zones C1–C5; vertical dashed lines indicate condition medians. **e**, Chronological Activation Pseudotime (CPT, *τ*_state_ ∈ [0.0, 1.0]) landscape on *u*_state_. Top (Canonical Activation Composite Wave): Standardized *Z*-score expression trajectories of eight sequential canonical T-cell activation waves across 50 uniform *τ*_state_ bins. Middle (Average Stage DA): Stage-averaged within-condition differential abundance (log_2_(KD/NT), solid curves with ±1 s.e.m.shaded bands) along *τ*_state_ for Early (*n* = 12), General (*n* = 29), and Late (*n* = 20) regulators and Negative Controls (NCGs) (*n* = 20) across developmental milestones (Rest NTC (0.16), 8hr Arrest Bridge, 8hr Peak (0.52), 48hr Early/Maturing, and 48hr Peak (0.82)) and seven contiguous developmental zones (Zone1– Zone7). Bottom left: Heatmap of within-condition Differential Abundance Log2(KD/NT) across 50 *τ*_state_ bins for target regulators and representative NCGs. Bottom right: Row-aligned discrete developmental zone differential abundance lollipop profiles across Rest (Early [0.00–0.16] versus Late [0.16–0.25]), Stim 8hr (Early [0.25–0.46] versus Late [0.46–0.60]), and Stim 48hr (Early [<0.8] versus Late [0.80–1.00]). **f**, UMAP embedding of the 30D basal state subspace (*u*_state_, *n* = 120,545 cells) colored by continuous Chronological Activation Pseudotime (State CPT, *τ*_state_ ∈ [0.0, 1.0]) across Rest, 8hr, and 48hr territories. **g**, Two-dimen-sional signed phenotypic state-shift plane comparing Clean-NCG17-standardized State shift at 8hr (z-scored) (*Z*_*u*,state,8h_, *x*-axis, symmetric logarithmic scale) versus State shift at 48hr (z-scored) (*Z*_*u*,state,48h_, *y*-axis, symlog scale) across all 81perturbations, classifying perturbations into Promoter (33), Rebounder (7), Brake (8), Crash (3), Buffered (13), and Clean-NCG (17). Dashed lines indicate *Z* = 0. **h**, Scatterplots comparing donor-pooled PyDESeq2 log_10_(DEGs + 1)(FDR < 0.05, *x*-axis) from the original study against PertVI counterfactual causal score effect size (PertVI CCS L2, ∥ CCS ∥_2_, *y*-axis) decoded from *z*_state_ across all 81perturbations at 8hr (top, *r* = 0.87) and 48hr (bottom, *r* = 0.88), colored by perturbation stage (Early, General, Late, NCG). **i**, Split-triangle pairwise Pearson correlation heatmap of gene-level *z*_state_ counterfactual causal score (CCS) profiles across all 61 target regulators, displaying 8hr correlations in the upper tri-angle and 48hr correlations in the lower triangle. **j**, Side-by-side scatterplots (symlog axes) at Stim 8hr comparing Clean-NCG17-standardized projected state shift (*x*-axis) and cycle shift (*y*-axis) in the unconstrained 30D scVI model versus PertVI basal *u*_state_ and *u*_cycle_ shifts across all 81perturbations. **k**, Side-by-side scatterplots (symlog axes) at Stim 48hr comparing Clean-NCG17-standardized projected state shift (*x*-axis) and cycle shift (*y*-axis) in 30D scVI versus PertVI across all 81 perturbations, colored and shaped by 48h mechanistic regime.

#### Cell-cycle ring reconstruction reveals stimulation-coupled proliferation changes on u_cycle_

Within the basal cell-cycle subspace (*u*_cycle_), PertVI recovered the closed ring topology of the mitotic cell cycle across all 120,545 cells, parameterized along a 6-waypoint polar geodesic (*τ*_cycle_ ∈ [0, 1], spanning five sequential phase zones: C1: G0/G1, C2: Activated G1, C3: Early S, C4: Late S, and C5: G2/M; **Extended Data Fig. 8d**,**e**; see **Methods**). Along *τ*_cycle_, canonical cell-cycle markers resolved into five sequential transcriptional waves from G0/early-G1 quiescence (W1) and G1/S restriction-point licensing (W2) through S-phase DNA replication (W3), G2/M checkpoint control (W4), and cytokinesis exit (W5). These wave trajectories were nearly identical between unperturbed NTC controls (*n* = 6,024) and all 120,545 cells (5-wave Pearson *r* = 0.995–1.000, median *r* = 0.9994; marker-level median Pearson *r* = 0.9998; **Extended Data Fig. 8f**,**g**), confirming that *u*_cycle_ provides a perturbation-agnostic cell-cycle coordinate. Tracking cell density along *τ*_cycle_ revealed staged, progressive proliferation upon TCR restimulation (**Fig. 4c, d**), advancing T cells from basal quiescence at Rest (53.8% in C1, median *τ*_cycle_ = 0.12) through acute G1 restriction-point entry at Stim 8hr (marked by a 5.21-fold surge in C2 from 4.1% to 21.5%, median *τ*_cycle_ = 0.30) to full-scale S and G2/M clonal blastogenesis at Stim 48hr (81.1% in C4 and C5, median *τ*_cycle_ = 0.70). Within-condition differential abundance against NTC across zones C1–C5 further revealed that the early (*n* = 12), general (*n* = 29), and late (*n* = 20) regulators classified for T-cell activation exhibit distinct stage-specific proliferation regulation across 8hr and 48hr post stimulation (**Extended Data Fig. 8h**). General regulators, dominated by the proximal TCR and co-stimulatory signalosome, gated both acute 8h G1 restriction-point licensing and 48h mitotic expansion. By contrast, early activation regulators spared acute activated G1 licensing at 8hr (0 of 12 depleting C2 at FDR < 0.05; median log_2_DA_*C*2_ = −0.01) and largely resumed S/G2/M cycling by 48hr (0 of 12 depleting C4 at FDR < 0.05). Conversely, at Stim 8hr, late regulators spared acute activated G1 entry (0 of 20 depleting C2 at FDR < 0.05; median log_2_DA_*C*2_ = −0.13) and only showed mild retention in G0/G1. By Stim 48hr, late regulators converged on pronounced G0/G1 enrichment (11 of 20 enrich C1 at FDR < 0.05; median log_2_DA_*C*1_ = +0.67).

#### T cell activation continuum reconstruction reveals phenotypic stage-hierarchy on u_state_

In parallel, the basal state subspace (*u*_state_) recovered the continuous progression of CD4+ T-cell activation across at rest, 8hr post-stimulation, and 48hr post-stimulation, parameterized along Chronological Activation Pseudotime (CPT, *τ*_state_ ∈ [0, 1]; **Fig. 4e, f**; see **Methods 6.2**). Along *τ*_state_, unperturbed NTC cells progressed monotonically from Rest (median *τ*_state_ = 0.16) through Stim 8hr (0.52) to Stim 48hr (0.82; Spearman *ρ* = +0.9251 in NTC and *ρ* = +0.9142 across all 120,545 cells; Kruskal-Wallis *H* = 104,909.6, *P* < 10^−300^; **Extended Data Fig. 9a**,**b**). Projecting 75 canonical T-cell activation markers onto *τ*_state_ resolved eight sequential transcriptional waves (W1–W8; **Fig. 4e**, top, and **Extended Data Fig. 9c–e**; see **Methods**), starting with naive quiescence and lymphoid homing at the resting pole (W1), proximal TCR signalosome remodeling (W2), immediate-early ignition and acute cytokines (W3), and negative-feedback attenuators (W4). During effector maturation, these early programs transitioned into co-stimulatory and STAT5 survival signaling (W5), nucleotide, glutathione, and amino-acid anabolism (W6), helper lineage transcription factors (W7), and late homologous-recombination repair and coinhibitory checkpoint receptors at the 48h apex (W8). Expression trajectories across all 75 markers were conserved between NTC controls (*n* = 6,024) and all 120,545 cells (marker-level median Pearson *r* = 0.9912; 8-wave composite median *r* = 0.9932; **Extended Data Fig. 9c–e**), confirming the perturbation-agnostic nature of *u*_*state*_ manifold.

Because this unsorted Perturb-seq screen enriched transduced cells solely by antibiotic resistance without fluorescence-activated cell sorting (FACS) on surface or cytokine markers, direct cellular phenotypes were not physically measured; instead, the original study classified the 61 target regulators into early (*n* = 12), general (*n* = 29), and late (*n* = 20) stages indirectly via timepoint-specific differentially expressed gene (DEG) counts and unsupervised clustering. Testing within-condition differential abundance against NTC across 50 *τ*_*state*_ bins and developmental zones (**Fig. 4e, f**) confirmed that baseline-invariant phenotypic shifting along *u*_*state*_ directly recovered this stage hierarchy (4-group Kruskal-Wallis *H* = 53.54, *P* = 1.41 × 10^−11^ at Stim 8hr). Specifically, general regulators, anchored by the proximal TCR signalosome, enforced an uncompensated checkpoint across both stimulated timepoints, trapping 74.8–90.3% of 8h cells in early arrest Zone 3 (log_2_DA up to +4.24) and depleting the mature Zone 7 effector apex at 48 h (log_2_DA down to −2.58). Early regulators selectively delayed acute 8h exit from quiescence but attenuated 3.7-fold in standardized state excursion from 4.57 at 8h to 1.24 at 48h (paired Wilcoxon *P* = 2.44 × 10^−3^) as knockdown cells resumed late maturation. Conversely, late regulators (*n* = 20) bypassed acute 8h arrest (19 of 20 showing no Zone 3 enrichment, mean log_2_DA = −0.12) and instead emerged at 48h, amplifying 3.9-fold in state excursion (paired Wilcoxon *P* = 2.67 × 10^−5^) and 4.3-fold in terminal peak differential abundance (*P* = 1.91 × 10^−6^), where late maturation promoters retained cells in lagging Zone 6 (log_2_DA up to +0.74) and depleted the mature Zone 7 effector apex (log_2_DA down to −0.90). Together, these stage-resolved shifts demonstrate that *u*_*state*_ serves as a continuous in silico phenotypic recovery readout directly from unsorted single-cell transcriptomes when physical sorting phenotypes are unavailable.

Beyond recapitulating the temporal stage hierarchy, signed phenotypic shifting along *u*_*state*_ also resolved regulatory directionality across timepoints, distinguishing perturbations that arrest activation from those that accelerate it. Projecting perturbations into signed shift plane (**Fig. 4g**) identified 27 uni-directional positive promoters whose knockdown arrested activation (*Z*_*u*,state_ < −1.0), versus 6 negative-feedback suppressors whose knockdown accelerated progression (*Z*_*u*,state_ > +0.9), comprising three 48h effector differentiation restraints (*KMT2E, CEP135*, and *LRP5*; *Z*_48h_ = +0.97 to + 2.03) and three transient 8h accelerators (*MYO9A, SOCS4*, and *SET*; *Z*_8h_ = +1.07 to + 1.53). Comparing signed shifts between 8h and 48h further uncovered two classes of biphasic regulators masked by static DEG counts (**Fig. 4g**; **Extended Data Fig. 9f**): (1) seven homeostatic rebounders (three retained in Cluster 99^15^ [*ORAI1, COG6, TLN1*] and four excluded as early-only [*ATP2A2, SLC3A2, SLC7A6, APPL2*]) that delayed acute 8h activation before reversing polarity by 48h to drive effector hyper-activation, and (2) three metabolic/DNA-repair crash switchers (*IMPDH2, GSS*, and *ERCC4*) that inverted from mild 8h acceleration to 48h maturation collapse. Among them, *ATP2A2* is independently validated by the original study’s accompanying arrayed CRISPRi bulk RNA-seq^15^ (GSE314342) that knockdown reversed transcriptome-wide directionality across 649 TCR-activated genes and 645 TCR-repressed quiescence genes (**Extended Data Fig. 9g**). In addition, within the 20 randomly sampled NCGs, PertVI also rescued three bona fide regulators missed by donor-pooled PyDESeq2 at 48 h (0–13 pooled DEGs) that exceeded the 1.90 SD null ceiling (**Fig. 4e**; **Extended Data Fig. 9h, i**): the Rab-GAP metabolic brake *TBC1D1*^*107*^ (+5.11 SD, *Z*_*u,state*,48*h*_ = +2.13, *Z*_*u,cycle*,48*h*_ = −3.04), the non-ciliary immunological synapse TCR-recycling subunit *IFT88*^*108,109*^ (+2.96 SD, *Z*_*u,state*,48*h*_ = +1.67), and the centriole duplication kinase *PLK4*^*110*^ (+5.74 SD, *Z*_*u,cycle*,48*h*_ = −5.73, omitted as untestable by PyDESeq2 at 48h after mitotic depletion to *n* = 51 cells).

#### Counterfactual analysis recapitulates state-dependent transcriptional response

We next interrogated the perturbation-modulated state space (*z*_*state*_) to resolve the condition-specific transcriptional responses and inter-regulator relationships identified by the original study^15^ (**Fig. 4h, i**; **Extended Data Fig. 10a, b**). For each perturbation and stimulation condition, we computed gene-level counterfactual causal scores (CCS) by passing unperturbed control cells through the learned Split-FiLM transition in *z*_*state*_ and summarized overall transcriptional effect size by the *L*_2_ norm (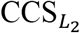; see **Methods**). Across all 81 perturbations, PertVI’s 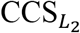 effect size correlated strongly with the DEG counts from the original study at both Stim 8hr (*r* = 0.87) and Stim 48hr (*r* = 0.88; **Fig. 4h**). Comparing 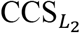 across timepoints further resolved the stage hierarchy among early, general, and late regulators (**Extended Data Fig. 10a, b**), demonstrating that PertVI accurately derives context-specific transcriptional responses. Beyond overall effect size, pairwise correlations of gene-level CCS profiles across all 61 target regulators reproduced the stage-specific co-functional relationships reported in the original study, capturing the co-clustering of early and general regulators at Stim 8hr (*r* = 0.84, upper triangle) and the transition to a shared general and late regulator module at Stim 48hr (median *r* = 0.63, lower triangle) as early perturbation effects subsided (*r* = −0.06; **Fig. 4i**).

To examine the functional programs underlying these stage-specific patterns, we projected gene-level *z*_*state*_ counterfactual causal scores onto curated CollecTRI transcription factor regulons (**Extended Data Fig. 10c, d**) and GOBP pathways (**Extended Data Fig. 10e, f**; see **Methods**). At Stim 8hr, knockdown of early and general promoters prevented the normal TCR-induced silencing of the naive quiescence transcription factor KLF2^111^ while coordinately suppressing immediate-early AP-1 and NF-*κ*B regulons (FOS, JUN, NFKB1, REL), the blastogenic transcription factor MYC], E2F1, ribosome biogenesis (Maturation of SSU-rRNA, Cytoplasmic Translation), and IL-2 signaling, whereas late-promoting knockdowns remained largely quiescent (**Extended Data Fig. 10c, e**). By Stim 48hr, early regulators (*SLC3A2, WAS, SYK*) resolved KLF2 retention and rebounded toward baseline across MYC, E2F1, and rRNA maturation programs; by contrast, general and late-promoting regulators (*IMPDH2, GSS, SNAP23, BCL11B*) converged on sustained suppression of MYC, the amino-acid stress regulator ATF4, SSU-rRNA maturation, and RNA modification (**Extended Data Fig. 10d, f**). Conversely, knockdown of late-suppressing brakes (*LRP5, KMT2E, CEP135*) and biphasic home-ostatic uncouplers (*ATP2A2, ORAI1*) drove mirror-image 48h upregulation of STAT1, the Th1 lineage factor TBX21, E2F1, and NF-*κ*B/NFAT regulons alongside lymphocyte activation, calcium-mediated signaling, and interferon-*γ* response pathways, while Clean-NCG17 perturbations and ectopic non-lymphoid pathway controls remained silent across both timepoints (**Extended Data Fig. 10c–f**).

#### PertVI eliminates mitotic confounding and remains SOTA integration performance

Because each perturbation cohort in this screen is distributed across four donors, standard donor-pooled pseudobulk differential expression is vulnerable to small-sample dispersion noise. To test whether PertVI preserves state-of-the-art integration and denoising performance, we trained a dimensionality-matched, unconstrained 30-dimensional scVI model^30^ (30D scVI), which was used in the original study for single-cell embedding (**Extended Data Fig. 11a**; see **Methods**). We then compared five methods by benchmarking cross-donor split-half reproducibility (Pearson *r*between Donors D1+D2 and Donors D3+D4 for all 61 target regulators across conditions; **Extended Data Fig. 11b**). Single-cell generative pooling in both PertVI (*r* = 0.52/0.73/0.79 across Rest / Stim 8hr / Stim 48hr) and 30D scVI (0.45/0.72/0.80) increased cross-donor reproducibility 3.6-to 4.2-fold over the donor-pooled PyDESeq2 approach used by the original study (0.13/0.20/0.19); meanwhile, PertVI substantially outperformed raw single-cell observational differential expression (Raw Obs-DE; 0.20/0.37/0.48) and a donor-blocked pseudo-bulk generalized linear model (Donor PB-GLM; 0.16/0.40/0.42; **Extended Data Fig. 11b**). However, although both 30D scVI and PertVI harmonized inter-donor variation across D1–D4, the undivided latent bottleneck of 30D scVI allowed high-amplitude mitotic variance to fracture the Stim 8hr and Stim 48hr condition islands into phase-segregated G1, S, and G2/M sub-lobes (**Extended Data Fig. 11a**), whereas PertVI isolated cell-cycle progression into *u*_*cycle*_ to preserve a phase-homogenized activation manifold in *u*_*state*_ (**Fig. 4b** and **Extended Data Fig. 8c**).

To evaluate how this mitotic sub-lobe fracturing impacts perturbation inference, we projected the 30D scVI latent space onto the Chronological Activation Pseudotime (CPT) state axis (**Extended Data Fig. 11c**) and the mitotic cell-cycle score axis (**Extended Data Fig. 11d**; see **Methods**). Inside both the Stim 8hr and Stim 48hr islands, the scVI state and cycle gradient vectors aligned in nearly parallel directions (**Extended Data Fig. 11c, d**). Consequently, across all 81 perturbations, scVI state and cycle shifts exhibited strong collinearity at pre-mitotic Stim 8hr (*r* = 0.60; **Fig. 4j**, left), pulling cytoskeletal and DNA-damage checkpoints (*WAS, RNF8*) into false activation arrest alongside proximal TCR and calcium regulators (*ZAP70, ORAI1, ATP2A2*), whereas PertVI reduced 8h collinearity (*r* = 0.40; **Fig. 4j**, right) and buffered *WAS* and *RNF8* near zero on *u*_*state*_. At rapidly dividing Stim 48hr, scVI state and cycle shifts collapsed onto an entangled diagonal (*r* = 0.62; **Fig. 4k**, left) that swept pure DNA-repair checkpoint *RNF8* and opposing-sign uncouplers (*LRP5, ORAI1*) into the arrest quadrant alongside *ZAP70*, whereas PertVI resolved all six mechanistic regimes across four quadrants (**Fig. 4k**, right). Moreover, scVI inflated the empirical dispersion of the 17 NCGs by 3.9-to 8.2-fold (State/Cycle SD = 8.2/5.6 at 8hr and 5.3/3.9at 48hr), whereas PertVI confined them to a compact unit-scale null envelope (SD = 1.0/1.1 at 8hr and 0.8/0.7at 48hr; **Extended Data Fig. 11e**). Direct comparison across the 12 mitotic-conflated regulators at 48h (**Extended Data Fig. 11f**) confirmed that scVI misclassified 5 of 5 Reg. II pure cell-cycle and DNA-repair checkpoints (*RNF8*^*112*^, *PHF23, FANCL, WAS*, and *FIGNL1*^*113*^) as false activation arrests. Simultaneously, across Reg. IV-A opposing-sign uncouplers, although CPT projection preserved extreme *ATP2A2* rebound, mitotic drag in scVI inverted the sign of four hyper-activators (*LRP5, COG6, ORAI1, TLN1*) into false arrest (**Extended Data Fig. 11f**). Together, these benchmarks demonstrate that PertVI’s disentanglement retains the statistical power of deep generative integration while preventing proliferative arrest from corrupting the inferred directionality and biological identity of T-cell activation regulators.

## DISCUSSION

Single-cell perturbation screens have rapidly expanded from uniform, steady-state cell lines^6^ into dynamic, multi-condition biological systems such as differentiating and proliferating primary human lymphocytes^12,13,15^. Existing linear, contrastive, and latent-shift frameworks were predominantly developed for steady-state contexts where a static cellular baseline holds^32,35,37^. Under those static conditions or strong perturbation regimes, approaches such as CPA^56^, ContrastiveVI^32^, MrVI^41^, and MoCAVI^42^ effectively isolate treatment effects against a single background representation. As perturbation screens transition into dynamic systems, however, localized intervention signals are overshadowed by macroscopic lineage progression, periodic cell-cycle oscillations, and donor heterogeneity that covary and share pleiotropic target genes. Because existing models either compress the unperturbed baseline into a single, undivided background space or distribute covarying state signals across unsupervised splits without axis specificity, variance entanglement becomes the primary bottleneck in dynamic screens. In particular, our ground-truth simulation study (**Fig. 2**) revealed that even in an idealized log-linear additive setting devoid of high-order nonlinear epistasis, covarying cell-cycle and differentiation programs significantly mask weak, target-overlapping perturbations when compressed into an undivided baseline. In addition to the simulation, the chemical screen in differentiating and actively dividing primary human CD8+ T cells^63^ (**Fig. 3**) represents a typical real-world case of this entanglement. We found that when the cellular baseline is encoded within a unified space, high-amplitude mitotic oscillations dominate the baseline embeddings of MrVI and MoCAVI, obscuring the continuous naive-to-effector differentiation trajectory while extreme stressors (such as Thapsigargin in MrVI) or class-imbalance control-memorization artifacts (MoCAVI) distort their sample covariate or perturbation spaces. Our approach, PertVI, resolves this entanglement across both simulated and primary dynamic screens by explicitly separating periodic cell division (*u*_*cycle*_) from cellular state (*u*_*state*_) prior to perturbation conditioning, recovering an unconfounded developmental trajectory alongside sensitive, condition-specific treatment responses.

Transitioning from steady-state cell lines to dynamic primary systems also requires rethinking the geometric assumptions used to represent perturbation responses. Current generative^31,56^ and optimal-transport^29^ frameworks operate under a closed, uni-manifold assumption, modeling every chemical or genetic intervention as a trajectory shift within a single latent coordinate space. In dynamic cellular systems, however, interventions simultaneously modulate physiological trajectories and trigger acute off-manifold toxicity. PertVI replaces this closed, uni-manifold view with an open, multi-manifold architecture (**Fig. 1**). Specifically, affine Split-FiLM conditioning^61^ captures on-manifold physiological transitions along perturbation-blind baseline anchors (*u*_*cycle*_ → *z*_*cycle*_ and *u*_*state*_ → *z*_*state*_), while unaccommodated nonlinear variance routes into an origin-anchored anomaly subspace (*v*_ood_) interrogated via the Marginalized Counterfactual Operator (MCO). In differentiating primary human CD8+ T cells, this multi-manifold decomposition reveals two widely neglected relationships (**Fig. 3**). First, within the on-manifold regime, phenotypic fate redirection can be largely decoupled from global transcriptional magnitude and governed instead by localized, stage-conditioned impulses. Consequently, low-magnitude lineage-gating catalysts such as CGP-60474, TWS-119, and decitabine redirect differentiation by concentrating their transcriptional impulse at narrow developmental transition windows. Conversely, high-magnitude compounds such as calcitriol, pomalidomide, and Belinostat induce broad transcriptomic programs that remain homeostatically buffered along *u*_*state*_. Second, neither on-manifold trajectory shifts nor off-manifold stress distances alone is sufficient to prioritize therapeutic immunomodulators. Joint evaluation of on-manifold shifting and off-manifold responses separates genuine memory-precursor adjuvants (e.g., decitabine, TWS-119) from toxic phenocopies (e.g., Serdemetan and Wortmannin). Whereas genuine adjuvants remodel state occupancy cleanly on-manifold, toxic phenocopies mimic *IL7R+* memory accumulation on *u*_state_ while triggering p53-mediated bioenergetic collapse or apoptosis in *v*_ood_.

Beyond isolating on-manifold shifting and off-manifold response, structuring deep architectures around variance disentanglement unlocks the full resolution of single-cell transcriptomics to dissect context-specific mechanisms and reconstruct non-observational cellular phenotypes *in silico* when physical sorting is unavailable. In unsorted primary human CD8+ T cells exposed to small-molecule perturbations (**Fig. 3**), standard discrete gene-set scoring and bulk differential expression obscure sub-phase and stage-specific drug mechanisms. By projecting cells onto continuous angular coordinates on *u*_*cycle*_, PertVI resolves a late-S replication crisis under camptothecin and Serdemetan that discrete phase-score crossover misclassifies as mitotic arrest. Simultaneously, along the deconfounded differentiation trajectory (*z*_*state*_), state-conditioned counterfactual analysis reveals that memory-shifting compounds may influence T cell fate switch via restraining ribosomal RNA maturation^102,103^, E2F/DREAM proliferation programs^104^, and the *CTCF* chromatin-looping regulon within late differentiation stage^105,106^. In pooled genetic screens, the capacity to reconstruct non-observational phenotypes is also particularly valuable (**Fig. 4**). Classical pooled CRISPR screens rely on fluorescence-activated cell sorting of a single surface receptor, cytokine, or proliferation dye to obtain a signed phenotypic readout at the expense of transcriptomic breadth. Conversely, droplet- and probe-based Perturb-seq screens capture transduced cells across conditions without physical marker sorting, making standard approaches hard to map regulators into directional regulatory modules (e.g., Cluster 99 in stimulated CD4+ T cells). By projecting unsorted CD4+ T cells onto disentangled activation (*u*_*state*_) and mitotic (*u*_*cycle*_) manifolds, PertVI reconstructs a continuous, *in silico* sorting axis directly from raw transcriptomes. These deconfounded coordinates recover chronological stage hierarchies (Early, General, and Late activation gates), resolve unsigned clusters into signed directional trajectories that separate positive TCR signalosome promoters from tonic checkpoint brakes, and uncouple cell-cycle checkpoint arrest from canonical immunological activation. Grounded by literature evidence across both screens, our downstream biological investigation demonstrates that through structural variance disentanglement, a computational engine can recapitulate in-depth regulatory relationships directly from unsorted singlecell data, establishing a deconfounded foundation that can be extended further by integrating formal statistical causal inference^114^.

Importantly, beyond its methodological framework, our study also illustrates how re-examining high-throughput screens through a disentangled multi-manifold architecture can uncover rich, previously neglected biological landscapes. Whereas the original study that generated the primary human CD8+ T cell chemical screen used the dataset primarily to train active-learning models for compound prioritization across cancer cell lines and primary cohorts^63^, applying PertVI to this screen reveals a comprehensive immuno-oncology landscape of how anti-cancer agents interact with primary human CD8+ T-cell proliferation, differentiation, and survival (**Fig. 3**). On the mitotic manifold (*u*_*cycle*_), cross-referencing primary T-cell sub-phase arrest profiles against tumor cell-line arrest phenotypes defines four immuno-oncology proliferation classes, separating conserved anti-proliferation agents (such as mTOR inhibitors and JAK in-hibitors) from tumor-active, T-cell-sparing oncology drugs including MEK inhibitors, RTK inhibitors and several epigenetic modulators. Along with the differentiation trajectory (*u*_*state*_), PertVI further resolves how anti-cancer agents can interact with effector T cell differentiation and concludes three major archetypes as naive-arrest, effector-promoting, and memory-shifting. Integrating proliferation, differentiation, and off-manifold toxicity, PertVI reveals a comprehensive landscape of heterogenous pharmacological interaction between anti-cancer drugs and CD8+ T cell effector differentiation at a granularity that conventional approaches alone can hardly achieve.

Together, our findings point to a broader methodological transition in functional genomics. As experimental platforms increasingly shift toward dynamic, multi-condition perturbation screens, the intertwined variation of proliferation, lineage progression, and cellular stress exceeds the assumptions of analytical frameworks originally designed for steady-state baselines. Advancing what computational tools can achieve in mechanistic dissection therefore calls for deep generative architectures that evolve in tandem with multi-condition experimental designs, providing a principled foundation to uncover the biological and therapeutic insights that modern perturbation screens are designed to reveal.

## LIMITATIONS OF CURRENT STUDY

PertVI is designed around a defined set of architectural assumptions, and we note three scopespecific constraints that users should consider when applying the model.

First, the current Split-FiLM transition module indexes each cell through a single categorical perturbation identifier. While this formulation matches standard low-multiplicity-of-infection (MOI) CRISPR screens in which each cell receives a single guide RNA, as well as single-agent chemical screens profiled at a fixed concentration, a single categorical lookup embedding cannot decompose additive main effects from non-linear epistasis in high-MOI multi-guide transductions or multi-drug combinations, nor can it interpolate across continuous dose-response gradients. Extending the transition module to support compositional multi-perturbation embeddings and continuous dose covariates would generalize the framework to combinatorial and dose-resolved screens.

Second, PertVI’s baseline disentanglement relies on semi-supervised gene masking and multiobjective regularization while compressing all non-mitotic physiological variation into a single continuous state manifold (*u*_*state*_). Specifically, isolating periodic division (*u*_*cycle*_) from *u*_state_ and the anomaly subspace (*v*_*ood*_) depends on a user-supplied cell-cycle marker mask (*M*_*cycle*_) in the bifurcated decoder. Consequently, separation quality is sensitive to marker-set completeness and specificity in cell types with atypical cycling programs. Besides, the multi-encoder objective introduces higher training costs alongside hyperparameter sensitivity. Insufficient repulsion will likely cause residual signal leakage between subspaces, whereas overly aggressive regularization can attenuate genuine biological covariation. Moreover, while acute perturbation-induced off-manifold stress routes into *v*_ood_, all remaining on-manifold baseline variation is modeled as a single continuous *u*_state_ representation; when multiple endogenous physiological programs unfold concurrently, such as simultaneous branching along distinct differentiation lineages or parallel cytokine polarization trajectories, they remain jointly encoded within *u*_state_. Automating penalty calibration via held-out validation metrics and factorizing *u*_state_ into priorguided baseline sub-manifolds would broaden interpretability in systems with higher-order baseline complexity.

Third, although PertVI deploys counterfactual pushforward operators to evaluate perturbation responses while holding baseline coordinates constant, structural variance disentanglement within a variational autoencoder is distinct from formal statistical causal inference^114^. By removing observational confounders including technical batch shifts, donor compositional imbalance, mitotic phase collinearity, and off-manifold cytotoxicity, Split-FiLM parameterizes a conditional affine push-forward that captures the net downstream transcriptional displacement of an intervention relative to baseline anchors. However, this affine transition does not establish directed acyclic graph structure identifiability, invariant causal prediction across environments^114^, or Pearl’s do-calculus over gene-regulatory networks^115^, and therefore does not resolve the directed gene-by-gene cascade connecting a primary molecular target to terminal phenotype. Integration of causal graph discovery with disentangled architectures will therefore be the natural future direction to achieve the complete causal disentanglement.

## METHODS

### 1. PertVI (Disentangled Variational Inference for Perturbation Response)

#### 1.1 Problem Formulation

Let 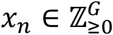 denote the raw UMI count vector for cell *n* across *G* genes, where *n* ∈ {1, …, *N*}. Each cell is associated with a perturbation identifier *s*_*n*_, encoding drug identity or guide RNA identity, and a technical covariate vector *b*_*n*_ encoding batch membership or biological replicate. PertVI factorizes the transcriptome of cell *n* as a composition of three causally disjoint latent variables: a low-dimensional cell-cycle anchor *u*_*cycle,n*_, a macroscopic biological state anchor *u*_*state,n*_, and an out-of-distribution (OOD) novelty variable *v*_*ood,n*_. The perturbation identifier *s*_*n*_ enters the model exclusively through a post-inference affine transition module, ensuring that all encoder computations remain strictly blind to the perturbation label during inference.

#### 1.2 Generative Model

*Priors*. All three latent variables are assigned standard Gaussian priors:

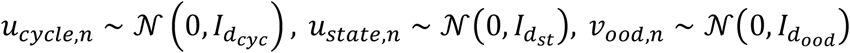

where *d*_*cyc*_ ∈ {2, 3}, *d*_*st*_ ≈ 30, and *d*_*ood*_ ≈ 10 denote the respective latent dimensionalities. The low dimensionality of *u*_*cycle*_ serves as a topological bottleneck: by constraining periodic transcriptional variance to a two- or three-dimensional manifold, the encoder is compelled to represent the continuous ring topology of the cell cycle without imposing an explicit topological prior.

##### Decoder

Gene expression is reconstructed through three structurally isolated decoder heads. Their pre-activation logits are summed before a joint SoftMax normalization:

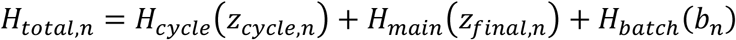

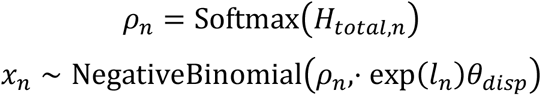

where *l*_*n*_ is a latent log-library size scalar and *θ*_*disp*_ ∈ ℝ^*G*^ is a globally trainable gene-level dispersion vector. The perturbation-conditioned vectors *z*_*cycle,n*_ and *z*_*state,n*_, and the full aggregated vector *z*_*final,n*_ = [*z*_*cycle,n*_; *z*_*state,n*_; *v*_*ood,n*_], are defined in the **1.4 Transition Module** section below.

##### Linear cycle head

*H*_*cycle*_ decodes from *z*_*cycle,n*_ through a single linear transformation governed by a hard binary gene mask *M*_*cycle*_ ∈ {0, 1}^*G*^:

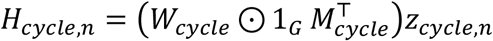

where 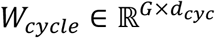 and non-zero entries in *M*_*cycle*_ correspond exclusively to pre-specified canonical cell-cycle marker genes. This hard mask prevents the cycle head from contributing reconstruction signal to non-cycle loci, enforcing a strict semantic boundary between periodic and non-periodic gene programs.

##### Dense main head

*H*_*main*_ is parameterized as a multi-layer perceptron (MLP) that receives the full aggregated representation *z*_*final,n*_. To prevent the MLP from bypassing the cycle head and reconstructing periodic expression at lower total reconstruction cost, a soft *L*_1_ penalty *ℒ*_*soft*_ is applied to the subset of the MLP’s first-layer input weight matrix that connects to cell-cycle gene output dimensions (see Objective Function). This gradient starvation constraint forces non-linear cycle decoding to remain unprofitable for the main head.

##### Batch bypass head

*H*_*batch*_ introduces a learned linear bias conditioned on the technical covariate vector *b*_*n*_, applied symmetrically across all *G* genes. The additive formulation ensures that technical batch shifts propagate only to the output logits and do not interact with the biological latent spaces or bias downstream counterfactual inference.

#### 1.3 Inference Model

Three encoder networks, one per latent space, map the observed count vector *x*_*n*_ to the parameters of diagonal-covariance Gaussian approximate posteriors. Critically, all three encoders receive only *x*_*n*_ as input; the perturbation identifier *s*_*n*_ is withheld throughout inference:

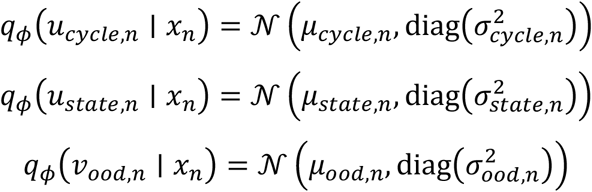

Latent samples are drawn via the reparameterization trick, for example, *u*_*cycle,n*_ = *µ*_*cycle,n*_ + *σ*_*cycle,n*_ ⊙ *ϵ* where ϵ ~ *N*(0, *I*), and analogously for *u*_*state,n*_ and *v*_*ood,n*_. A separate encoder infers the log-library size as *l*_*n*_ ~ *N*(*E*_*n*_, *I*), where *E*_*n*_ is the observed log-total UMI count for cell *n*.

- **Transition Module: The Split-FiLM Perturbation Modulation**

Perturbation identity enters the model through a dedicated FiLM transition module applied after inference, preserving the perturbation-blindness of all encoders. The perturbation identifier *s*_*n*_ is first mapped to a continuous input embedding via a learned drug embedding lookup table:

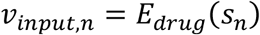

Two independent linear projection heads then generate perturbation-specific affine scale and shift coefficients for the cell-cycle and state spaces, respectively:

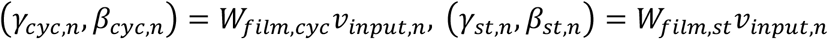

These coefficients are applied to layer-normalized baseline anchors to produce the on-manifold transcriptomic momentum vectors:

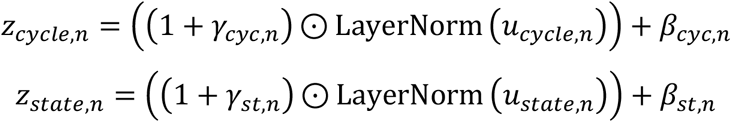

The residual formulation (1 + *γ*) ⊙ *u* + *β* has a natural Taylor expansion interpretation. Defining the composite baseline anchor *u*_*base,n*_ = [*u*_*cycle,n*_; *u*_*state,n*_] and the perturbed representation *z*_*base,n*_ = [*z*_*cycle,n*_; *z*_*state,n*_], the perturbation effect decomposes additively as:

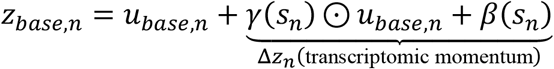

The momentum vector Δ*z*_*n*_ encodes both the direction and magnitude of the perturbation-induced displacement along the baseline biological manifold. Because the FiLM operator is restricted to element-wise affine transformations along existing latent axes, it cannot represent topological ruptures or non-linear deviations from the baseline manifold. Perturbation-induced variance that exceeds this linear expressivity, such as cytotoxic endoplasmic reticulum stress or apoptotic signaling, is rejected from the FiLM mechanism and routes instead into the OOD novelty variable *v*_*ood,n*_, which bypasses the transition module entirely. The full aggregated representation passed to the main decoder is:

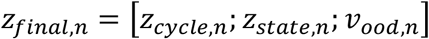

#### 1.5 Objective

The total training objective is a weighted sum of the negative ELBO and five targeted regularization penalties:

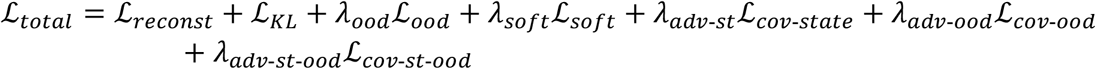

##### Reconstruction loss

The negative log-likelihood under a Negative Binomial observation model:

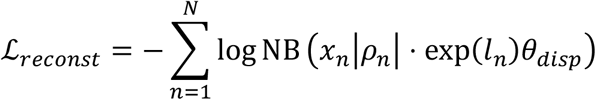

##### Total KL divergence

All latent spaces are regularized against their respective standard Gaussian priors:

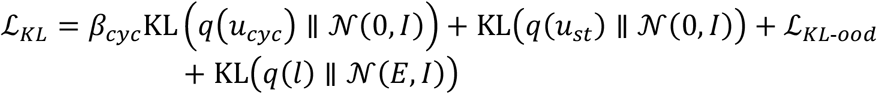

The KL weight for *u*_*cycle*_ is down-weighted by a scalar *β*_*cyc*_ ∈ (0, 1), following the *β*-VAE convention, to allow the two- or three-dimensional cycle manifold to deviate from the prior sufficiently to form a closed ring topology without posterior collapse. The OOD KL term *ℒ*_*KL*-*ood*_ is applied asymmetrically with respect to treatment status *t*_*n*_ ∈ {0, 1}: for control cells (*t*_*n*_ = 0), only the variance term is penalized,

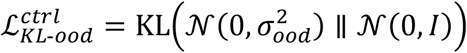

while for treated cells (*t*_*n*_ = 1), the full KL including the posterior-mean is applied:

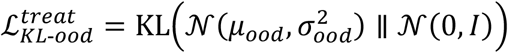

This asymmetry anchors unperturbed control cells to the origin of the OOD space, defining the unperturbed state as the causal reference, while granting treated cells the freedom to deviate in proportion to their off-manifold perturbation response.

##### Penalty (i), OOD L_1_ anchor

To mathematically define the origin of *v*_*ood*_ as the unperturbed biological baseline, an *L*_1_ sparsity penalty is applied exclusively to the OOD posterior mean of control cells (*t*_*n*_ = 0):

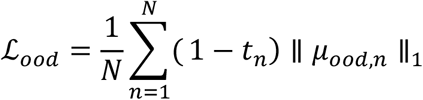

Treated cells are exempt from this penalty, allowing *µ*_*ood,n*_ to expand freely in response to off-manifold perturbation stress.

##### Penalty (ii), Soft L_1_ decoder mask

A soft *L*_1_ penalty on the intersection of the main head’s input weight matrix *W*_*main*_ and a cell-cycle gene indicator mask *M*_*cycle*_ enforces gradient starvation on non-linear cycle reconstruction:

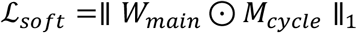

##### Penalty (iii), State-cycle cross-covariance repulsion

The squared Frobenius norm of the cross-covariance matrix between column-centered *U*_*state*_ and *U*_*cycle*_ across the minibatch of *N* cells:

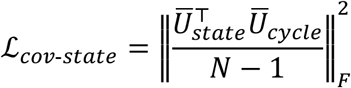

This penalty keeps the macroscopic differentiation trajectory orthogonal to the cell-cycle manifold across all cells regardless of perturbation status.

##### Penalty (iv), OOD-cycle cross-covariance repulsion

Applied exclusively to the treated-cell minibatch subset (*t*_*n*_ = 1, *N*_*t*_ cells) to prevent cell-cycle pleiotropy from contaminating the OOD space:

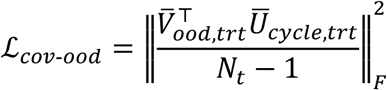

This penalty is deliberately shielded from control cells. Because control-cell OOD posterior means are origin-crushed by *ℒ*_*ood*_, including them in the cross-covariance computation would introduce artificial repulsion gradients that distort the continuous ring topology of *u*_*cycle*_.

##### Penalty (v), State-OOD cross-covariance repulsion

Applied to treated cells only, preventing macroscopic state variance from leaking into the novelty space:

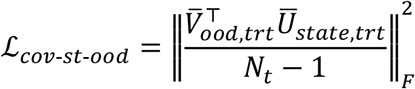

All penalty weights {*λ*_*ood*_, *λ*_*soft*_, *λ*_*adv*-*st*_, *λ*_*adv*-*ood*_, *λ*_*adv*-*st*-*ood*_} and the cycle KL scalar *β*_*cyc*_ are treated as hyperparameters. Recommended values and a tuning protocol are provided in the Supplementary Methods.

### 2. Post-training Counterfactual Analyses

Overall, PertVI enables three major downstream counterfactual analyses, each of which queries the trained main head by contrasting perturbed and control latent inputs while holding nuisance inputs fixed. The first, a global path-integrated momentum score (2.1), integrates the effect of each perturbation’s learned FiLM modulation along a path through basal state space, such as an activation or differentiation trajectory. It summarizes the average transcriptional drive of the perturbation across that continuum and, optionally, how strongly the drive varies along the path. The second, a counterfactual causal score (2.2), asks how the trained decoder predicts that control cells of the same condition would respond if they were shifted by the perturbation’s empirical displacement in basal state space. This is evaluated either with the perturbation’s own FiLM modulation or with the control modulation, giving condition-resolved, model-based estimates of perturbation effects at the gene and pathway level. The third, a marginalized coun-terfactual operator (2.3), first screens perturbations by how far their cells are displaced in the OOD subspace. It then extracts gene-level OOD signatures by substituting each perturbation’s OOD centroid into the decoder across a single fixed control reference population, so that the perturbation’s own state distribution does not enter the evaluation. Together, these modules separate on-manifold state transitions from off-manifold responses, and the detailed derivation of each module is presented below.

#### 2.1 Global Path-integrated Momentum Score

Let 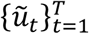 be an ordered path in LayerNorm-transformed basal state space. The OOD input is clamped to 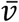 and the batch embedding to 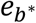, both shared across all steps and perturbations. The hidden contrast at step *t* is

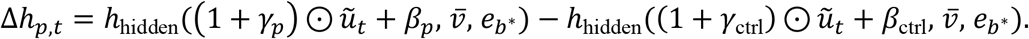

By default, 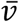 is the control-mean OOD vector (zero if none is stored) and *b*^*\**^ is the most frequent control batch. The momentum score is the path-averaged main-head logit difference. Because the readout is linear, *W*_head_ factors out of the average:

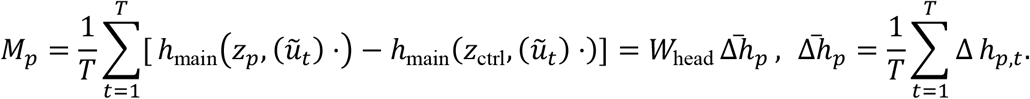

Two optional summaries describe how the contrast varies along the path, the trajectory interaction variance (TIV) and the landmark divergence (LD):

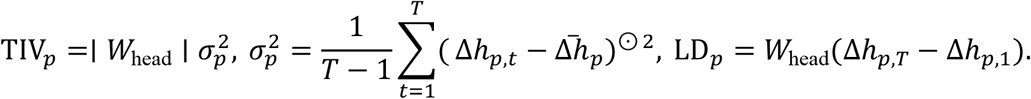

The momentum score summarizes, on the logit scale, how a perturbation’s learned state FiLM operator differs from the control operator, averaged along a user-defined path in basal state space rather than at one point. For *M*_*p*_, LD and path-integrated momentum, accumulating in the *d*_*h*_-dimensional hidden space is algebraically identical to accumulating gene-level differences.

Three path styles are available through style parameters. The default *linear* path interpolates uniformly between the mean transformed coordinates of a start and an end condition, 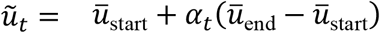, with *α*_*t*_ = (*t* − 1)/(*T* − 1) and *T* = 1,000 by default. This straight chord can cross regions that no cell occupies. The *pseudotime-binned* path orders control cells by a supplied pseudotime, restricting them to the start and end conditions when both are given. It splits the pseudotime range into 100 bins by default, with quantile edges by default or equal-width edges, and uses the bin-means as path points. The *waypoint* path starts at the control centroid of the start condition. It then visits pseudotime-binned centroids, 50 bins by default, formed from control cells of the end condition, or of all other conditions when no end condition is given. When no such control cells exist, all cells are used. Consecutive nodes are joined by *S* linear sub-steps (20 by default), which gives *T* = *WS* + 1 points for *W* retained waypoints, or 1,001 when every bin is populated.

#### 2.2 Counterfactual Causal Score (CCS)

Within each condition *c*, let *C*_*p,c*_ be the cells carrying perturbation *p* and *C*_0,*c*_ the control cells of the same condition, with input coordinates *u*_*i*_. The empirical latent state displacement and the counterfactual decoder input of control cell *j* ∈ *C*_0,*c*_ are

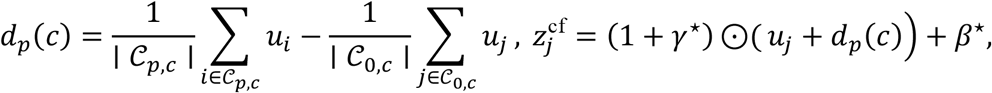

where (*γ*^*⋆*^, *β*^*⋆*^) = (*γ*_*p*_, *β*_*p*_) for Coupled-CCS (the default) and (*γ*_ctrl_, *β*_ctrl_) for State-CCS. The reference input is *z*^0^ = (1 + *γ*_ctrl_) ⊙ *u*_*j*_ + *β*_ctrl_. With the OOD input fixed at *v*_0_1 and the batch embedding at 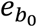, the score is

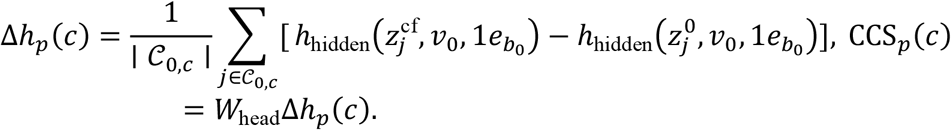

At the decoder input, the Coupled-CCS shift decomposes exactly as follows, and its first term alone is the State-CCS shift:

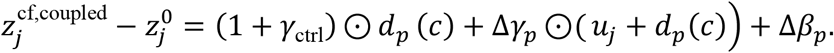

CCS is a model-based counterfactual. It asks how, according to the trained decoder, the mainhead logits of the control cells in condition *c* would change if each cell were displaced by the perturbation’s observed mean state shift and, in coupled mode, also modulated by the pertur-bation’s FiLM operator. State-CCS gives the consequence of relocating cells along the basal manifold under control regulatory rules, whereas Coupled-CCS adds the perturbation’s direct FiLM remodeling.

#### 2.3 Marginalized Counterfactual Operator for OOD Signatures (MCO)

For each perturbation with at least five cells, pooled over conditions, the OOD screen compares centroids:

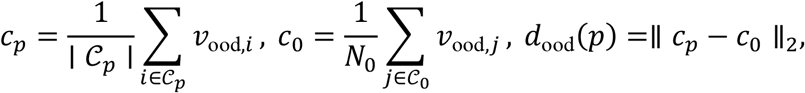

where *C*_0_ is the set of all *N*_0_ control cells. The marginalized counterfactual operator (MCO) replaces the OOD input of every control cell by the perturbation centroid. Each control cell keeps its own stored state input *z*_*j*_ and batch *b*_*j*_:

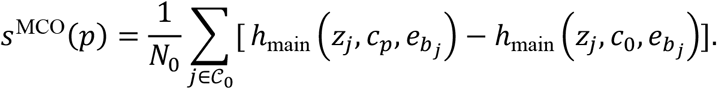

In evaluation mode BatchNorm is an affine map with linear part 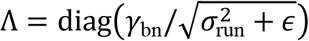. Let 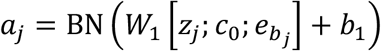 be the pre-activation of control cell *j* at the baseline, and let 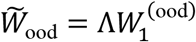 be the rescaled OOD columns of *W*. Linearizing in *v* at *c* and averaging the Jacobian over *n* control cells gives the canonical linear tangent operator (CLTO):

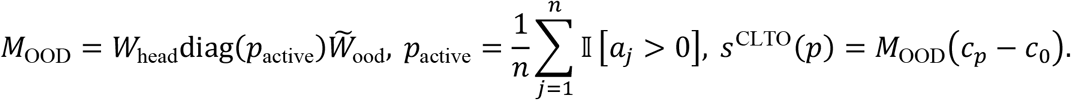

The module first triages perturbations by how far their cells move in the OOD subspace and then extracts the gene program decoded from that displacement.

### 3. Synthetic Multi-covariate Perturb-seq Data Simulation

We generated a synthetic multi-covariate single-cell RNA-seq dataset of 10,000 cells and 100 genes to benchmark the latent-factor disentanglement methods under controlled ground truth. The simulation was designed to simultaneously embed four sources of variation – cell differentiation state, cell-cycle progression, sequencing batch, and transcriptional perturbation – in a hierarchical variance structure (*State > Cell Cycle (CC) > Batch > Perturbation*), mimicking the relative magnitudes observed in real Perturb-seq experiments.

#### Cell Metadata

Each cell was assigned a continuous cell-cycle pseudotime coordinate *φ* drawn uniformly from [0,3) and a continuous differentiation pseudotime coordinate τ drawn uniformly from 0,2. Cells were labelled with a discrete cell-cycle phase (G1: *φ* ∈ [0,1), S: *φ* ∈ [1,2), G2M: *φ* ∈ [2,3)), yielding approximately equal phase proportions (G1: 33.8%, S: 33.9%, G2M: 32.3%). From *τ*, cells were assigned to three differentiation states via equal-width intervals (State1: *τ* < 2/3, State2: *τ* ∈ [2/3,4/3), State3: *τ* ≥ 4/3). Five experimental conditions (one non-targeting control, four perturbations) were represented with 2,000 cells each. Batch assignment was performed by stratified randomization within each of the 45 cross-tabulated groups of perturbation × CC phase × state, ensuring balanced representation across two batches.

#### Gene layout

Of the 100 simulated genes, 30 were designated cell-cycle markers (10 G1-phase markers: genes 050–059; 10 S-phase markers: genes 000–009; 10 G2M markers: genes 010– 019), 30 were differentiation state markers (10 per state, genes 020–049), and 40 were background genes (genes 060–099). The G1 marker set was a key design feature introduced to prevent G1-phase cells from collapsing to the expression mean in the absence of phase-specific signal.

#### Cell-cycle effect model

Cell-cycle marker expression was modelled using a symmetric half-sinusoidal bump function applied to all three marker groups. Each bump was parameterized by an onset position placed at the midpoint of the preceding phase zone, a width of 1.5 pseudotime units (equal to half the ring circumference), and a peak log fold change (LFC) of 3.5. Formally, for a cell at pseudotime *φ*, the CC contribution to log-expression for marker group *k* was:

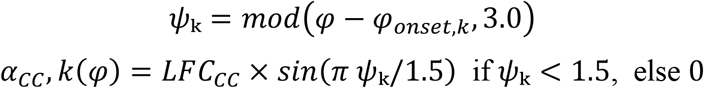

with onset positions *φ*_*onset,G*1_ = 2.5, *φ*_*onset,S*_ = 0.5, and *φ*_*onset,G*2*M*_ = 1.5. This design creates a 0.5-unit overlap between each adjacent pair of phases (*G*1 ∩ *S*: *φ* ∈ [0.5,1.0); *S* ∩ *G*2*M*: *φ* ∈ [1.5,2.0); *G*2*M* ∩ *G*1: *φ* ∈ [2.5,3.0)), during which two marker sets are simultaneously expressed at intermediate levels, reflecting the gradual transitions observed in biological cell-cycle progression. Phase marker sets are mutually exclusive at their respective peaks (each marker is exactly zero at the peaks of all other markers). The symmetric design ensures that all three phase transitions are equally represented in the expression manifold, producing a nearequilateral triangular arrangement of phase centroids in CC-gene PCA space (pairwise centroid distances: G1↔S = 5.58, S↔G2M = 5.142, G2M↔G1 = 5.08; symmetry ratio = 1.10).

#### Differentiation state effect model

Differentiation state was encoded using tent (triangular hat) basis functions *w*_1_(*τ*), *w*_2_(*τ*), *w*_3_(*τ*) that form a partition of unity over *τ* ∈ [0,2]. State marker genes received a log-mean contribution *βk*(*τ*) = *Σ*_j_*Akj* · *w*_j_(*τ*), where the coefficient matrix was *A* = [[+5, −1, −3], [−3, +5, −3], [−3, −1, +5]] (rows: states 1–3; columns: basis weights *w*_1_–*w*_3_). The peak LFC of +5.0 was selected to achieve Cell State *η*^2^ > 1.5 × Cell Cycle *η*^2^.

#### Cross-covariate coupling

To reflect the biological reality that state and CC axes are not fully orthogonal, a small cross-coupling term was included: state marker genes received an additional CC-dependent log-mean contribution weighted by a gene-specific coupling coefficient drawn from *Clip*(*N*(0.35,0.05), 0.2,0.5), and CC marker genes received a state-dependent contribution weighted by *Clip*(*N*(0.50,0.05), 0.3,0.7).

#### Batch effects

A per-gene batch shift *δ*^g^ was drawn from *N*(0,0.45). Batch-0 cells received +*δ*^g^ and Batch-1 cells received −*δ*^g^ for each gene, creating symmetric batch separation with no net effect on population mean. Library sizes were drawn from batch-specific lognormal distributions (Batch-0: mean ≈ 5,800 UMIs; Batch-1: mean ≈ 6,800 UMIs) and used to scale expected total counts per cell via a cell-specific normalization constant.

#### Perturbation effects

Perturbation *A* targeted 50 genes (20 sampled from the CC/state gene pool, 30 from the background pool) with effect sizes drawn from *N*(0,0.8). Perturbation *B* was constructed to be highly correlated with A: 45 of the 50 *A*-targeted genes were retained (90% overlap) with *B*-specific effects defined as 0.85 × *γ*_*A*_ + *N*(0,0.10), yielding a Pearson correlation *r*(*γ*_*A*_, *γ*_*B*_) = 0.993. Perturbation *C* targeted 18 genes drawn exclusively from genes not targeted by either *A* or *B* (0% overlap with *A* ∪ *B*), representing an orthogonal perturbation. Perturbation *D* added negligible genome-wide noise (*N*(0,0.05) across all genes) to serve as a near-null control. The non-targeting control (NT) carried no perturbation effect.

#### Count generation

Final expected log-means were assembled as:

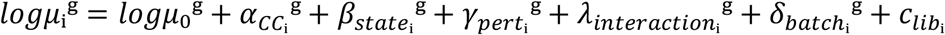

where 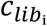 is a cell-specific scaling constant that normalises expected total counts to the sampled library size. Counts were sampled from a Zero-Inflated Negative Binomial (ZINB) model: Poisson rates λ_i_^g^ were first sampled from a Γ(*r*^g^, *µ*_i_^g^/*r*^g^) distribution (where *r*^g^ = 1/*BCV*^2^ encodes gene-level overdispersion), raw counts were drawn from *Poisson*(*λ*_i_^g^), and a gene- and expression-level-dependent dropout probability *π* = *sigmoid*(−1.5 − 0.8*log*(*µ* + 0.1)) was applied as a *Bernoulli* mask to introduce excess zeros. The resulting dataset had 41.2% sparsity and a mean of 6,263 UMIs per cell.

#### Variance decomposition

The relative contributions of each biological and technical covariate to transcriptome-wide variance were quantified using the mean per-gene *η*^2^ computed on *log*1*p*-normalized counts. The final variance hierarchy follows *Cell State* (19.6%) > *Cell Cycle* (10.9%) > *Batch* (4.6%) > *Perturbation* (3.6%), with a State-to-Cell Cycle ratio of 1.81× to achieve the situation where cell state should be the dominant source of variation while cell cycle remains clearly distinguishable above batch (**Fig. 2a** and **Extended Data Fig. 1**). All simulations were performed in Python 3.11 with a fixed random seed of 42 for reproducibility.

### 4. Benchmark

#### 4.1 Shared Training Protocol

To ensure comparability across models, a common training configuration was enforced for all benchmarked methods except for two standard approaches (COMBAT and LDA). All model-based methods were evaluated on the complete 10,000 cells × 100 genes synthetic dataset, using raw UMI counts for count-based probabilistic models and ln(CP10K + 1)-normalized expression for continuous matrix methods. To evaluate generalization to unseen cells under a data-sparse training regime (20% combined training and validation vs. 80% held-out test, 8,000 cells), the dataset was partitioned via stratified random splitting stratified jointly across the 10 perturbation × batch strata (5 perturbations × 2 batches), yielding a 2,000-cell training/validation pool (1,800 training and 200 validation cells for the 18%/2%/80% three-way split used by LinearSCVI, DrVI, ContrastiveVI, and MoCAVI, or 2,000 training and 8,000 validation/test cells passed via external indexing for PertVI and MrVI). PertVI and MrVI were trained for up to 2,000 epochs with early stopping (max_epochs = 2000, early_stopping = True, converging at 755 and 225 epochs, respectively), whereas LinearSCVI, DrVI, ContrastiveVI, and MoCAVI were trained for a fixed 200 epochs (max_epochs = 200, early_stopping = False), using minibatch size batch_size = 512, the Adam optimizer with learning rate 10^−3^, and two-layer 128-unit multi-layer perceptron (MLP) encoders and decoders on a single NVIDIA Ge-Force RTX 5090 GPU (32 GB VRAM; **Extended Data Fig. 2a**). No model received ground-truth cell_cycle_phase, cc_pseudotime, cell_state, or diff_pseudotime annotations during training. For uniform 2D visualization (**Fig. 2c**), *k*-nearest-neighbor graphs (scanpy.pp.neighbors) and Uniform Manifold Approximation and Projection (UMAP) embeddings (scanpy.tl.umap) were computed from each learned latent representation in Scanpy^38^.

#### 4.2 Benchmarked Approaches

##### Uncorrected

For the uncorrected baseline representation, raw UMI counts were library-size normalized, natural-log transformed (scanpy.pp.log1p), centered and scaled to unit variance per gene, and reduced via PCA without batch or covariate adjustment, followed by 15-nearest-neighbor graph construction and 2D UMAP projection (**Fig. 2c, d** and **Extended Data Fig. 2b**).

##### ComBat

Parametric empirical Bayes batch adjustment (scanpy.pp.combat) was applied to the ln(CP10K + 1)-normalized expression matrix specifying sequencing batch (Batch0 versus Batch1) as the batch key and perturbation as the preserved biological covariate, followed by unit-variance scaling, 30-component PCA, and 2D UMAP projection (**Fig. 2c, d** and **Extended Data Fig. 2b**).

##### LDA

Supervised multi-class Linear Discriminant Analysis was fitted on the top 30 uncorrected principal components using the 5-class perturbation identity as the supervision target (test_size = 0.20, random_state = 42), projecting all 10,000 cells onto the canonical discriminant axes prior to neighbor graph construction and 2D UMAP embedding (**Fig. 2c, d** and **Extended Data Fig. 2b**).

##### LinearSCVI

LinearSCVI^35^ was configured with a 30-dimensional latent space (n_latent = 30), layer = “counts”, batch_key = “batch”, a non-linear MLP encoder (dropout_rate = 0.1), and a linear factor-loading decoder under a Negative Binomial likelihood (gene_likelihood = “nb”, dispersion = “gene”), reconstructing log-normalized expression as 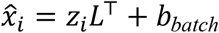, where *L* ∈ ℝ^100×30^ is the learned gene loading matrix (model.get_loadings()) and *b*_*batch*_ is a batch-specific intercept (**Extended Data Fig. 2a**). No perturbation labels were supplied during training. Analytical gene-level perturbation effect estimates (**Extended Data Fig. 2f, g**) were computed in the linear decoder’s log-normalized space as 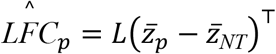, where 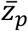 and 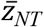 denote the mean latent representations of perturbation pand NT cells, respectively.

##### MrVI

Multi-resolution Variational Inference^41^ was configured on raw UMI counts (layer = “counts”) with batch_key = “batch” and sample_key = “perturbation”, parameterizing a 30-dimensional sample-unaware basal latent space (n_latent_u = 30) and a 30-dimensional sample-aware latent space (n_latent = 30). The model was trained with max_epochs = 2000, lr = 1e-3, n_epochs_kl_warmup = 40, batch_size = 512, and early stopping (early_stopping = True), followed by 30-nearest-neighbor graph construction (n_neighbors = 30) and 2D UMAP embedding (min_dist = 0.1; **Fig. 2c, d** and **Extended Data Fig. 2a, b**).

##### DrVI

Disentangled Representation Variational Inference^37^ was configured as a fully unsupervised variational autoencoder on raw UMI counts (layer = “counts”, batch_key = “batch”, with no perturbation supervision) using a 30-dimensional total latent space (n_latent = 30) partitioned into n_split_latent equal sub-blocks, each independently KL-regularized, with two-layer 128-unit MLP encoders and decoders (encoder_dims = [128, 128], decoder_dims = [128, 128]) and a Negative Binomial likelihood (gene_likelihood = “nb”). The split hyperparameter was swept over all integer divisors of 30(n_split_latent ∈ {1, 2, 3, 5, 6, 10, 30}; **Extended Data Fig. 2a**), where n_split_latent = 1 achieved the highest test-set perturbation classification accuracy (0.678). Targeted split-level experiments were conducted at n_split_latent = 3 (10 dimensions per split) and n_split_latent = 5 (6 dimensions per split) to test whether matching the number of latent splits to the number of ground-truth variance sources improved axis-specific disen-tanglement (**Fig. 2c, d** and **Extended Data Fig. 2d, e**).

##### ContrastiveVI

ContrastiveVI^32^ was configured on raw UMI counts (layer = “counts”, batch_key = “batch”) with 30-dimensional background (n_background_latent = 30) and salient (n_salient_latent = 30) latent spaces, two-layer 128-unit MLP encoders (n_hidden = 128, n_layers = 2, dropout_rate = 0.1), and a single joint MLP decoder under a Zero-Inflated Negative Binomial likelihood. Unperturbed NT cells (2,000 cells) were assigned as background_indices (whose salient posterior is architecturally constrained toward *N*(0, I) via per-cell KL divergence during training) and perturbed cells (Pert-A through Pert-D, 8,000 cells) as target_indices. An additional Wasserstein penalty (wasserstein_penalty) was swept over {0.0, 0.1, 0.5, 1.0} (**Extended Data Fig. 2a**), selecting Wasserstein penalty = 0.5 based on maximum test-set salient perturbation accuracy (0.675; **Fig. 2c, d**). Gene-level perturbation effect estimates (**Fig. 2g** and **Extended Data Fig. 2g**) were computed from 25 posterior samples of model.get_salient_normalized_expression(n_samples = 25) as 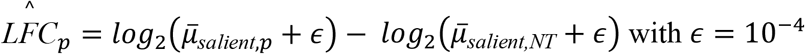 with ϵ = 10^−4^.

##### MoCAVI

Multi-modal Contrastive Analysis via Variational Inference^42^ was configured on raw UMI counts (layer = “counts”, batch_key = “batch”, labels_key = “perturbation”) with 30-dimensional background (n_background_latent = 30) and salient (n_salient_latent = 30) latent spaces, two-layer 128-unit MLP encoders and decoders (n_hidden = 128, n_layers_encoder = 2, n_layers_decoder = 2, dropout_rate = 0.05), n_regions = 0, and a Negative Binomial count likelihood (gene_likelihood = “nb”). A zero-valued dummy protein matrix was supplied to satisfy the multi-omic API without contributing to the training loss. Background (NT, 2,000 cells) and target (Pert-A through Pert-D, 8,000 cells) indices were intersected with the shared external_indexing partition by directly instantiating MoCAVIDataSplitter, TrainingPlan, and TrainRunner. The Wasserstein regularization weight was swept over {0.0, 0.1, 0.5, 1.0} (**Extended Data Fig. 2a**), selecting wasserstein_reg = 0.5 based on peak test-set salient perturbation classification accuracy (0.641; **Fig. 2c, d**). Gene-level perturbation effect estimates (**Fig. 2h** and **Extended Data Fig. 2g**) were obtained via model.synthetic_differential_abundance(n_samples = 25, delta = 0.25, fdr = 0.05), which samples from the salient posterior and decodes to gene space to compute log-fold changes relative to NT.

##### PertVI

Following the PertVI architecture, Split-FiLM transition module, and multi-objective governance described in the preceding **Methods** section, PertVI.setup_anndata was called on the synthetic dataset (10,000 cells × 100 genes) with layer = “counts”, batch_key = “batch”, sample_key = “perturbation”, treatment_mask_key = “is_treated” (0.0 for the 2,000 NT control cells and 1.0 for the 8,000 perturbed cells across Pert-A through Pert-D), and cycle_gene_mask_key = “is_cycle_gene” (registering the binary mask of the 30 ground-truth cell-cycle markers is_cc_marker). PertVI was instantiated with a 3-dimensional circular cellcycle subspace (n_latent_cycle = 3; *u*_*cycle*_, *z*_*cycle*_ ∈ ℝ^3^), a 30-dimensional macroscopic state subspace (n_latent_state = 30; *u*_*state*_, *z*_*state*_ ∈ ℝ^30^), and a 3-dimensional out-of-distribution anomaly subspace (n_latent_ood = 3; *v*_*ood*_ ∈ ℝ^3^), paired with 16-dimensional perturbation and batch lookup embeddings and two-layer 128-unit MLP encoders operating without access to perturbation or batch labels. For the primary benchmark configuration, PertVI was trained under the three-phase optimization schedule (warmup_epochs = 50, anneal_epochs = 100, n_epochs_kl_warmup = 40, max_epochs = 2000, batch_size = 512, lr = 1e-3, early_stopping = True, early_stopping_patience = 20) with regularization weights 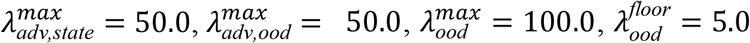, and *λ*_*soft*_ = 20.0, converging at 755 epochs with training and validation reconstruction losses of 252.7and 256.2 (**Extended Data Fig. 2a**). Posterior mean coordinates were extracted for all PertVI subspaces (*u*_*cycle*_, *u*_*state*_, *v*_*ood*_, *z*_*cycle*_, *z*_*state*_, and *z*_*final*_ = [*z*_*cycle*_, *z*_*state*_, *v*_*ood*_]) to construct neighbor graphs and 2D UMAP projections (**Fig. 2c–e** and **Extended Data Fig. 2b, c**).

#### 4.3 Downstream Analysis

##### Latent space disentanglement benchmarking

Across all benchmarked representations (Uncorrected, ComBat, LDA, LinearSCVI, MrVI *u* and *z*, DrVI at n_split_latent = 3 and n_split_latent = 5 plus individual split blocks, ContrastiveVI background and salient, MoCAVI background and salient, and PertVI *u*_cycle_, *u*_state_, *z*_cycle_, *z*_state_, and *z*_final_), disentanglement and covariate isolation were evaluated independently across the four ground-truth categorical variance sources (batch [2 classes], cell_cycle_phase [3 classes], cell_state [3 classes], and perturbation [5 classes]) using adversarial F1 (aF1) and Local Inverse Simpson’s Index (LISI)^116^ (**Fig. 2d** and **Extended Data Fig. 2b, d, e**). Specifically, aF1was computed as the uncalibrated 5-fold stratified cross-validation (StratifiedKFold, n_splits = 5, shuffle = True, random_state = 42) macro-averaged F1 score (f1_score, average = “macro”) of a *k*-nearest-neighbor classifier (KNeighborsClassifier, n_neighbors = 10, weights = “uniform”). For local neighborhood mixing, per-cell LISI scores were computed across all cells using harmonypy.compute_lisi (perplexity = 30) and summarized as log_10_(median LISI). For the targeted DrVI split-latent experiments (n_split_latent = 3 with three 10-dimensional blocks and n_split_latent = 5 with five 6-dimensional blocks), each contiguous split block was evaluated independently via aF1 and LISI across the four ground-truth covariates (**Extended Data Fig. 2d, e**).

##### Continuous cell-cycle ring and differentiation trajectory evaluation

To evaluate whether each method reconstructed the periodic ring topology of the cell cycle (G1–S–G2M) separately from the ordered differentiation continuum (State1–State2–State3), ground-truth continuous cell-cycle pseudotime (*φ*_*i*_ ∈ [0, 3)) and differentiation pseudotime (*τ*_*i*_ ∈ [0, 2]) were overlaid onto each method’s UMAP embeddings (**Fig. 2c**). In addition, diffusion-based pseudotime (Palantir^117^ initialized from the State1 origin) was inferred on each model’s baseline or state manifold and compared against ground-truth *τ*_*i*_ via Spearman rank correlation (**Fig. 2e** and **Extended Data Fig. 2c**).

##### Gene-level perturbation effect estimation benchmarking

Gene-level perturbation effect inference was benchmarked across all four generative methods equipped with an analytical or generative gene-space decoder (LinearSCVI, ContrastiveVI, MoCAVI, and PertVI) alongside a standard observational differential expression baseline (Standard DE, computed via two-sided Wilcoxon rank-sum testing in scanpy.tl.rank_genes_groups between each perturbation condition and NT on ln(CP10K + 1)-normalized expression; **Fig. 2f**). For PertVI, gene-level perturbation momentum scores were computed for each condition *p* ∈ {Pert-A, Pert-B, Pert-C, Pert-D} relative to NT via Global Path-integrated Momentum Score in the preceding **Methods** section. For each of the five methods (Standard DE, LinearSCVI, ContrastiveVI, MoCAVI, and PertVI), the inferred 100-gene effect vectors were compared against the known ground-truth perturbation log-fold-change vectors (*γ*_Pert-A_, *γ*_Pert-B_, *γ*_Pert-C_, *γ*_Pert-D_) across all *G* = 100 genes by computing the 4 × 4 matrix of pairwise Pearson correlation coefficients, where diagonal entries quantify condition-specific ground-truth recovery (Pert-A, Pert-B, Pert-C, and Pert-D) and off-diagonal entries diagnose cross-condition signal leakage (**Fig. 2f–i** and **Extended Data Fig. 2f**). Finally, to evaluate whether each model preserved the ground-truth relational structure among the four perturbations, 4 × 4 within-method pairwise Pearson correlation matrices were computed among the inferred 100-gene perturbation effect vectors of Pert-A, Pert-B, Pert-C, and Pert-D for LinearSCVI, ContrastiveVI, MoCAVI, and PertVI (**Extended Data Fig. 2g**).

### 5. Modeling on CD8+ T Cell Data

#### 5.1 Data collection, subset, and preprocess

*Data collection*. Single-cell RNA-sequencing (scRNA-seq) profiles from a high-throughput chemical perturbation screen in primary human CD8+ T cells were obtained from GSE306429. In the experimental assay (**Fig. 3a**), purified peripheral blood mononuclear cell (PBMC)-derived naive CD8+ T cells from two healthy blood donors (Donor 1 and Donor 2, serving as two biological replicates *b*_*n*_ ∈ {1, 2}) were pre-incubated for 24h with one of 62 structurally diverse small-molecule compounds at 1*µ*Mor with dimethyl sulfoxide (DMSO) vehicle control, followed by 72h of TCR stimulation by anti-CD3/CD28-coated beads and recombinant human IL-2 to induce clonal proliferation and effector differentiation. The resulting cohort comprises *N* = 62,207 quality-controlled single CD8+ T cells (34,548 cells from Donor 1 and 27,659 cells from Donor 2) spanning 63 experimental conditions: 6,472 unperturbed DMSO vehicle control cells and 55,735 compound-treated cells across the 62 small-molecule perturbations.

##### Preprocess

For generative latent manifold modeling and cross-method benchmarking, we selected the top *G* = 4,000 donor-aware highly variable genes (HVGs) from raw UMI counts using the Seurat v3 variance-stabilizing transformation (scanpy.pp.highly_variable_genes, flavor = “seurat_v3”, n_top_genes = 4000). This 4,000-HVG feature set retained 68 canonical Sphase and G2/M-phase cell-cycle marker genes from the 97-gene reference cell-cycle signature^18^. Conventional discrete cell-cycle phase annotations (G1, S, and G2M) and continuous S score and G2M score values were computed across all 62,207 cells using scanpy.tl.score_genes_cell_cycle.

#### 5.2 Model training

##### Uncorrected

For the uncorrected baseline representation (“Orig.”; **Extended Data Fig. 33, c**), library-size normalized and l*og*1*P*-transformed expression values across 4,000 HVGs were centered and scaled to unit variance per gene (*Z*_*ng*_ = (*Y*_*ng*_ − *µ*_*g*_)/*σ*_*g*_) without batch or covariate regression. Principal Component Analysis (PCA) was performed (scanpy.tl.pca, n_comps = 50) to extract the top 50 principal components, which were used to construct 15-nearest-neighbor Euclidean graph (scanpy.pp.neighbors, n_neighbors=15) and 2D Uniform Manifold Approximation and Projection (UMAP) embedding in Scanpy.

##### ComBat

Linear empirical Bayes batch adjustment was performed on the l*og*1*P* expression matrix in HVG space using ComBat (scanpy.pp.combat), specifying biological replicate *b*_*n*_ ∈ {1, 2} as the batch key and perturbation *p*_*n*_ ∈ {0, 1, …, 62} as covariate (**Extended Data Fig. 33, c**). The resulting batch-adjusted expression matrix was centered, scaled to unit variance, reduced to its top 50 principal components (n_comps=50), and embedded into a 2D UMAP representation via a 15-nearest-neighbor graph.

##### LDA

Supervised multi-class Linear Discriminant Analysis was applied to the top 50 uncorrected principal components (n_pcs = 50) using perturbation identity (*p*_*n*_ ∈ {0, 1, …, 62}) as the target supervision label (**Extended Data Fig. 33, c**). The discriminant projection matrix *W*_LDA_ ∈ ℝ^50×(*C*−1)^ maximizing Fisher’s criterion det(*W*^⊤^*S*_*B*_*W*)/det(*W*^⊤^*S*_*W*_*W*)between the between-class scatter matrix 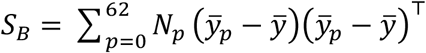 and the within-class scatter matrix 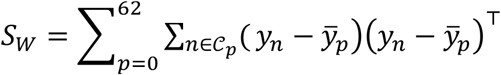 was fitted on a stratified 80% training split. The trained projection projected all 62,207 cells into the 63-dimensional discriminant sub-space, followed by 15-nearest-neighbor graph construction and 2D UMAP projection.

##### MrVI

MrVI was trained on the 4,000-HVG raw UMI count matrix with biological replicate (batch_key = “replicate”) registered as the technical nuisance batch covariate and compound identity (sample_key = “compound_name”) registered as the target sample covariate (**Extended Data Fig. 33, c**). Following training under learning rate at 1e-4, minibatch size 512, and 2000 max epochs (early-stop enabled), the sample-unaware basal latent representation *u* and the sample-aware perturbation latent representation *z* were obtained across all 62,207 cells and used to construct 15-nearest-neighbor graphs and 2D UMAP embeddings.

##### MoCAVI

MoCAVI was trained on the 4,000-HVG raw UMI count matrix by partitioning the dataset into an unperturbed background reference cohort (6,472 DMSO control cells) and a perturbed target cohort (55,735 compound-treated cells), with biological replicate supplied as the categorical batch covariate (**Extended Data Fig. 33, c**). Both the shared background encoder (base) and the perturbation-specific salient encoder were parameterized as two-layer 128-unit multi-layer perceptrons mapping onto 30-dimensional latent spaces under Wasserstein independence regularization (*λ*_wasserstein_ = 0.5). From the trained model, 30-dimensional background and salient embeddings were extracted to compute 15-nearest-neighbor graphs and 2D UMAP projections. Compound-level MoCAVI salient displacement magnitudes (**Fig. 3k**) were quantified as the Euclidean (*L*_2_) distance between each compound’s mean salient latent centroid and the DMSO control salient centroid in ℝ^30^.

##### PertVI

PertVI was trained on the 4,000-HVG raw UMI count matrix with biological replicate (batch_key = “replicate”), chemical perturbation identity (sample_key = “compound_name”), and the binary treatment indicator (treatment_mask_key = “is_treated”, set to 0 for the 6,472 DMSO control cells and 1 for the 55,735 compound-treated cells). The model was instantiated with a 3-dimensional basal and perturbation-modulated cell-cycle subspace (*u*_cycle_, *z*_cycle_ ∈ ℝ^3^), a 30-dimensional basal and perturbation-modulated developmental state subspace (*u*_state_, *z*_state_ ∈ ℝ^30^), and a 5-dimensional out-of-distribution anomaly subspace (*v*_ood_ ∈ ℝ^5^, anchored at the origin in DMSO control cells via binary treatment indicator). Subspace disentanglement was achieved through regularization weights at *λ*_adv,state_ = 100.0, *λ*_adv,ood_ = 50.0, *λ*_ood_ = 50.0 (with a minimum floor of 20.0), and soft cell-cycle gradient-starvation penalty *λ*_soft_ = 50.0. Using an 80% training (49,765 cells) and 20% validation (12,442 cells) split stratified jointly by compound and biological replicate, PertVI was trained for 600 epochs (max_epochs = 2000, early-stop enabled) using the Adam optimizer (learning rate 10^−4^, minibatch size 512) under a Negative Binomial count likelihood (**Extended Data Fig. 33**). After training, embeddings were extracted across all latent representations *(u*_*cycle*_, *u*_*state*_, *z*_*cycle*_, *z*_*state*_, *v*_*ood*_, and the concatenated *z*_*final*_ = [*z*_*cycle*_, *z*_*state*_, *v*_*ood*_]). Batch-harmonized denoised log1P-expression profiles (X_pertvi_log1p) were decoded and standardized to reference donor D1. UMAP projections were computed for each latent space (**Fig. 3b, f, j, m and Extended Data Fig. 33**).

#### 5.3 Downstream analyses

##### Latent space integration and disentanglement benchmarking

We quantitatively benchmarked 12 latent embeddings (uncorrected Orig., ComBat, LDA, MrVI *u*, MrVI *z*, MoCAVI base, MoCAVI salient, and PertVI *u*_cycle_, *u*_state_, *z*_cycle_, *z*_state_, and *z*_final_) across four biological and technical annotation axes: biological donor replicate (Donor, 2 classes), canonical discrete cellcycle phase (CC Phase, 3 classes), CD8+ T-cell developmental state (Cell State, 5 classes), and chemical perturbation identity (Perturbation, 63 classes; **Extended Data Fig. 33, c**). Clustering separation and label conservation were evaluated via the adjusted F1 score (aF1) using 5-fold stratified cross-validation with a 5-nearest-neighbor classifier. Within each fold, the observed macro-averaged F1 score (F1_obs_) was calibrated against an empirical label-permutation null baseline (F1_null_) via aF1 = max(0, (F1_obs_ − F1_null_)/(1 − F1_null_)) and averaged across the five folds. Local neighborhood mixing was quantified across all 62,207 cells using the Local Inverse Simpson’s Index (harmonypy.compute_lisi, perplexity = 30) and normalized on a logarithmic scale.

##### Continuous cell-cycle ring reconstruction

On the 2D circular projection of *u*_cycle_, continuous angular cell-cycle progression pseudotime (CCPT, *t* ∈ [0, 1]) was oriented from quiescent G1 entry (*t* = 0) through DNA synthesis (S) and mitosis (G2/M, *t* → 1) and discretized into 10 uniform decile bins (*B*_*m*_ = [*m*,/10(*m* + 1)/10) for *m* ∈ {0, …, 9}, corresponding to 0%–10% through 90%–100%; **Fig. 3b, e**). Based on inflection boundaries of denoised canonical cellcycle marker waves, the 10 decile bins were grouped into five sequential sub-phases: earlyG1/G1 (Bins 0–1), early/mid-S (Bins 2–3), late-S (Bins 4–5), G2 phase (Bins 6–7), and late-M (Bins 8–9; **Fig. 3e**). Next, we classified compound-induced cell-cycle remodeling using a two-stage unsupervised pipeline (**Fig. 3c–e**). In Stage 1, each compound’s 10-bin empirical cumulative distribution function *F*_*k*_(*t*) was compared against the pooled DMSO control distribution *F* (*t*) using the 1D Wasserstein distance 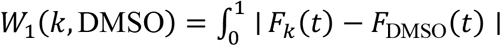 *dt*(scipy.stats.wasserstein_distance) alongside the maximum absolute single-bin probability shift max_*m*_ ∣ Δ*p*_*k*_(*B*_*m*_) ∣, where Δ*p*_*k*_(*B*_*m*_) = *P*(*B*_*m*_ ∣ *k*) − *P*(*B*_*m*_ ∣ DMSO). Calibrating against empirical inter-replicate DMSO variation established the activity cutoff *W*_1_ ≥ 0.015, partitioning the 63 conditions into 29 active cell-cycle modulators and 34 non-cell-cycle-shifting (T-cell-sparing) conditions (33 compounds plus DMSO; **Fig. 3c**). In Stage 2, the 10-bin differential abundance vectors Δ*p*_*k*_ ∈ ℝ^10^ of the 29 active modulators were row-normalized to unit *L*_2_ norm 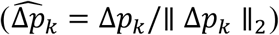 and clustered via spherical *k*-means (*k* = 4, grouping pure G1 arrest and dual G1 plus late-M cytokinesis retention into G1-arrest) and phase-peak localization into G1-arrest (*n* = 10, peaking in Bins 0–1), S-crisis (*n* = 7, peaking in Bins 2– 5), and G2/M-arrest (*n* = 12, peaking in Bins 6–9; **Fig. 3d, e**). Differential 2D Gaussian kernel density estimation contours (ΔDensity versus DMSO on *u*_cycle_) were computed for each arrest regime (**Fig. 3d**). Cross-referencing these primary CD8+ T-cell phenotypes against published arrest profiles in solid-tumor and leukemic cell lines partitioned all 63 conditions into four immuno-oncology classes (**Fig. 3c**): Conserved (*n* = 21 active in both tumor lines and primary T cells), Tumor-active-only / T-cell-sparing (*n* = 10 canonical phase-specific cell-cycle blockers), Novel in T cells (*n* = 8 *de novo* T-cell proliferation remodelers), and Neutral (*n* = 24 active in neither tumor lines or primary T cells).

##### Continuous Developmental trajectory reconstruction

To define canonical CD8+ T-cell developmental subsets along the 30-dimensional perturbation-agnostic basal state manifold (*u*_state_), we constructed a 15-nearest-neighbor graph and performed unsupervised Leiden community detection (scanpy.tl.leiden, resolution = 0.5) across all *N* = 62,207 quality-controlled cells. The resulting five communities (Clusters 0–4) were annotated into sequential developmental subsets based on PertVI-denoised expression (X_pertvi_log1p) of 14 canonical naive, proliferative, memory-precursor, cytolytic, and exhaustion marker genes (**Fig. 3f and Extended Data Fig. 44**): quiescent naive T cells (Tn, Cluster 1, *n* = 16,060, defined by high expression of naive homing and stemness markers *TCF7, LEF1, SELL*, and *CCR7*), early proliferating blasts (Tn.prolif, Cluster 0, *n* = 18,796, defined by co-expression of naive markers with cellcycle progression genes *TOP2A* and *PCNA*), intermediate memory precursors (T.eff.IL7R+, Cluster 4, *n* = 5,076, defined by induction of GZMB and peak *IL7R* expression), cytolytic effectors (T.eff, Cluster 2, *n* = 15,505, defined by induction of *GZMB, IFNG*, and *NKG7*), and terminally differentiated effectors (T.eff.Term, Cluster 3, *n* = 6,770, defined by maximal co-expression of *TOX, LAG3*, and *PRDM1* alongside *GZMB* and *IFNG*). Continuous developmental progression along the *u*_state_ was parameterized using Diffusion Maps (scanpy.tl.diffmap, n_comps = 15) and Diffusion Pseudotime (*τ* ∈ [0, 1], scanpy.tl.dpt, n_dcs = 10). Cell exhibiting maximal denoised *TCF7* expression within the quiescence subset was set as root cell for trajectory inference.

##### Characterization of state-shifting archetypes

The continuous *τ* ∈ [0, 1] coordinate was partitioned into 10 equal-width decile bins. For each compound *k*, we computed the per-bin probability mass difference vector Δ*P*_*k*_ = [Δ, PMF_*k*_, (*b*_0_) … ΔPMF_*k*_(*b*_9_)]^⊤^ relative to DMSO 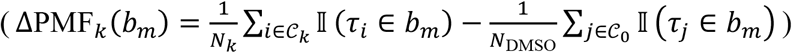 and the 5-state compositional log-fold-change. To classify perturbation-induced state shifts, we implemented a twostep classification framework (**Extended Data Fig. 44, c**): in Step 1, Ward’s minimum-variance hierarchical linkage on pairwise Euclidean distances ∥ Δ*P*_*j*_ − Δ*P*_*k*_ ∥_2_ across the 10 DPT deciles defined three candidate shift topologies; in Step 2, compounds were required to pass either a Benjamini-Hochberg false discovery rate (FDR)-corrected two-sample KolmogorovSmirnov test *q*_KS_(*k*) < 0.05 on the unbinned single-cell *τ* distribution against DMSO or a 2 × 5 discrete state contingency Pearson chi-square test 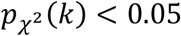 (df = 4) paired with a peak discrete state enrichment magnitude max_*s*∈*S*_ ∣ log_2_FC_*k*_(*s*) ∣≥ 0.20. Compounds failing both criteria (*n* = 34) were assigned to “Non-shifting”.

##### Cross-manifold comparison

Cross-manifold statistical coupling between the four *u*_cycle_ proliferation classes (G1-arrest, S-crisis, G2/M-arrest, and No-CC-shift) and the four *u*_state_ differentiation archetypes (C1, C2, C3, and non-shifting) across all 62 compounds (**Fig. 3i**) was evaluated on the 4 × 4 contingency table via Pearson’s chi-square test and standardized cell residuals 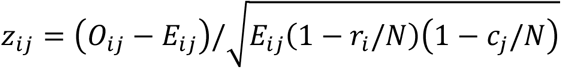.

##### Multi-resolution Mechanism of Action enrichment

To evaluate how pharmacological mechanism of action (MoA) maps onto T-cell differentiation archetypes (**Extended Data Fig. 44 and Extended Data Fig. 5**), the 62 compounds were annotated across three hierarchical phar-macological resolutions: eight high-level MoA categories (Category 1: PI3K/mTOR and intracellular kinases, *n* = 11; Category 2: RTK/TKI and JAK/SYK kinases, *n* = 12; Category 3: cell-cycle and mitotic kinases, *n* = 6; Category 4: epigenetic and chromatin modifiers, *n* = 8; Category 5: proteostasis, chaperones, and transport, *n* = 7; Category 6: DNA damage, p53, and apoptosis, *n* = 6; Category 7: ion pumps, cAMP, and metabolism, *n* = 5; and Category 8: nuclear receptors and transcription, *n* = 7, alongside a 7-category sensitivity merge combining Categories 7 and 8 into *n* = 12), 32 fine-grained drug classes (*M* = 12 multi-drug classes with *n* ≥ 2), and a two-way macro-class partition comparing all kinase inhibitors (Categories 1–3, *n* = 29) against non-kinase modulators (Categories 4–8, *n* = 33). Global omnibus contingency tests were evaluated using asymptotic Pearson chi-square tests, 100,000-iteration Monte Carlo permutation chi-square tests (random seed 42), and exact Freeman-Halton tests, complemented by non-parametric Kruskal-Wallis *H* tests on continuous shift magnitudes (**Extended Data Fig. 5a**). To determine whether apparent MoA insensitivity reflected lineage-specific target absence in primary human CD8+ T cells (**Extended Data Fig. 5c, d**), each compound *k* was mapped to its canonical biochemical target genes *T*_canon_(*k*) and, where established by competitive kinome binding and chemoproteomic profiling studies, documented nanomolar off-target T-cell kinase genes *T*_off_(*k*) (specifically neratinib cross-inhibiting *LCK, MAP4K1*/HPK1, and *ITK* alongside canonical *ERBB2*/*EGFR*, and NVP-BHG 712 cross-inhibiting *LCK* and *RAF1*/c-Raf alongside canonical *EPHB4*). Across the *N*_DMSO_ = 6,472 control CD8+ T cells in the unfiltered 25,078-gene transcriptome, single-cell target detection frequency was computed as 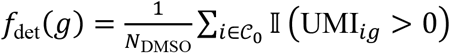 and compound-level effective target availability was defined as 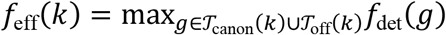 (**Extended Data Fig. 5c**). Differences in target expression between intracellular kinases and receptor/non-receptor tyrosine kinases were evaluated via two-sided Mann-Whitney *U* tests, and the single-cell missing-target gate was tested via Haldane-Anscombe odds ratios and Boschloo’s exact tests at strict (*f*_eff_(*k*) < 5%, *n* = 8 target-absent versus *n* = 54 target-expressed compounds) and trace (*f*_eff_(*k*) < 15%, *n* = 9 versus *n* = 53) cutoffs, alongside an unadjusted canonical-target-only control (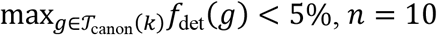 versus *n* = 52; **Extended Data Fig. 5d**).

##### Global transcriptomic Momentum and state-conditioned Counterfactual Shifts

To separate bulk, trajectory-averaged transcriptional responses from stage-restricted lineage-driving impulses on *z*_state_ (**Fig. 3j–l and Extended Data Fig. 6a**), we used PertVI’s Split-FiLM decoder (holding *v*_ood_ = 0) to compute two counterfactual vector fields across the *G* = 4,000 HVGs for each compound *k*: global transcriptomic velocity (Momentum_*k*_ ∈ ℝ^4000^, computed via trajectory-basic as the net directional shift across the basal manifold relative to DMSO, with scalar magnitude ∥ Momentum_*k*_ ∥_2_) and state-conditioned Counterfactual Shifts (CSS_*k,s*_ ∈ ℝ^4000^, computed with inject_film=True independently within each of the five developmental states). Comparing global ∥ Momentum_*k*_ ∥_2_ against phenotypic state-shift activity on *u*_state_ partitioned the 62 compounds into four decoupling quadrants (**Extended Data Fig. 6**): Q1 (high-Momentum state shifters, *n* = 15), Q2 (low-Momentum lineage-gating catalysts, *n* = 13), Q3 (high-Momentum buffered non-shifters, *n* = 7), and Q4 (silent non-shifters, *n* = 27).

##### Functional analysis

Across all developmental spaces and all 62 compounds, transcription factor (TF) regulon activities were inferred using the univariate linear model (ULM) in decoupleR^118^ against the curated human CollecTRI signed transcriptional regulatory network^119^ filtered to regulons with at least 5 expressed target genes among the 4,000 HVGs (518 tested TFs). For each compound *k*, space *s*, and transcription factor tf, ULM fitted the linear regression *Y*_*g,k,s*_ = *β*_0_ + *β*_tf_*M*_*g*,tf_ + ϵ_*g*_ and returned the signed studentized *t*-statistic 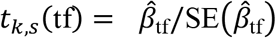. In parallel, biological process enrichment across the six spaces was computed via pre-ranked Gene Set Enrichment Analysis^120^ against Gene Ontology Biological Process (GOBP) gene sets to obtain normalized enrichment scores (NES_*k,s*_(*p*)). To identify consensus drivers within each archetype (C1–C3) and across the eight Archetype ×MoA sub-modules, compound-level directional votes 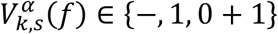 across four empirical quantile cutoffs *α* ∈ {0.01, 0.025, 0.05, 0.10} were integrated via Robust Rank Aggregation (RRA)^121^ with Benjamini-Hochberg FDR control (*q*_shared_ < 0.05), summarized by the net vote fraction 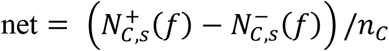, and tested for between-archetype specificity via one-vs-rest Mann-Whitney *U*tests (*p*_sp_ and BH-adjusted *q*_sp_).

##### Off-manifold anomaly quantification and functional toxicity classification

Within the 5-dimensional anomaly subspace *v*_ood_ ∈ ℝ^5^, each compound’s scalar off-manifold displacement was quantified as the Euclidean distance between its treatment centroid and the DMSO control centroid, 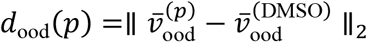 (**Fig. 3m, n**). Across all 62 compounds, *d* was calibrated against three statistical thresholds (**Extended Data Fig. 7a**): an empirical 99.9th percentile DMSO bootstrap noise floor (*d*_null_ = 0.0550, estimated from 10,000 resamples of control cells), a bimodal two-component Gaussian Mixture Model (on-manifold mean *µ*_1_ = 0.0885 versus off-manifold mean *µ*_2_ = 0.4082) that established the operational threshold *d*_ood_ ≥ 0.30 separating *N* = 14 off-manifold stress-inducing compounds from *N* = 48 on-manifold compounds, and a Tukey upper outlier fence (Q3 + 1.5 × IQR = 0.5623) isolating five extreme off-manifold outliers (ruxolitinib 1.156, serdemetan 0.719, BMS-536924 0.677, thapsigargin 0.670, and sapanisertib 0.662). Enrichment of off-manifold displacement across the four differentiation archetypes and across MoA classes was evaluated using two-sided Fisher’s exact tests with Benjamini-Hochberg FDR correction (**Fig. 3n and Extended Data Fig. 7b**). To extract pure off-manifold gene signatures free of developmental state confounding, we used PertVI’s Marginalized Counterfactual Operator (MCO), which integrates the *v*_ood_ counterfactual intervention exclusively across the fixed, unperturbed DMSO vehicle control cell distribution. Gene-level MCO log-fold-change signatures were then profiled across 18 canonical oxidative, xenobiotic, endoplasmic reticulum stress, and interferon-stimulated cytolytic marker genes (**Extended Data Fig. 7c**). Pre-ranked Gene Set Enrichment Analysis (gseapy.prerank, 1,000 permutations) was executed on the 4,000-HVG MCO vector of each compound against all 50 MSigDB Hallmark gene sets. Two-sided Mann-Whitney *U*tests with Benjamini-Hochberg FDR control (FDR < 0.05) comparing Hallmark NES distributions between the *N* = 14 off-manifold compounds and *N* = 48 on-manifold compounds identified eight core differentially regulated stress and cell-cycle pathways (**Fig. 3o**), and unsupervised Ward hierarchical clustering across the 48 Hallmark pathways stratified the 14 off-manifold compounds into six mechanism-specific toxicity classes (Classes I–VI) across five functional pathway programs (**Extended Data Fig. 7e, f**).

### 6. Modeling on CD4+ T Cell Data

#### 6.1 Data collection, subset, and preprocess

##### Pilot dataset creation

Single-cell CRISPRi Perturb-seq data and accompanying arrayed CRIS-PRi bulk RNA-seq data from primary human CD4+ T cells were obtained from a published genome-scale perturbation atlas^15^ (GSE314342). The parent Perturb-seq atlas profiles 21.99 million single-guide primary human CD4+ T cells targeting 12,779 expressed genes (26,504 sgRNAs including 5% non-targeting controls) across four healthy blood donors (D1–D4) and three T-cell receptor (TCR; anti-CD3/CD28/CD2) stimulation conditions: unstimulated resting baseline (Rest), acute activation at 8h post-stimulation (Stim 8hr), and late blastogenic effector maturation at 48h post-stimulation (Stim 48hr), defining 111 co-functional perturbation clusters across conditions. To construct a balanced multi-condition benchmark dataset, we first filtered out low-quality droplets and multi-guide transductions (~low_quality and guide_id != ‘multi-guide’) and applied a water-filling stratified sampling algorithm across donors and conditions to assemble 120,545 high-quality single-guide cells spanning Rest (42,344 cells), Stim 8hr (40,272 cells), and Stim 48hr (37,929 cells) across donors D1 (25,751 cells), D2 (41,133 cells), D3 (26,491 cells), and D4 (27,170 cells). Overall, the pilot dataset comprises 82 perturbation labels (81 gene-targeting knockdowns across 114,521 cells plus non-targeting controls across 6,024 cells). Specifically, 61 curated T-cell activation regulators (77,003 cells) were selected from the original study’s Figure 3 HDBSCAN T-cell activation clusters (Cluster 21 at 8h, Cluster 22 at 8h and 48h, and Cluster 99 at 48h). Based on their annotated active timepoints, these regulators were stratified into early-(active at 8h in Cluster 21, *n* = 12; 19,248 cells), general-(shared across 8h in Clusters 21 and 22 and 48h in Cluster 99, *n* = 29; 34,081 cells), and late-(active at 48h in Cluster 99, *n* = 20; 23,674 cells) stage regulators spanning seven functional categories: T-cell activation, vesicle trafficking and cytoskeleton dynamics, transcriptional regulators, cell cycle and genome integrity, metabolism, calcium ho-meostasis, and signaling. To establish an empirical null baseline alongside the curated regulators, 20 non-essential negative control genes (NCGs; 37,518 cells) were randomly sampled from perturbed genes falling strictly outside all co-functional HDBSCAN clusters in the parent atlas, and 6,024 non-targeting control (NTC) cells were uniformly sampled across the 12 donor-condition pools (502 cells per pool, 2,008 cells per condition, and 1,506 cells per donor) to match the 5.00% NTC proportion of the parent library.

##### Preprocess

We identified the top 10,000 donor-aware highly variable genes (HVGs) directly from raw UMI counts using the Seurat v3 variance-stabilizing transformation (flavor=“seurat_v3”, n_top_genes=10000, batch_key=“donor”) implemented in Scanpy and formed their union with 94 expressed canonical cell-cycle marker genes (42 S-phase and 52 G2/M-phase markers). This union yielded 10,004 expressed genes comprising 9,910 non-cycle HVGs and 94 canonical cell-cycle markers. Raw UMI counts were preserved for modeling. To benchmark canonical cell-cycle phase calling, phase scoring (scanpy.tl.score_genes_cell_cycle) was executed independently within each stimulation condition on library size-normalized and log-transformed expression (X_log1p) to assign per-cell S score, G2M score, and discrete phase labels (G1, S, and G2M).

#### 6.2 Model training

*PertVI*. PertVI was configured on the CD4+ T-cell dataset (120,545 cells ×10,004 genes) with a 3-dimensional basal cell-cycle subspace (*u*_cycle_, *z*_cycle_ ∈ ℝ^3^), a 30-dimensional basal cell-state subspace (*u*_state_, *z*_state_ ∈ ℝ^30^), and a 5-dimensional out-of-distribution subspace (*v*_ood_ ∈ ℝ^5^, configured with encoder dropout rate 0.3 and a baseline treatment mask set to 0.0 for resting NTC cells and 1.0 otherwise). Subspace independence was enforced during training using cross-subspace orthogonality regularization between *u*_cycle_ and *u*_state_ (*λ*_adv,state_ = 100.0, *λ*_adv,ood_ = 20.0) and an *L*_1_ origin-anchoring penalty on *v*_ood_(*λ*_ood_ = 20.0, floor 5.0). Using an 80% training (96,436 cells) and 20% validation (24,109 cells) split stratified jointly by perturbation label and donor (random seed 42), the model was trained on raw UMI counts for 2000 epochs (early-stop enabled) using the Adam optimizer (learning rate 10^−4^, minibatch size 512, 40-epoch KL warmup, and 50-epoch adversarial warmup followed by 100-epoch annealing) under a Negative Binomial count likelihood, with convergence monitored across total evidence lower bound (ELBO), Negative Binomial reconstruction error, and Kullback-Leibler (KL) divergence (**Extended Data Fig. 8b**). Batch-corrected denoised expression (X_pertvi_log1p) was computed by decoding normalized expression standardized to reference donor D2 (transform_batch=“D2”, scaled to total sum at 10^4^) followed by log(1 + *x*) transformation. Uniform Manifold Approximation and Projection (UMAP) embeddings were computed in Scanpy (n_neighbors=15) across uncorrected PCA coordinates and each of PertVI’s six latent representations (*u*_cycle_, *u*_state_, *z*_cycle_, *z*_state_, *v*_ood_, *z*_final_; **Fig. 4b and Extended Data Fig. 8c**).

##### scVI

To benchmark PertVI against the single-cell generative embedding architecture used in the original study^15^, we trained a dimensionality-matched, unconstrained 30-dimensional scVI model^30^ (30D scVI) using scvi-tools^62^ on the identical 120,545 cells and 10,004 genes (**Extended Data Fig. 11a**). Matching the latent embedding protocol of the original study, the scVI model was configured on raw UMI counts with donor batch conditioning (batch_key=“donor”, encode_covariates=True) and was trained for 2000 epochs (early-stop enabled) with a minibatch size of 2,048 and learning rate at 10^−4^. All downstream visualization followed the same settings used for PertVI.

#### 6.3 Downstream analyses

##### Cell-cycle ring reconstruction and phase-zone differential abundance

Within the 3-dimensional basal cell-cycle subspace (*u*_cycle_), where the first two principal components captured 99.94% of total variance as a planar ring (**Extended Data Fig. 8d**), continuous mitotic progression was parameterized along a 6-waypoint polar geodesic pseudotime (*τ*_cycle_ ∈ [0, 1]; **Fig. 4c, d and Extended Data Fig. 8e**). Six ordered biological waypoints (*W*_0_ to *W*_5_) were defined in 3D *u*_cycle_ space from the quiescent Rest G1 centroid (*W*_0_), the activated Stim 8hr G1 centroid (*W*_1_), early and late S-phase centroids (*W*_2_, *W*_3_), and early and mitotic-peak G2/M centroids (*W*_4_, *W*_5_); cells were orthogonally projected onto the piecewise-linear polygonal curve *W*_0_ → *W*_1_ → *W*_2_ → *W*_3_ → *W*_4_ → *W*_5_ with cumulative arc length quantile-scaled to [0, 1]. Based on inflection boundaries of canonical phase-marker expression, *τ*_cycle_ was partitioned into five se-quential mitotic phase zones: basal quiescence (C1: G0/G1, [0.00, 0.22)), restriction-point licensing (C2: Activated G1, [0.22, 0.40)), early DNA synthesis (C3: Early S, [0.40, 0.58)), late DNA replication (C4: Late S, [0.58, 0.76)), and mitotic division (C5: G2/M, [0.76, 1.00]). Denoised expression profiles (X_pertvi_log1p) of canonical cell-cycle markers were smoothed across *τ*_cycle_ bins and grouped into five sequential transcriptional waves spanning G0/early-G1 quiescence (W1), G1/S restriction-point licensing (W2), S-phase DNA replication (W3), G2/M checkpoint control (W4), and cytokinesis exit (W5). To verify that *u*_cycle_ provides a perturbation-agnostic mitotic coordinate, trajectory conservation was quantified via Pearson correlation (*r*) between unperturbed NTC controls (*n* = 6,024) and all 120,545 cells at both the 5-wave composite level and the individual marker level (**Extended Data Fig. 8f, g**). For each perturbation and stimulation condition, within-condition phase-zone differential abundance (log_2_DA) across zones C1–C5 was computed as the log_2_fold ratio of perturbed cell proportions to condition-matched NTC cell proportions within each zone (with pseudocount *ε* = 0.005), with statistical significance assessed via two-sided Fisher’s exact tests and corrected for multiple hypothesis testing using the Benjamini-Hochberg false discovery rate (FDR < 0.05; **Extended Data Fig. 8h**).

##### Chronological Activation Pseudotime and state-zone differential abundance

To quantify continuous T-cell activation progression along the 30-dimensional basal state manifold (*u*_state_), we constructed Chronological Activation Pseudotime (CPT, *τ*_state_ ∈ [0, 1]; **Fig. 4e, f and Extended Data Fig. 9a, b**). Seven sequential biological reference landmarks (*L*_0_ to *L*_6_) were defined in 30D *u*_state_ space spanning the deep quiescent Rest NTC apex (*L*_0_), bulk Rest NTC centroid (*L*_1_), Stim 8hr early-arrest and intermediate bridge centroids (*L*_2_, *L*_3_), Stim 8hr acute activation peak NTC centroid (*L*_4_), early-maturing Stim 48hr NTC centroid (*L*_5_), and mature effector apex Stim 48hr NTC centroid (*L*_6_). Each cell was projected onto the piecewise-linear geodesic *L*_0_ → *L*_1_ → ⋯ → *L*_6_ via Gaussian distance-weighted softmin interpolation (*σ*_soft_ = 0.75) and quantile-normalized to [0, 1], with monotonic alignment across Rest, 8hr, and 48hr evaluated via Spearman rank correlation (*ρ*) and three-condition Kruskal-Wallis *H*-tests in both NTC cells and all 120,545 cells. To map transcriptional dynamics along *τ*_state_, denoised expression (X_pertvi_log1p) of 75 curated CD4+ T-cell activation, metabolic, and checkpoint markers^122–124^ was projected across 50 uniform *τ*_*state*_ bins (Gaussian-smoothed with *σ* = 1.0 bin) and organized into eight sequential transcriptional waves (W1–W8), spanning naive quiescence and lymphoid homing (W1), proximal TCR signalosome remodeling (W2), immediate-early ignition and acute cytokines (W3), negative-feedback attenuators (W4), co-stimulatory and STAT5 survival signaling (W5), nucleotide, glutathione, and amino-acid anabolism (W6), helper lineage transcription factors (W7), and late homologous-recombination repair and coinhibitory checkpoint receptors (W8; **Fig. 4e and Extended Data Fig. 9c–e**). Conservation of activation wave trajectories between NTC cells (*n* = 6,024) and all 120,545 cells was evaluated via Pearson correlation (*r*) across the 50 bins. The *τ*_state_ continuum was further segmented into seven contiguous developmental zones (Zones 1–7, spanning resting quiescence in Zones 1–2, early 8 h arrest and ignition transition in Zones 3–5, lagging 48 h maturation in Zone 6, and the mature 48 h effector apex in Zone 7). Within-condition differential abundance (log_2_DA) relative to condition-matched NTC cells was computed across all 50 bins and discrete developmental zones, evaluated via two-sided Fisher’s exact tests with Benjamini-Hochberg FDR control (**Fig. 4e, f)**. Overall phenotypic excursion for each regulator was summarized by its standardized state shift across timepoints, with differences across the four perturbation classes (early, general, late, and NCG) evaluated via four-group Kruskal-Wallis tests and temporal amplification or attenuation between 8h and 48h evaluated via two-sided paired Wilcoxon signed-rank tests.

##### Signed phenotypic shift standardization, bulk RNA-seq validation, and negative control calibration

For each perturbation *g* and stimulation condition *c* ∈ {Rest, Stim 8hr, Stim 48hr}, raw phenotypic shifts along the activation and mitotic manifolds were calculated as the difference in mean pseudotime between perturbed cells and condition-matched NTC cells 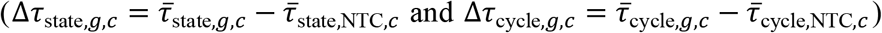. To account for finite-sample empirical dispersion and calibrate the negative control pool, we audited all 20 candidate NCGs across *u*_state_ and *u*_cycle_ via two-sided Welch’s *t*-tests, Mann-Whitney *U* tests, and leave-one-out empirical null standardization (**Extended Data Fig. 9h, i**). This audit identified 17 genuinely inert negative controls (Clean-NCG17) whose maximum joint two-dimensional state-cycle excursion norm across conditions 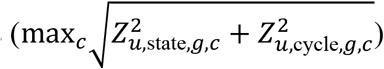 remained strictly bounded within a 1.90 SD empirical null ceiling, while isolating and rescuing three misannotated active regulators (*TBC1D1, IFT88*, and *PLK4*) that exceeded the null boundary. All raw state and cycle shifts were standardized against the empirical mean and standard deviation of Clean-NCG17 within each condition to yield signed *Z*-scores (*Z*_*u*,state,*g,c*_ and *Z*_*u*,cycle,*g,c*_). Projecting all perturbations into the two-dimensional signed state-shift plane (*Z*_*u*,state_ at 8hr versus *Z*_*u*,state_ at 48hr; **Fig. 4g and Extended Data Fig. 9f**) partitioned regulators into uni-directional positive promoters (*Z*_*u*,state_ < −1.0), negative-feedback suppressors (*Z*_*u*,state_ > +0.9), biphasic homeostatic rebounders that delayed acute 8h activation (*Z*_*u*,state,8h_ < −1.0) before inverting polarity to drive 48h hyper-activation (*Z*_*u*,state,48h_ > +0.9), and biphasic metabolic or DNA-repair crash switchers that shifted from mild 8h acceleration to 48h maturation collapse. To independently validate the biphasic polarity reversal of *ATP2A2* knockdown, we analyzed DESeq2^43^ Wald test statistics from the original study’s accompanying arrayed CRISPRi bulk RNA-seq dataset (GSE314342)^15^ at 8h and 48h post-stimulation across 649 TCR-activated genes and 645 TCR-repressed quiescence genes, evaluating directional reversal between 8h and 48h via two-sided Wilcoxon tests (**Extended Data Fig. 9g**).

##### Counterfactual causal scoring, inter-regulator correlation, and functional analysis

To isolate direct, condition-specific transcriptional responses in the perturbation-modulated state space (*z*_state_) free of cell-cycle and donor confounding, we computed gene-level counterfactual causal scores (CCS_*g,c*_ ∈ ℝ^10,004^) for each perturbation *g* in condition *c*, yielding the 10,004-gene counterfactual causal score vector CCS_*g,c*_. Overall transcriptional effect size was summarized by the Euclidean norm 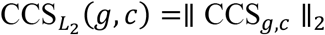 (*g, c*) =∥ CCS_*g,c*_ ∥_2_ and compared against log_10_(DEGs + 1)from the original study’s donor-pooled PyDESeq2^45^ differential expression table (FDR < 0.05) across all 81 perturbations at 8hr and 48hr (**Fig. 4h**). Stage-hierarchy resolution was assessed by comparing the temporal ratio 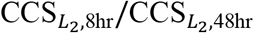 across early (*n* = 12), general (*n* = 29), and late (*n* = 20) regulator classes via two-sided Mann-Whitney *U*tests (**Extended Data Fig. 10a**) and by evaluating continuous receiver operating characteristic (ROC) curves comparing target regulators against Clean-NCG17 and early against late regulators (**Extended Data Fig. 10b**). Pairwise Pearson correlation coefficients 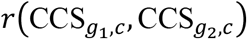 were computed across all 61 target regulators separately at Stim 8hr (upper triangle) and Stim 48hr (lower triangle; **Fig. 4i**). For functional landscape characterization (**Extended Data Fig. 10c–f**), genelevel CCS_*g,c*_ vectors were projected onto curated human transcription factor regulons from CollecTRI^119^ (retaining regulons with at least 5 overlapping target genes) using the univariate linear model (ULM) in decoupleR^118^ to obtain *t*-statistics across 36 curated T-cell and specificity-control transcription factors (**Extended Data Fig. 10c, d**). Pathway enrichment was computed on gene-level CCS_*g,c*_ rankings across 36 curated Gene Ontology Biological Process (GOBP) gene sets using pre-ranked Gene Set Enrichment Analysis (gseapy.prerank, gene set size 5 to 500, 300 permutations, random seed 42) to obtain normalized enrichment scores (NES; **Extended Data Fig. 10e, f**), alongside Clean-NCG17 mean profiles and three ectopic non-lymphoid GOBP negative-control pathways (Ossification GO:0030278, Chemical Synaptic Transmission GO:0007268, and Cardiac Muscle Contraction GO:0060048).

##### Cross-donor split-half reproducibility benchmarking and scVI mitotic confounding evaluation

To evaluate cross-donor replicability of 10,004-gene perturbation signatures across the four biological donors (**Extended Data Fig. 11b**), we partitioned the dataset into two disjoint donor halves comprising Donors D1+D2 and Donors D3+D4. Within each condition (Rest, 8hr, and 48hr) and across all 61 target regulators, we computed the median Pearson correlation (*r*) between the D1+D2 and D3+D4 perturbation effect vectors across five methods: the donor-pooled PyDESeq2 pseudobulk correlation reported in the original study^15^, raw single-cell observational differential expression (Raw Obs-DE, computed as mean log(1 + *x*)-normalized expression differences between perturbed and condition-matched NTC cells within each donor half), a donor-blocked pseudobulk generalized linear model (Donor PB-GLM, computed by centering each donor’s perturbation pseudobulk profile by that same donor’s matched NTC pseudobulk mean before averaging within each split half), 30D scVI (computed as the difference between perturbed and condition-matched NTC mean ridge-decoded expression with *L*_2_ penalty *λ* = 10.0 from the 30-dimensional scVI latent space within each donor half), and PertVI CCS, with pairwise method comparisons evaluated via two-sided paired Wilcoxon signed-rank tests. To evaluate whether mitotic sub-lobe fracturing in the unconstrained 30D scVI latent space confounds activation-state inference (**Fig. 4j, k and Extended Data Fig. 11c–f**), 30D scVI latent coordinates within each condition *c* were standardized relative to condition-matched NTC cells 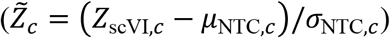 and projected onto the unit Chronological Activation Pseudotime (*τ*_state_) state direction 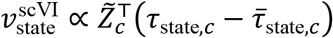 (**Extended Data Fig. 11c**) and the unit mitotic cell-cycle score (*s*_cyc_ = S_score + G2M_score) direction 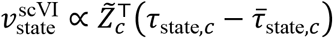 (**Extended Data Fig. 11d**), with local spatial gradient vectors overlaid on the 30D scVI UMAP embedding. For each of the 81 perturbations at Stim 8hr and Stim 48hr, sample-size-scaled scVI projected state and cycle shifts were standardized against the Clean-NCG17 baseline and compared head-to-head against PertVI *Z*_*u*,state_ and *Z*_*u*,cycle_ shifts via Pearson correlation (*r*) to quantify state-cycle collinearity (**Fig. 4j, k**). In the 48h two-dimensional (*Z*_state_, *Z*_cycle_) plane outside the Clean-NCG17 null envelope, regulators were classified across four quadrants into six mechanistic regimes (Reg. I: coupled TCR activation and proliferation block with *Z*_state_ < −1.0 and *Z*_cycle_ < −1.0; Reg. II: pure cell-cycle and DNA-repair checkpoints with ∣ *Z*_state_ ∣≤ 1.0 and *Z*_cycle_ < −1.8; Reg. III: S-phase metabolic replication trapping with *Z*_cycle_ > +1.5; Reg. IV-A: opposing-sign hyper-activating uncouplers with *Z*_state_ > +0.9 and *Z*_cycle_ < −1.0; Reg. IV-B: reverse uncouplers with *Z*_state_ < −1.0 and *Z*_cycle_ > +0.8; and Reg. V: pure activation-state modulators with ∣ *Z*_state_ ∣> 1.0 and ∣ *Z*_cycle_ ∣≤ 1.0; **Fig. 4k**). Empirical null stability was benchmarked by comparing the standard deviation of Clean-NCG17 perturbations along the state and cycle axes between 30D scVI and PertVI at 8h and 48h (**Extended Data Fig. 11e**), and directional classification accuracy was evaluated across the 12 mitotic-conflated regulators at 48h comprising five Reg. II pure cell-cycle and DNA-repair checkpoints (*RNF8, PHF23, FANCL, WAS*, and *FIGNL1*) and seven Reg. IV-A opposing-sign uncouplers (*LRP5, COG6, ORAI1, TLN1, CEP135, KMT2E*, and *ATP2A2*; **Extended Data Fig. 11f**).

## DATA AVAILABILITY

Single-cell RNA-seq data from the primary human CD8+ T-cell chemical perturbation screen^63^ (**Fig. 3 and Extended Data Figs. 3–7**) are available in NCBI Gene Expression Omnibus (GEO) under accession GSE306429. Single-cell CRISPRi Perturb-seq and arrayed bulk RNA-seq validation data from the primary human CD4+ T-cell multi-condition perturbation study^15^ (**Fig. 4 and Extended Data Figs. 8–11**) are available in NCBI GEO under accession GSE314342. External biological annotations were obtained from Tirosh et al.^18^ (cell-cycle signatures), CollecTRI^119^, and MSigDB^120^. The synthetic multi-covariate Perturb-seq dataset, processed AnnData (.h5ad) objects, trained model checkpoints, and summary tables supporting **Figs. 1– 4** and **Extended Data Figs. 1–11** are available at https://github.com/lenore-yue/PertVI. Source data are provided with this paper.

## CODE AVAILABILITY

The open-source PertVI Python package (PertVI, v3.1.0), built on PyTorch and scvi-tools^62^, is publicly available on GitHub at https://github.com/lenore-yue/PertVI. The repository provides all custom Python scripts and Jupyter notebooks for synthetic benchmark generation, hyperpa-rameter sweeps, single-cell preprocessing, and baseline model comparisons (ComBat^64^, LDA, scVI^30^, LinearSCVI^35^, MrVI^41^, ContrastiveVI^32^, DrVI^37^, and MoCAVI^42^. Scripts for downstream trajectory, regulon, pathway, and differential expression analyses, and for reproducing **Figs. 1–4** and **Extended Data Figs. 1–11**, are included in the same repository.

## ACKNOWLEDGEMENTS

We thank U. Kammula for his invaluable advice on the immunological aspects that guided model design, and for critically piloting and evaluating early versions of PertVI. This work was supported by NCI Pathway to Independence Award 1K99CA297120-01A1 to Y.W.

## AUTHOR CONTRIBUTION

Y.W. conceived the study, designed the model and methodology, implemented the software, conducted all formal analyses, acquired funding, and wrote the original manuscript. Y.Z. assisted with data curation and preprocessing, contributed to methodological discussions, and edited the manuscript. S.L. contributed to methodological discussions and manuscript review. D.Y. provided overall mentorship and supervisory support. All authors contributed to the critical review of the manuscript.

## DECLARATION OF AI USAGE

During the preparation of this work, the authors utilized Gemini Argon and Claude Opus 5.5 for code refactoring/debugging assistance, grammatical refinement, and high-level schematic design. Generative AI tools were **not** used to synthesize/modify images, generate synthetic data, or derive any formal scientific conclusions. All visualizations are plotted through deterministic code, and all plotting scripts and final figures were manually inspected, cross-checked against raw data tables, and verified by the authors. The authors take full responsibility for the accuracy and integrity of all code, figures, and manuscript content.

## Figure Legends

**Extended Data Fig. 1. Ground-truth generative design and multi-covariate structure of the synthetic Perturb-seq dataset**.

**a**, Mean ln(CP10K + 1) expression trajectories (*y*-axis) for G1 markers (genes 050–059, *n* = 10, cyan), S markers (genes 000–009, *n* = 10, orange), and G2M markers (genes 010–019, *n* = 10, red) in Batch0 (*n* = 4,989 cells, left) and Batch1 (*n* = 5,011 cells, right). Curves are plotted along continuous cell-cycle pseudo-time (*φ* ∈ [0, 3)) and along continuous differentiation pseudotime (*τ* ∈ [0, 2]). **b**, Single-cell ln(CP10K + 1) expression heatmaps across the 30 cell-cycle markersfor Batch0 and Batch1, with cells ordered along *φ* ∈ [0, 3). **c**, Mean ln(CP10K + 1) expression trajectories (*y*-axis) for State1 markers (genes 020–029, *n* = 10), State2 markers (genes 030–039, *n* = 10), and State3 markers (genes 040–049, *n* = 10) in Batch0 (left) and Batch1 (right), plotted along continuous differentiation pseudotime (*τ* ∈ [0, 2]) and along continuous cell-cycle pseudotime (*φ* ∈ [0, 3)). **d**, Single-cell ln(CP10K + 1) expression heatmaps across the 30 differentiation state markers for Batch0 and Batch1, with cells ordered along *τ* ∈ [0, 2]. **e**, Binary target-gene architecture heatmap across all *G* = 100 genes grouped by marker category across the five experimental conditions (black for targeted, white for untargeted). **f**, Pairwise Pearson correlation matrix among the ground-truth perturbation LFC vectors across all *G* = 100 genes for conditions A, B, C, and D. **g**, Total UMI library size distributions per cell (total UMIs per cell on the *x*-axis versus density on the *y*-axis) comparing Batch0, Batch1, and the combined dataset. **h**, Two-dimensional cycle-by-state interaction heatmaps of mean cell-cycle marker expression, plotting cell-cycle pseudotime *φ* ∈ [0, 3) (*x*-axis) against differentiation pseudotime *τ* ∈ [0, 2] (*y*-axis) for G1 markers, S markers, and G2M markers, illustrating vertical expression modulation along *τ* from state-to-cell-cycle cross-coupling. **i**, Two-dimensional state-by-cycle interaction heatmaps of mean state marker expression, plotting cell-cycle pseudotime *φ* ∈ [0, 3) (*x*-axis) against differentiation pseudotime *τ* ∈ [0, 2](*y*-axis) for State1, State2, and State3 markers, illustrating horizontal expression modulation along *φ* from cell-cycle-to-state cross-coupling.

**Extended Data Fig. 2. Training convergence, local neighborhood mixing, DrVI split-latent evaluation, and inter-perturbation correlation recovery on simulated data**.

**a**, Training (top row) and validation (bottom row) Negative Binomial and ZINB reconstruction loss trajectories across training epochs (*x*-axis) for the six probabilistic generative models. Final training and validation reconstruction losses are shown for LinearSCVI, DrVI, ContrastiveVI, MoCAVI, MrVI, and PertVI. **b**, Heatmap of log_10_(median LISI) across 17 latent representations. Scores are evaluated across Batch, CC Phase, Cell State, and Perturbation. **c**, Scatter plots of reconstructed Palantir differentiation pseudotime ([0, 1], *y*-axis) against ground-truth Cell State pseudotime across all *N* = 10,000 cells for DrVI nsplit = 5, ContrastiveVI background, PCA after ComBat, and uncorrected PCA. **d**, UMAP embeddings of individual split latent blocks from DrVI (nsplit = 3) and DrVI (nsplit = 5). **e**, Heatmaps of aF1 score and log_10_(median LISI) across the three split blocks of DrVI (nsplit = 3) and five split blocks of DrVI (nsplit = 5) for Batch, CC Phase, Cell State, and Perturbation. **f**, Satter-plot matrix for LinearSCVI comparing predicted LFC against ground-truth LFC (*x*-axis, columns A, B, C, D) across all *G* = 100 genes. **g**, Within-method pairwise Pearson correlation matrices across conditions computed among the inferred 100-gene perturbation effect vectors. Within-method correlations are shown for LinearSCVI, ContrastiveVI, MoCAVI, and PertVI.

**Extended Data Fig. 3. Disentanglement performance and cell cycle analysis (PertVI-CD8 model)**.

**a**, Model convergence curves across 0 to 600 training (top) and validation (bottom) epochs for Evidence Lower Bound (ELBO), Reconstruction Loss, and Total KL divergence (stabilizing near ~ 14.5without overfitting). **b**, 6-row × 12-column UMAP benchmark grid comparing Standard Approach (Orig., ComBat, LDA), MrVI (*u* and *z*), MoCAVI (base and salient), and PertVI (*u*_cycle_, *u*_state_, *z*_cycle_, *z*_state_, and *z*_final_) colored across six biological and technical overlays (Donor, Cell Cycle, Cell State, Perturbation, Cell-cycle Pseudo-time, and Cell-state Pseudotime), accompanied by 63-condition and 35-MoA legends. **c**, Quantitative 4 × 12 benchmark heatmaps of adjusted F1 score (aF1) and Local Inverse Simpson’s Index (LISI) across the 12 latent embeddings and four biological/technical factors (Donor, CC Phase, Cell State, and Perturbation). **d– h**, Resolution of the Late-S Crisis Zone (*t* ∈ [0.45, 0.65]) on *u*_cycle_: (d) canonical S-score and G2M-score trajectories along PertVI circular pseudotime showing premature score crossover at *t* ≈ 0.48; (e) min-max normalized expression waves ([0, 1]) of S-phase (*PCNA, TYMS*), S/G2 (*CCNA2*), and mitotic (*CCNB1, PLK1*) markers; (f) circular pseudotime KDE density curves comparing late-S crisis agents against mitotic blockers and DMSO; (g) discrete Canonical CC Phase stacked bars showing G2/M misclassification of camptothecin (60.7% G2/M) and serdemetan (54.7% G2/M) alongside 87.4% G2/M misclassification in Bin 5; and (h) PertVI CC Phase stacked bars rescuing camptothecin (49.9% Late-S) and serdemetan (55.3% Late-S) relative to true mitotic blockers BI 2536 (64.6% Early-G2/M) and tozasertib (72.6% Late-M).

**Extended Data Fig. 4. PertVI’s state space enables clean reconstruction of differentiation trajectory (PertVI-CD8 model)**.

**a**, 2 × 5 UMAP feature plot grid on *u*_state_ displaying denoised expression of 10 canonical differentiation markers across the five CD8+ T-cell states (*TCF7, LEF1, SELL, CCR7, IL7R, KLRG1, GZMB, IFNG, LAG3*, and *TOX*). **b, c**, Two-step identification of the four differentiation archetypes on *u*_state_: (b) 62-compound ×5state compositional abundance log_2_FC vs. DMSO heatmap comparing unconstrained Step 1: Unsupervised Clusters (55 shifters / 7 non-shifters) against dual-gate Step 2: Score-based filtering (*q*_KS_ < 0.05 or 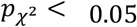 with max ∣ log_2_FC ∣≥ 0.20, pruning 34 non-shifting compounds with ∣ ΔPMF ∣≤ 0.0048 and resolving C1 Naive-Arrest *n* = 8, C2 Effector-promoting *n* = 10, and C3 Memory-shifting *n* = 10); and (c) 1 × 4 continuous DPT density shift curves (ΔPMF vs. DMSO, mean ±1 s.e.m. across DPT Deciles 1 to 10) for the four archetypes. **d**, Alluvial ribbon diagram mapping the eight high-level MoA categories (*N* = 62 compounds) to the four differentiation archetypes with active state-shifting compound callouts.

**Extended Data Fig. 5. PertVI *u***_***state***_ **space reveals the phenotypic landscape of anti-cancer drug’s regulation on CD8+ T cell effector differentiation**.

**a, b**, Multi-resolution MoA enrichment analysis: (a) stacked proportion bars and log_2_(OR)forest plot of binary state-shifting ability across the eight high-level MoAs and merged Categories 7+8); and (b) 8 × 3 active-shifter routing (*N* = 28) standardized residual matrix (*z*_adj_) and multi-resolution bar chart highlighting 100%C1 Naive-arrest routing of mTOR inhibitors and the two-way macro-class division of labor between kinase inhibitors in C1 and active non-kinase modulators in C2/C3. **c, d**, Single-cell target transcript auditing and missing-target gate across *N*_DMSO_ = 6,472 unperturbed CD8+ T cells and 25,078 genes: (c) dotplot of basal target gene expression across the five T-cell states (left) and horizontal bar plot of percentage of DMSO CD8+ T cells with UMI > 0 (right), contrasting ubiquitous expression of Category 1 intracellular kinases (median 66.59%) and *ZAP70* (74.89%) with near-absence of Category 2 solid-tumor RTK and B-cell/myeloid *SYK* (0.99%; Category 2 median 1.64%, *p* = 0.00019) targets; and (d) scatter plot of single-cell target detection frequency versus phenotypic shift magnitude (max ∣ log_2_FC ∣), demonstrating the < 5% and < 15% missing-target inert zones and off-target polypharmacological rescue of neratinib.

**Extended Data Fig. 6. PertVI *z***_***state***_ **space reveals the functional landscape of anti-cancer drug’s regulation on CD8+ T cell effector differentiation**.

**a**, Detailed 4-Quadrant Decoupling landscape. **b**, Dot matrix across 12 key transcription factors and four GOBP pathways comparing the five divergent same-MoA sibling groups. **c**, Pairwise Pearson correlation boxplots across the six functional hierarchy levels for 32 curated TFs (left) and 33 curated GOBP pathways spaces (right). **d**, Dual-panel 1,000-permutation PERMANOVA variance partitioning (*R*^2^%) trajectories across the global momentum and five developmental stages for TF Regulon Space and GOBP Pathway Space. **e, f**, Structured 3-layer dot matrices of 36 CollecTRI transcription factor regulons and 35 GOBP pathways across the eight sub-modules, Q3 compounds, and two Q4 summary columns (*n* = 20 buffered and *n* = 7 targetless RTK, totaling *n* = 27). **g**, 28 × 28 active-shifter pairwise correlation matrix ordered by sub-modules C1a to C3c (Upper Triangle: 32 curated TF regulons; Lower Triangle: 33 curated GOBP pathways). **h, i**, Stage-resolved (Mom, Tn, Prol, IL7R+, Teff, Term) archetype consensus vote dot matrices for the 36 TF regulons (h) and 35 GOBP pathways (i) across C1, C2, and C3. **j**, Stage-resolved decoupleR ULM *t*-statistic trajectory bifurcations across developmental stages for TP53 and E2F4.

**Extended Data Fig. 7. PertVI *v***_***ood***_ **space reveals off-manifold toxicity and stress responses across anticancer drugs**.

**a**, Bimodal two-component Gaussian Mixture Model (GMM) distribution of off-manifold displacement *d*_ood_ across all 62 compounds, defining the operational cutoff, DMSO 99.9th percentile bootstrap floor, and Tukey outlier fence. **b**, Bar chart of MoA enrichment among off-manifold perturbations (*d*_ood_ ≥ 0.30). **c, d**, Clean MCO gene-level stress signatures and universal interferon collapse: (c) MCO OOD log-FC heatmap of 18 signature genes (9 upregulated oxidative, xenobiotic, and ER stress genes and 9 downregulated interferon-stimulated and cytolytic genes) across the 14 off-manifold compounds and 3 on-manifold controls (ponatinib, sorafenib, BI 2536); and (d) Scatter plot of off-manifold displacement *d*_ood_ versus Hallmark Interferon Alpha Response NES across all 62 compounds (linear fit *r* = −0.36, *p* < 0.001). **e, f**, Stratification of the 14 off-manifold compounds into six functional toxicity classes: (e) four discriminating Hallmark pathway NESbox-plots; and (f) 48-Hallmark-pathway dot-matrix clustermap stratified into five functional pathway programs.

**Extended Data Fig. 8. Disentanglement performance and cell cycle analysis (PertVI-CD4 model)**.

**a**, Summary table of high-quality single-guide cell counts (*n* = 120,545 total) across the four healthy blood donors and three TCR stimulation conditions. **b**, Training (80% split, *n* = 96,436 cells, left) and Validation (20% split, *n* = 24,109 cells, right) optimization curves across 400epochs showing evidence lower bound (ELBO), Negative Binomial Reconstruction Loss, and Total KL divergence. **c**, Left: Complete UMAP grid across all 120,545 cells comparing PertVI’s six latent representations (*u*_cycle_ ∈ ℝ^3^, *u*_state_ ∈ ℝ^30^, *z*_cycle_ ∈ ℝ^3^, *z*_state_ ∈ ℝ^30^, *v*_ood_ ∈ ℝ^5^, and *z*_final_ ∈ ℝ^38^), colored by Donor, Cell Cycle, Cell State, Perturbation, and Control Type. Right: Catalog of all 82 perturbation labels in the pilot dataset grouped by functional pathway module and colored by Regulatory Stage. **d, e**, Two-dimensional principal component projections (PC1 (75.54% Var) versus PC2 (24.40% Var), capturing 99.94% of total variance) of the 3D basal cell-cycle subspace (*u*_cycle_, *n* = 120,545 cells), colored in **d** by discrete Canonical Phase overlaid with the 6-Waypoint Geodesic polygonal curve, and colored in **e** by Continuous Cell-Cycle Pseudotime (*τ*_cycle_ ∈ [0.0, 1.0]). **f**, Min-max scaled denoised expression (Scaled Wave Expression (X_pertvi_log1p), [0.0, 1.0]) trajectories of the five sequential mitotic transcriptional waves (W1 (C1 Early/G1/G0): Quiescence & Restraint; W2 (C2 Act G1): Restriction & Anabolic Ignition; W3 (C3 Early/Mid-S): Origin Licensing & DNA Synthesis; W4 (C4 Late-S/G2): Replication-Fork & HR Repair; and W5 (C5 Late-M): Mitotic Spindle & Cytokinesis) along Continuous Cell-Cycle Pseudotime (*τ*_cycle_ ∈ [0.0, 1.0]), comparing unperturbed NTC Controls (n=6,024) (Solid curves) and All Cells (n=120,545) (Dashed curves). Top color bar marks zones C1–C5. **g**, Stacked heatmaps of Z-Scored PertVI Expression (−2.0 to +2.0) for canonical cell-cycle markers ordered by peak pseudotime across zones C1–C5, comparing NTC Controls Only (n = 6,024) (top) and All Cells (n = 120,545) (bottom; marker-level median Pearson *r* = 0.9998). **h**, Heatmap of Within-Condition log2(DA vs. NTC) across cellcycle phase zones C1–C5 at Rest, Stim 8hr, and Stim 48hr for all 61 curated target regulators ordered by stage (Early, General, and Late). Exact log_2_DA values and two-sided Fisher’s exact test FDR are annotated (* FDR < 0.05, ** FDR < 0.01, *** FDR < 0.001).

**Extended Data Fig. 9. PertVI’s state space enables clean reconstruction of activation continuum (PertVI-CD4 model)**.

**a, b**, Condition Composition and Density Along CPT (*τ*_state_ ∈ [0.0, 1.0] across 50 uniform bins) on *u*_state_ for NT Cells Only (n = 6,024) and all Cells (n = 120,545). Left *y*-axis (Condition Fraction, stacked area plot) displays the proportion of Rest, Stim 8hr, and Stim 48hr cells per bin; right *y*-axis (solid black curve) displays log-transformed cell density log_10_(Cells/Bin + 1). Vertical dashed lines mark Rest Med (0.16), 8hr Peak (0.52), and 48hr Med (0.82). **c**, Side-by-side heatmaps of Canonical Marker Expression (PertVI corrected) (Row-Normalized Expression, [0.0, 1.0]) for 75 canonical CD4+ T-cell activation markers organized into eight sequential waves (W1: Quiescence, W2: TCR Signalosome, W3: Immediate-Early, W4: Neg. Feedback, W5: Costim & STAT5, W6: Metabolic Anabolism, W7: Lineage TFs, and W8: Checkpoints & Cycle) across 50 *τ*_state_ bins, comparing NT cells only (left) and all Cells (*r*ight; marker-level median Pearson *r* = 0.9912). **d, e**, Canonical Wave Trajectories along *τ*_state_ ∈ [0.0, 1.0] for NT cells only (*n* = 6,024, **d**) and all Cells (r=0.985 vs. NT) (*n* = 120,545), with wave peak coordinates (Pk) indicated in the legend. **f**, Bar chart of within-condition CPT log2 DA vs. NTC for the 10 biphasic regulators (7 homeostatic rebounders and 3 metabolic and DNA-repair crash switchers, separated by a horizontal dashed line). **g**, Violin and embedded box plots of DESeq2 Wald test statistics from the original study’s independent arrayed CRISPRi bulk RNA-seq dataset following *ATP2A2* knockdown at 8hr and 48hr across 649 TCR-activated genes (Act. 8hr median −0.56, *P* = 1.62 × 10^−30^; Act. 48hr median +0.31, *P* = 1.44 × 10^−7^) and 645 TCR-repressed quiescence genes (Quies. 8hr median +0.50, *P* = 7.62 × 10^−17^; Quies. 48hr median −0.42, *P* = 1.95 × 10^−14^; two-sided Wilcoxon tests). Center line, median; box limits, upper and lower quartiles; whisk-ers, 1.5 ×interquartile range; horizontal dashed line, Wald stat = 0. **h**, Scatterplot comparing donor-pooled Pooled PyDESeq2 log10(DEGs + 1) (*x*-axis) against PertVI Excursion |Z| (symlog *y*-axis), highlighting paralog-buffered or unexpressed perturbations shrunk by PertVI (Pooled FP Shrunk, orange downward triangles: *ATP2A3, UBIAD1, FANCL, WAS*) below the Clean-NCG17 threshold (horizontal dotted line at ∣ *Z* ∣ = 1.90) and bona fide active regulators rescued by PertVI (PertVI FN Rescue, red upward triangles: *PLK4, TBC1D1, IFT88*, and *ATP2A2*; vertical dashed line marks 75 DEGs). **i**, Ranked bar chart of PertVI Excursion |Z| Above Clean-NCG17 (SD) across all 20 candidate NCGs, separating the 17 validated inert controls (Clean-NCG17, grey bars bounded below the Clean-NCG17 Max (1.90) dashed line) from the three rescued active regulators (*PLK4, TBC1D1*, and *IFT88*).

**Extended Data Fig. 10. PertVI’s counterfactual analysis recapitulate regulator temporal hierarchy (PertVI-CD4 model)**.

**a**, Box-and-jitter plot of the cross-timepoint counterfactual causal score effect-size ratio (CCS Ratio (8hr / 48hr), 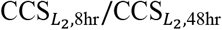) across Early (*n* = 12, median 1.80, interquartile range [IQR] 1.26–2.57), General (*n* = 29, median 1.68, IQR 1.18–2.04), and Late (*n* = 20, median 0.71, IQR 0.55–0.85) regulators. Center line, median; box limits, upper and lower quartiles; whiskers, 1.5 × IQR; horizontal dashed line marks a ratio of 1.0. **b**, Receiver operating characteristic (ROC) curves evaluating classification performance across three benchmarks: true target regulators (*n* = 58) versus Clean-NCG17 (*n* = 17) using 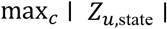 (Target vs NCG CCS (0.94)), Early (*n* = 12) versus Late (*n* = 20) regulators using 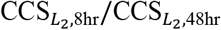 (Early vs Late Ratio (0.92)), and true target regulators (*n* = 58) versus Clean-NCG17 (*n* = 17) using max_*c*_ ∣ *Z*_*u*,state_ ∣ (Target vs NCG State Shift (0.87)). Diagonal grey dashed line indicates random chance (AUROC = 0.50). **c, d**, Dual-layer dot-and-tile heatmaps of the CollecTRI transcription factor regulon activity landscape (TF Regulon Landscape) at 8hr Stimulation (Acute TCR Ignition) (**c**) and 48hr Stimulation (Late Blastogenic & Effector) (**d**) across 36 curated CollecTRI TF regulons (rows, including four unexpressed ectopic lineage controls at the bottom: *SP7, HNF4A, MYOD1*, and *NEUROD1*, mRNA=0%) and 64 columns (61 target regulators grouped into Early (n=12), General Stage (n=29: 21 TCR + 8 Non-TCR), Late Promote (n=17), and Supp(3), plus three NCG summary columns: Clean NCG17 Mean, PLK4 [rescued NCG], and TBC1D1 [rescued NCG]). Top color bars denote regulator stage and pathway category. Background tile color encodes direct condition-specific ULM *t*-statistic; overlaid circle color encodes cross-timepoint interaction shift (TF Rest / Delta_CSS ULM t); circle size encodes −log_10_(ULM *p*). **e, f**, Dual-layer dot-and-tile heatmaps of the GOBP Pathway Landscape enrichment at 8hr Stimulation (**e**) and 48hr Stimulation (**f**) across 36 curated GOBP pathways (rows, including three bottom ectopic non-lymphoid negative-control pathways: Ossification (GO:0030279), Synaptic Transm. (GO:0007268), and Cardiac Muscle (GO:0060048)) and the same perturbation columns as in **c, d**. Background tile color encodes direct condition-specific preranked GSEA normalized enrichment score (Pathway CSS NES); overlaid circle color encodes cross-timepoint interaction shift (Pathway Rest / Delta_CSS NES); circle size encodes −log_10_(NOM *p*).

**Extended Data Fig. 11. Benchmark of PertVI against SCVI in disentangling mitotic variances (PertVI-CD4 model)**.

**a**, UMAP of the dimensionality-matched, unconstrained 30-dimensional scVI baseline embedding (30D SCVI Latent Embedding, *n* = 120,545 cells, n_latent = 30, batch_key=“donor”), colored by Donor, Cell Cycle Phase, Condition, Perturbation, and Control Type. **b**, Grouped bar chart comparing Median Split-Half r (61 Targets) at Rest, Stim 8hr, and Stim 48hr across five methods: original study donor-pooled Pub. PyDESeq2, Raw Obs-DE, Donor PB-GLM, 30D scVI, and PertVI CCS. **c, d**, UMAP embeddings of the 30D scVI latent space colored by within-condition NTC-standardized score along the Chronological Activation Pseudotime state axis (SCVI CPT State Projection, **c**) and the continuous mitotic cell-cycle score axis (SCVI Cycle Score Projection, S_score + G2M_score, **d**). Overlaid arrows inside the 8hr and 48hr condition islands indicate local spatial gradient vectors for the Cycle Gradient Vector and State Gradient Vector. **e**, Two-dimensional state-versus-cycle shift scatterplots (symlog axes, dotted zero lines) comparing empirical null calibration across annotated regulators (61), Clean-NCG (17), Clean-NCG Min-Max bounding box (dashed rectangle), and Rescued NCGs (red upward triangles) for 8hr 30D scVI, 8hr PertVI, 48hr 30D scVI, and 48hr PertVI. **f**, Grouped bar chart comparing Clean-NCG17-standardized 48 h state shifts (48hr State Shift (Z-scored), symlog *y*-axis) between scVI Projected State Shift (Z-scored) (blue bars) and PertVI u-State Shift (Z-scored) (red bars) across 12 mitotic-conflated regulators separated by a vertical dashed line: five Reg. II: Pure Cycle/DDR Checkpoints and seven Reg. IV-A: State+/Cycle-Uncouplers.

