## Extended Data Fig. 1 for "Disentangled generative modeling of single-cell perturbation responses across dynamic cellular states"

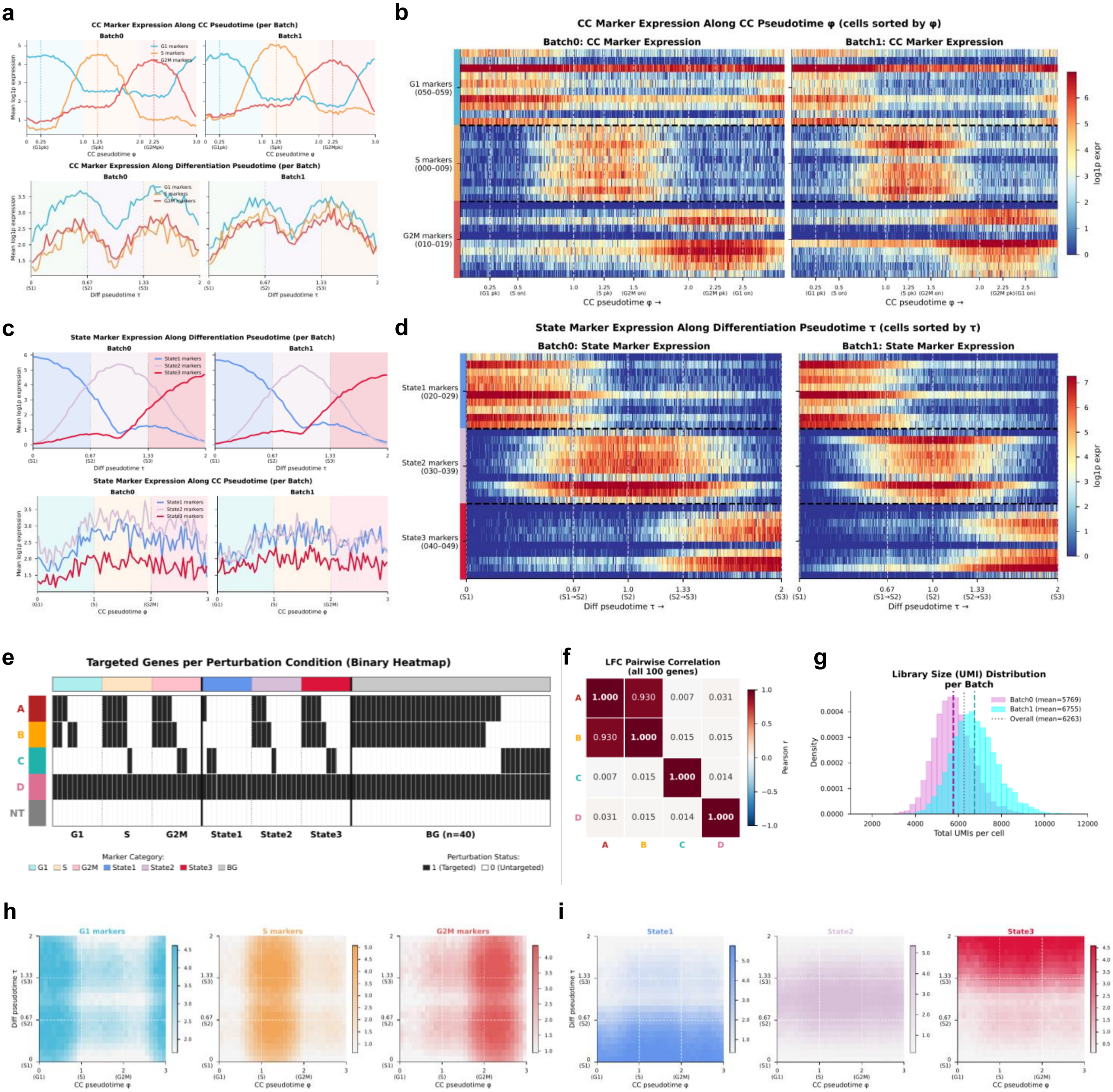
a
CC Marker Expression Along CC Pseudotime (per Batch)
Batch0
Batch1
G1 markers
S markers
G2M markers
Mean log1p expression
CC pseudotime φ
CC Marker Expression Along Differentiation Pseudotime (per Batch)
Diff pseudotime τ
b
CC Marker Expression Along CC Pseudotime φ (cells sorted by φ)
Batch0: CC Marker Expression
Batch1: CC Marker Expression
G1 markers (050–059)
S markers (000–009)
G2M markers (010–019)
CC pseudotime φ →
log1p expr
c
State Marker Expression Along Differentiation Pseudotime (per Batch)
State1 markers
State2 markers
State3 markers
State Marker Expression Along CC Pseudotime (per Batch)
d
State Marker Expression Along Differentiation Pseudotime τ (cells sorted by τ)
Batch0: State Marker Expression
Batch1: State Marker Expression
State1 markers (020–029)
State2 markers (030–039)
State3 markers (040–049)
Diff pseudotime τ →
e
Targeted Genes per Perturbation Condition (Binary Heatmap)
A
B
C
D
NT
G1
S
G2M
State1
State2
State3
BG (n=40)
Marker Category:
G1 S G2M State1 State2 State3 BG
Perturbation Status:
1 (Targeted) 0 (Untargeted)
f
LFC Pairwise Correlation (all 100 genes)
A 1.000 0.930 0.007 0.031
B 0.930 1.000 0.015 0.015
C 0.007 0.015 1.000 0.014
D 0.031 0.015 0.014 1.000
Pearson r
g
Library Size (UMI) Distribution per Batch
Batch0 (mean=5769)
Batch1 (mean=6755)
Overall (mean=6263)
Density
Total UMIs per cell
h
G1 markers
S markers
G2M markers
Diff pseudotime τ
CC pseudotime φ
i
State1
State2
State3
