## Extended Data Fig. 2 for "Disentangled generative modeling of single-cell perturbation responses across dynamic cellular states"

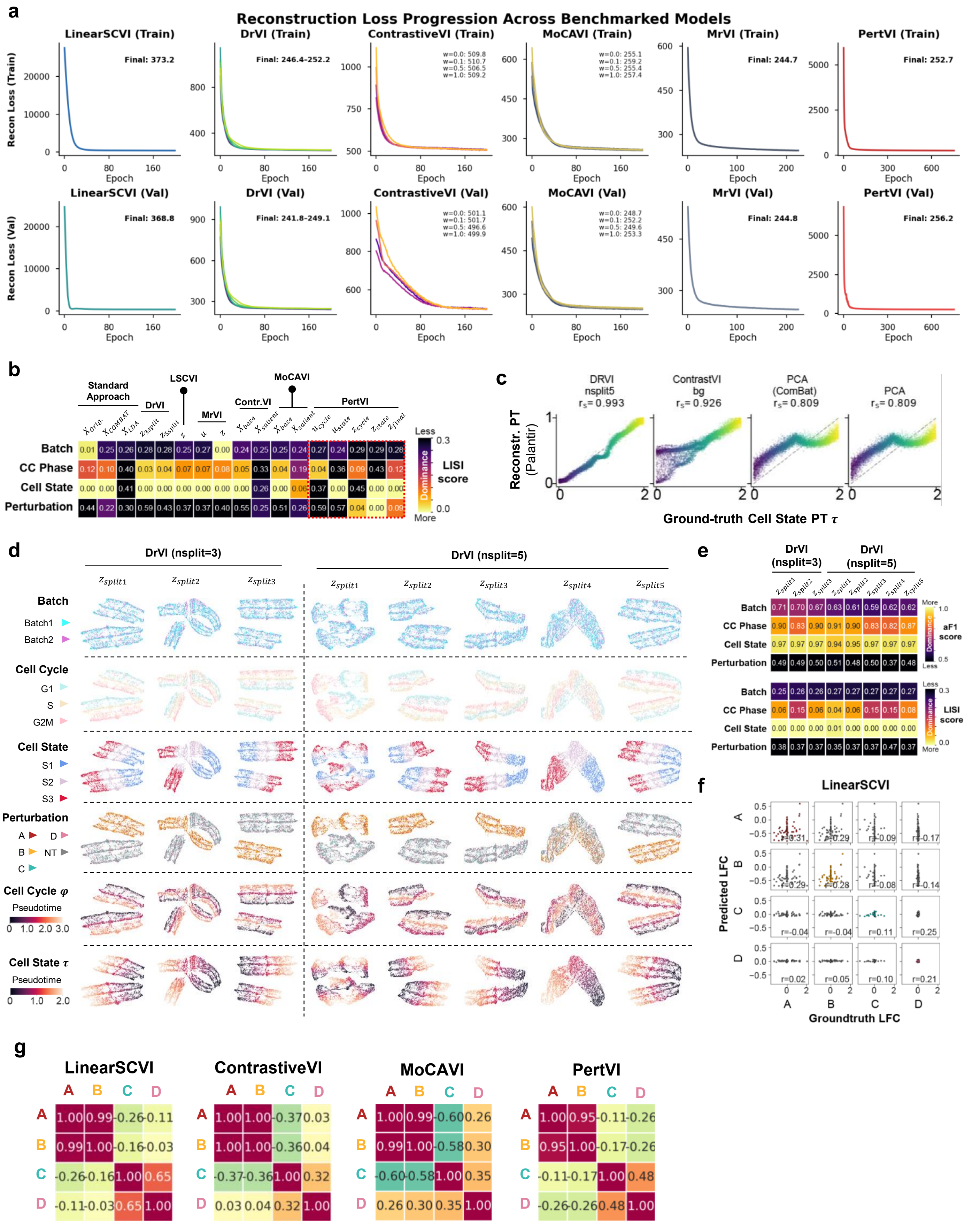
a
Reconstruction Loss Progression Across Benchmarked Models
LinearSCVI (Train)
Final: 373.2
DrVI (Train)
Final: 246.4-252.2
ContrastiveVI (Train)
w=0.0: 509.8
w=0.1: 510.7
w=0.5: 506.5
w=1.0: 509.2
MoCAVI (Train)
w=0.0: 255.1
w=0.1: 259.2
w=0.5: 255.4
w=1.0: 257.4
MrVI (Train)
Final: 244.7
PertVI (Train)
Final: 252.7
Recon Loss (Train)
Epoch
LinearSCVI (Val)
Final: 368.8
DrVI (Val)
Final: 241.8-249.1
ContrastiveVI (Val)
w=0.0: 501.1
w=0.1: 501.7
w=0.5: 496.6
w=1.0: 499.9
MoCAVI (Val)
w=0.0: 248.7
w=0.1: 252.2
w=0.5: 249.6
w=1.0: 253.3
MrVI (Val)
Final: 244.8
PertVI (Val)
Final: 256.2
Recon Loss (Val)
Epoch
b
Standard Approach
DrVI
LSCVI
MrVI
Contr.VI
MoCAVI
PertVI
Batch 0.01 0.25 0.26 0.28 0.28 0.25 0.27 0.00 0.24 0.25 0.25 0.24 0.27 0.24 0.29 0.29 0.28
CC Phase 0.12 0.10 0.40 0.03 0.04 0.07 0.07 0.08 0.05 0.33 0.04 0.19 0.04 0.36 0.09 0.43 0.12
Cell State 0.00 0.00 0.41 0.00 0.00 0.00 0.00 0.00 0.00 0.26 0.00 0.06 0.37 0.00 0.45 0.00 0.00
Perturbation 0.44 0.22 0.30 0.59 0.43 0.37 0.37 0.40 0.55 0.25 0.51 0.26 0.59 0.57 0.04 0.00 0.09
Less
More
Dominance
LISI score
c
DRVI nsplit5 $r_s$= 0.993
ContrastVI bg $r_s$= 0.926
PCA (ComBat) $r_s$= 0.809
PCA $r_s$= 0.809
Reconstr. PT (Palantir)
Ground-truth Cell State PT $\tau$
d
DrVI (nsplit=3)
DrVI (nsplit=5)
$z_{split1}$ $z_{split2}$ $z_{split3}$ $z_{split4}$ $z_{split5}$
Batch
Batch1
Batch2
Cell Cycle
G1
S
G2M
Cell State
S1
S2
S3
Perturbation
A D
B NT
C
Cell Cycle $\varphi$
Pseudotime
0 1.0 2.0 3.0
Cell State $\tau$
Pseudotime
0 1.0 2.0
e
DrVI (nsplit=3)
DrVI (nsplit=5)
Batch 0.71 0.70 0.67 0.63 0.61 0.59 0.62 0.62
CC Phase 0.90 0.83 0.90 0.91 0.90 0.83 0.82 0.87
Cell State 0.97 0.97 0.97 0.94 0.95 0.97 0.97 0.97
Perturbation 0.49 0.49 0.50 0.51 0.48 0.50 0.37 0.48
aF1 score
Batch 0.25 0.26 0.26 0.27 0.27 0.27 0.27 0.27
CC Phase 0.06 0.15 0.06 0.04 0.06 0.15 0.15 0.08
Cell State 0.00 0.00 0.00 0.01 0.00 0.00 0.00 0.00
Perturbation 0.38 0.37 0.37 0.35 0.37 0.37 0.47 0.37
LISI score
f
LinearSCVI
r=0.31 r=0.29 r=-0.09 r=-0.17
r=0.29 r=0.28 r=-0.08 r=-0.14
r=-0.04 r=-0.04 r=0.11 r=0.25
r=0.02 r=0.05 r=0.10 r=0.21
Predicted LFC
Groundtruth LFC
g
LinearSCVI
A 1.00 0.99 -0.26 -0.11
B 0.99 1.00 -0.16 -0.03
C -0.26 -0.16 1.00 0.65
D -0.11 -0.03 0.65 1.00
ContrastiveVI
A 1.00 1.00 -0.37 0.03
B 1.00 1.00 -0.36 0.04
C -0.37 -0.36 1.00 0.32
D 0.03 0.04 0.32 1.00
MoCAVI
A 1.00 0.99 -0.60 0.26
B 0.99 1.00 -0.58 0.30
C -0.60 -0.58 1.00 0.35
D 0.26 0.30 0.35 1.00
PertVI
A 1.00 0.95 -0.11 -0.26
B 0.95 1.00 -0.17 -0.26
C -0.11 -0.17 1.00 0.48
D -0.26 -0.26 0.48 1.00
