## Extended Data Fig. 3 for "Disentangled generative modeling of single-cell perturbation responses across dynamic cellular states"

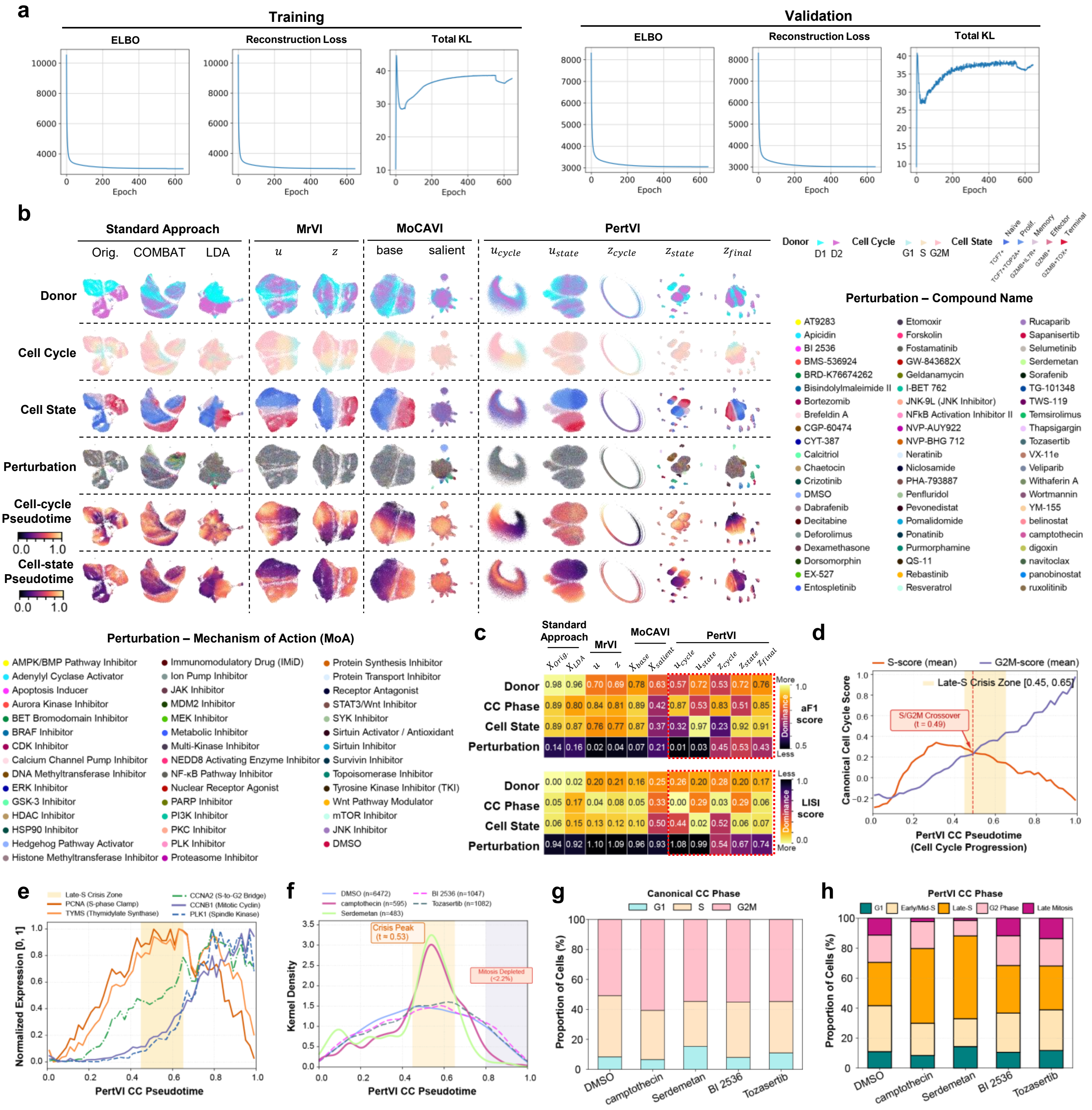
a
Training
Validation
ELBO
Reconstruction Loss
Total KL
Epoch
b
Standard Approach
Orig.
COMBAT
LDA
MrVI
MoCAVI
base
salient
PertVI
Donor
Cell Cycle
Cell State
Perturbation
Cell-cycle Pseudotime
Cell-state Pseudotime
D1 D2
G1 S G2M
Naive
Prolif.
Memory
Effector
Terminal
Perturbation – Compound Name
AT9283
Apicidin
BI 2536
BMS-536924
BRD-K76674262
Bisindolylmaleimide II
Bortezomib
Brefeldin A
CGP-60474
CYT-387
Calcitriol
Chaetocin
Crizotinib
DMSO
Dabrafenib
Decitabine
Deforolimus
Dexamethasone
Dorsomorphin
EX-527
Entospletinib
Etomoxir
Forskolin
Fostamatinib
GW-843682X
Geldanamycin
I-BET 762
JNK-9L (JNK Inhibitor)
NFkB Activation Inhibitor II
NVP-AUY922
NVP-BHG 712
Neratinib
Niclosamide
PHA-793887
Penfluridol
Pevonedistat
Pomalidomide
Ponatinib
Purmorphamine
QS-11
Rebastinib
Resveratrol
Rucaparib
Sapanisertib
Selumetinib
Serdemetan
Sorafenib
TG-101348
TWS-119
Temsirolimus
Thapsigargin
Tozasertib
VX-11e
Veliparib
Withaferin A
Wortmannin
YM-155
belinostat
camptothecin
digoxin
navitoclax
panobinostat
ruxolitinib
Perturbation – Mechanism of Action (MoA)
AMPK/BMP Pathway Inhibitor
Adenylyl Cyclase Activator
Apoptosis Inducer
Aurora Kinase Inhibitor
BET Bromodomain Inhibitor
BRAF Inhibitor
CDK Inhibitor
Calcium Channel Pump Inhibitor
DNA Methyltransferase Inhibitor
ERK Inhibitor
GSK-3 Inhibitor
HDAC Inhibitor
HSP90 Inhibitor
Hedgehog Pathway Activator
Histone Methyltransferase Inhibitor
Immunomodulatory Drug (IMiD)
Ion Pump Inhibitor
JAK Inhibitor
MDM2 Inhibitor
MEK Inhibitor
Metabolic Inhibitor
Multi-Kinase Inhibitor
NEDD8 Activating Enzyme Inhibitor
NF-κB Pathway Inhibitor
Nuclear Receptor Agonist
PARP Inhibitor
PI3K Inhibitor
PKC Inhibitor
PLK Inhibitor
Proteasome Inhibitor
Protein Synthesis Inhibitor
Protein Transport Inhibitor
Receptor Antagonist
STAT3/Wnt Inhibitor
SYK Inhibitor
Sirtuin Activator / Antioxidant
Sirtuin Inhibitor
Survivin Inhibitor
Topoisomerase Inhibitor
Tyrosine Kinase Inhibitor (TKI)
Wnt Pathway Modulator
mTOR Inhibitor
JNK Inhibitor
DMSO
c
Standard Approach
MrVI
MoCAVI
PertVI
Donor 0.98 0.96 0.70 0.69 0.78 0.63 0.57 0.72 0.53 0.72 0.76
CC Phase 0.89 0.80 0.84 0.81 0.89 0.42 0.87 0.53 0.83 0.51 0.85
Cell State 0.89 0.87 0.76 0.77 0.87 0.37 0.32 0.97 0.23 0.92 0.91
Perturbation 0.14 0.16 0.02 0.04 0.07 0.21 0.01 0.03 0.45 0.53 0.43
aF1 score
Donor 0.00 0.02 0.20 0.21 0.16 0.25 0.26 0.20 0.28 0.20 0.17
CC Phase 0.05 0.17 0.04 0.08 0.05 0.33 0.00 0.29 0.03 0.29 0.06
Cell State 0.06 0.15 0.13 0.12 0.10 0.50 0.44 0.02 0.52 0.06 0.07
Perturbation 0.94 0.92 1.10 1.09 0.96 0.93 1.08 0.99 0.54 0.67 0.74
LISI score
More
Less
Dominance
d
S-score (mean)
G2M-score (mean)
Late-S Crisis Zone [0.45, 0.65]
S/G2M Crossover (t = 0.49)
Canonical Cell Cycle Score
PertVI CC Pseudotime (Cell Cycle Progression)
e
Late-S Crisis Zone
PCNA (S-phase Clamp)
TYMS (Thymidylate Synthase)
CCNA2 (S-to-G2 Bridge)
CCNB1 (Mitotic Cyclin)
PLK1 (Spindle Kinase)
Normalized Expression [0, 1]
PertVI CC Pseudotime
f
DMSO (n=6472)
camptothecin (n=595)
Serdemetan (n=483)
BI 2536 (n=1047)
Tozasertib (n=1082)
Crisis Peak (t = 0.53)
Mitosis Depleted (<2.2%)
Kernel Density
PertVI CC Pseudotime
g
Canonical CC Phase
G1
S
G2M
Proportion of Cells (%)
DMSO
camptothecin
Serdemetan
BI 2536
Tozasertib
h
PertVI CC Phase
G1
Early/Mid-S
Late-S
G2 Phase
Late Mitosis
Proportion of Cells (%)
