## Extended Data Fig. 4 for "Disentangled generative modeling of single-cell perturbation responses across dynamic cellular states"

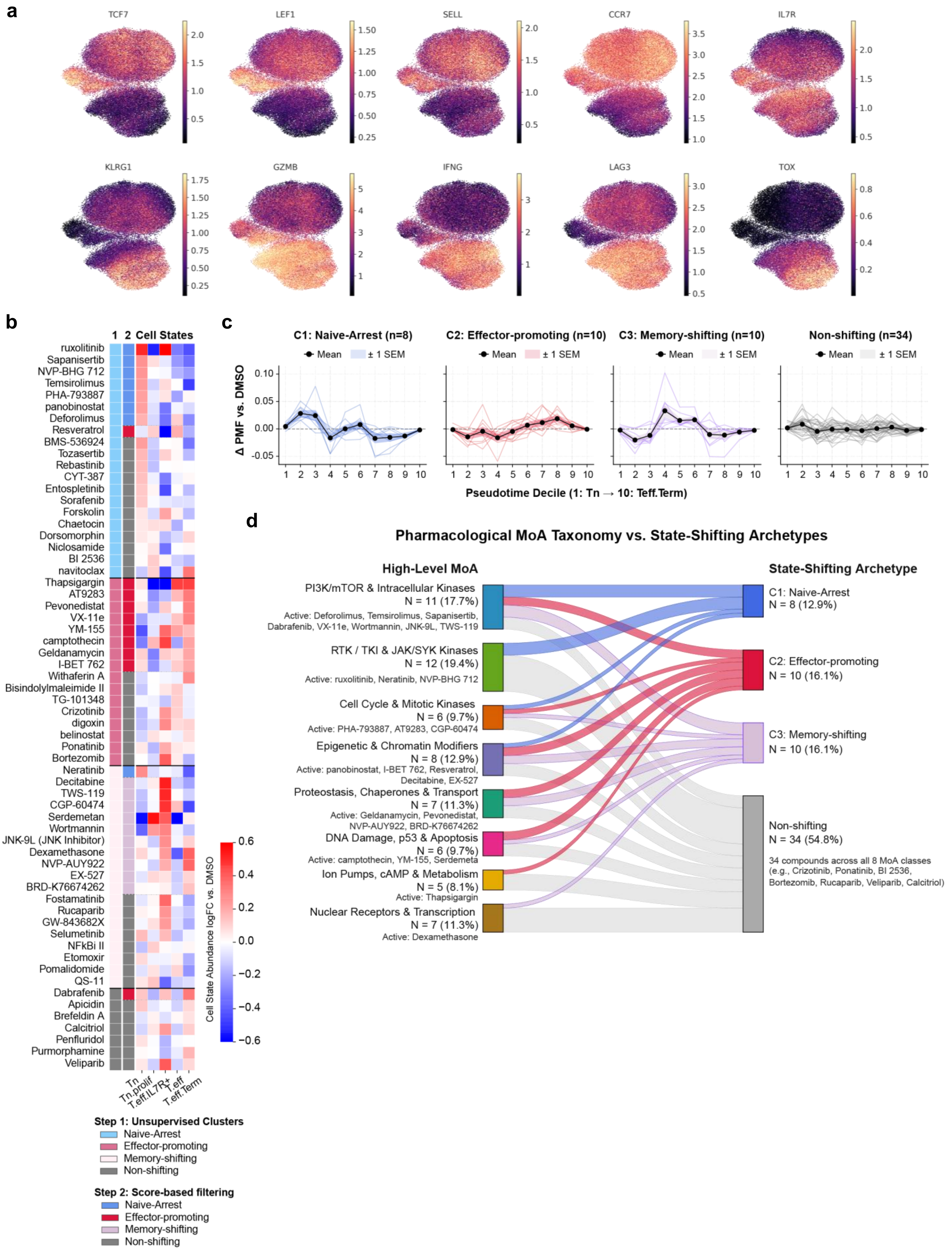
a
TCF7
LEF1
SELL
CCR7
IL7R
KLRG1
GZMB
IFNG
LAG3
TOX
b
1 2 Cell States
ruxolitinib
Sapanisertib
NVP-BHG 712
Temsirolimus
PHA-793887
panobinostat
Deforolimus
Resveratrol
BMS-536924
Tozasertib
Rebastinib
CYT-387
Entospletinib
Sorafenib
Forskolin
Chaetocin
Dorsomorphin
Niclosamide
BI 2536
navitoclax
Thapsigargin
AT9283
Pevonedistat
VX-11e
YM-155
camptothecin
Geldanamycin
I-BET 762
Withaferin A
Bisindolylmaleimide II
TG-101348
Crizotinib
digoxin
belinostat
Ponatinib
Bortezomib
Neratinib
Decitabine
TWS-119
CGP-60474
Serdemetan
Wortmannin
JNK-9L (JNK Inhibitor)
Dexamethasone
NVP-AUY922
EX-527
BRD-K76674262
Fostamatinib
Rucaparib
GW-843682X
Selumetinib
NFkBi II
Etomoxir
Pomalidomide
QS-11
Dabrafenib
Apicidin
Brefeldin A
Calcitriol
Penfluridol
Purmorphamine
Veliparib
Tn
Tn.prolif
T.eff.IL7R+
T.eff
T.eff.Term
Cell State Abundance logFC vs. DMSO
Step 1: Unsupervised Clusters
Naive-Arrest
Effector-promoting
Memory-shifting
Non-shifting
Step 2: Score-based filtering
Naive-Arrest
Effector-promoting
Memory-shifting
Non-shifting
c
C1: Naive-Arrest (n=8)
C2: Effector-promoting (n=10)
C3: Memory-shifting (n=10)
Non-shifting (n=34)
Mean
± 1 SEM
Δ PMF vs. DMSO
Pseudotime Decile (1: Tn → 10: Teff.Term)
d
Pharmacological MoA Taxonomy vs. State-Shifting Archetypes
High-Level MoA
State-Shifting Archetype
PI3K/mTOR & Intracellular Kinases
N = 11 (17.7%)
Active: Deforolimus, Temsirolimus, Sapanisertib, Dabrafenib, VX-11e, Wortmannin, JNK-9L, TWS-119
RTK / TKI & JAK/SYK Kinases
N = 12 (19.4%)
Active: ruxolitinib, Neratinib, NVP-BHG 712
Cell Cycle & Mitotic Kinases
N = 6 (9.7%)
Active: PHA-793887, AT9283, CGP-60474
Epigenetic & Chromatin Modifiers
N = 8 (12.9%)
Active: panobinostat, I-BET 762, Resveratrol, Decitabine, EX-527
Proteostasis, Chaperones & Transport
N = 7 (11.3%)
Active: Geldanamycin, Pevonedistat, NVP-AUY922, BRD-K76674262
DNA Damage, p53 & Apoptosis
N = 6 (9.7%)
Active: camptothecin, YM-155, Serdemeta
Ion Pumps, cAMP & Metabolism
N = 5 (8.1%)
Active: Thapsigargin
Nuclear Receptors & Transcription
N = 7 (11.3%)
Active: Dexamethasone
C1: Naive-Arrest
N = 8 (12.9%)
C2: Effector-promoting
N = 10 (16.1%)
C3: Memory-shifting
N = 10 (16.1%)
Non-shifting
N = 34 (54.8%)
34 compounds across all 8 MoA classes
(e.g., Crizotinib, Ponatinib, BI 2536, Bortezomib, Rucaparib, Veliparib, Calcitriol)
