## Extended Data Fig. 5 for "Disentangled generative modeling of single-cell perturbation responses across dynamic cellular states"

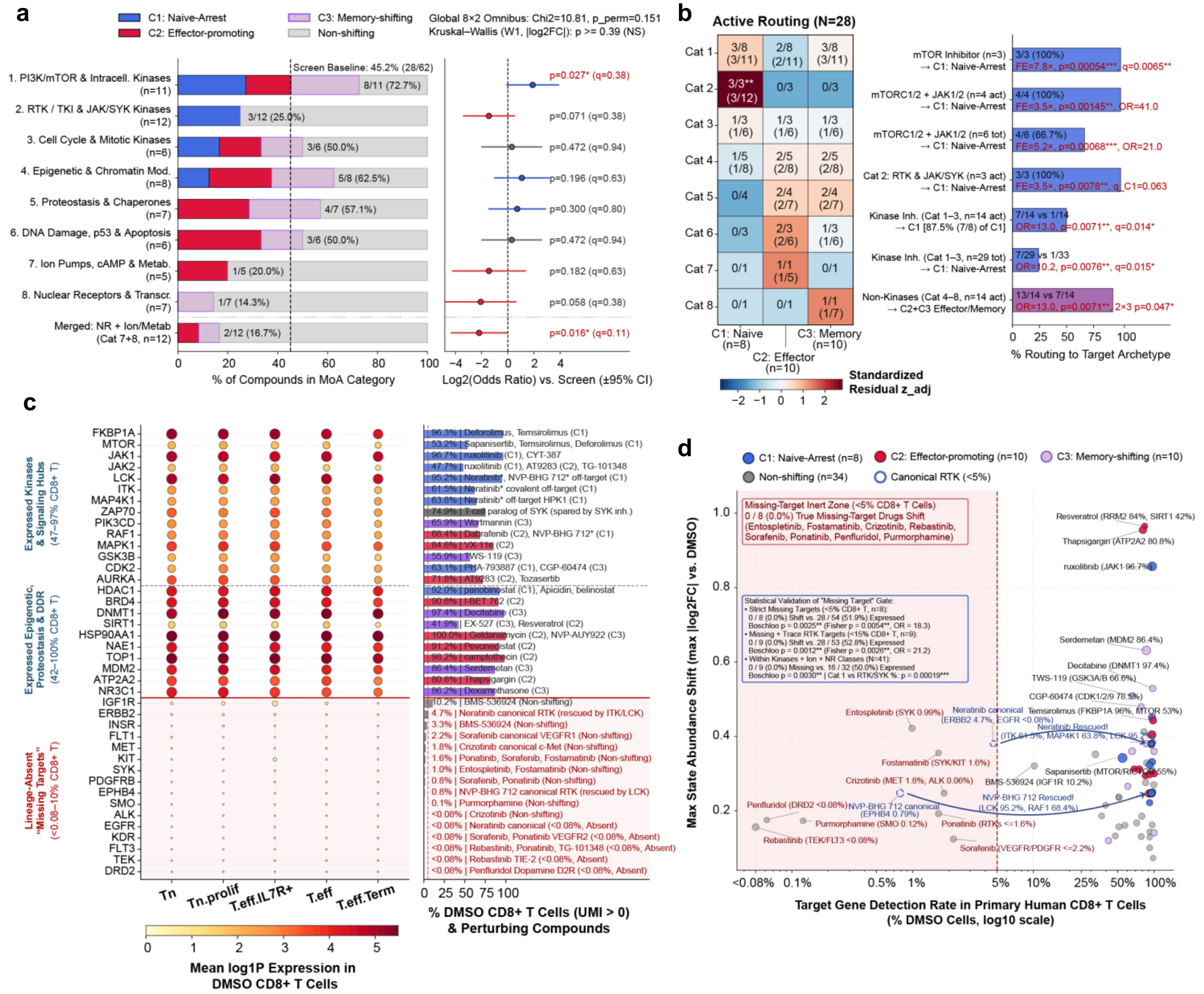

a
C1: Naive-Arrest
C2: Effector-promoting
C3: Memory-shifting
Non-shifting
Global 8×2 Omnibus: Chi2=10.81, p_perm=0.151
Kruskal–Wallis (W1, |log2FC|): p >= 0.39 (NS)
Screen Baseline: 45.2% (28/62)
1. PI3K/mTOR & Intracell. Kinases (n=11)
8/11 (72.7%)
p=0.027* (q=0.38)
2. RTK / TKI & JAK/SYK Kinases (n=12)
3/12 (25.0%)
p=0.071 (q=0.38)
3. Cell Cycle & Mitotic Kinases (n=6)
3/6 (50.0%)
p=0.472 (q=0.94)
4. Epigenetic & Chromatin Mod. (n=8)
5/8 (62.5%)
p=0.196 (q=0.63)
5. Proteostasis & Chaperones (n=7)
4/7 (57.1%)
p=0.300 (q=0.80)
6. DNA Damage, p53 & Apoptosis (n=6)
3/6 (50.0%)
p=0.472 (q=0.94)
7. Ion Pumps, cAMP & Metab. (n=5)
1/5 (20.0%)
p=0.182 (q=0.63)
8. Nuclear Receptors & Transcr. (n=7)
1/7 (14.3%)
p=0.058 (q=0.38)
Merged: NR + Ion/Metab (Cat 7+8, n=12)
2/12 (16.7%)
p=0.016* (q=0.11)
% of Compounds in MoA Category
Log2(Odds Ratio) vs. Screen (±95% CI)
b
Active Routing (N=28)
Cat 1: 3/8 (3/11) | 2/8 (2/11) | 3/8 (3/11)
Cat 2: 3/3** (3/12) | 0/3 | 0/3
Cat 3: 1/3 (1/6) | 1/3 (1/6) | 1/3 (1/6)
Cat 4: 1/5 (1/8) | 2/5 (2/8) | 2/5 (2/8)
Cat 5: 0/4 | 2/4 (2/7) | 2/4 (2/7)
Cat 6: 0/3 | 2/3 (2/6) | 1/3 (1/6)
Cat 7: 0/1 | 1/1 (1/5) | 0/1
Cat 8: 0/1 | 0/1 | 1/1 (1/7)
C1: Naive (n=8)
C2: Effector (n=10)
C3: Memory (n=10)
Standardized Residual z_adj
mTOR Inhibitor (n=3) → C1: Naive-Arrest
3/3 (100%) FE=7.8×, p=0.00054***, q=0.0065**
mTORC1/2 + JAK1/2 (n=4 act) → C1: Naive-Arrest
4/4 (100%) FE=3.5×, p=0.00145**, OR=41.0
mTORC1/2 + JAK1/2 (n=6 tot) → C1: Naive-Arrest
4/6 (66.7%) FE=5.2×, p=0.00068***, OR=21.0
Cat 2: RTK & JAK/SYK (n=3 act) → C1: Naive-Arrest
3/3 (100%) FE=3.5×, p=0.0078**, q_C1=0.063
Kinase Inh. (Cat 1–3, n=14 act) → C1 [87.5% (7/8) of C1]
7/14 vs 1/14 OR=13.0, p=0.0071**, q=0.014*
Kinase Inh. (Cat 1–3, n=29 tot) → C1: Naive-Arrest
7/29 vs 1/33 OR=10.2, p=0.0076**, q=0.015*
Non-Kinases (Cat 4–8, n=14 act) → C2+C3 Effector/Memory
13/14 vs 7/14 OR=13.0, p=0.0071**, 2×3 p=0.047*
% Routing to Target Archetype
c
Expressed Kinases & Signaling Hubs (47–97% CD8+ T)
FKBP1A, MTOR, JAK1, JAK2, LCK, ITK, MAP4K1, ZAP70, PIK3CD, RAF1, MAPK1, GSK3B, CDK2, AURKA
96.3% | Deforolimus, Temsirolimus (C1)
53.2% | Sapanisertib, Temsirolimus, Deforolimus (C1)
96.7% | ruxolitinib (C1), CYT-387
47.7% | ruxolitinib (C1), AT9283 (C2), TG-101348
95.2% | Neratinib*, NVP-BHG 712* off-target (C1)
61.5% | Neratinib* covalent off-target (C1)
63.8% | Neratinib* off-target HPK1 (C1)
74.9% | T-cell paralog of SYK (spared by SYK inh.)
65.9% | Wortmannin (C3)
68.4% | Dabrafenib (C2), NVP-BHG 712* (C1)
84.6% | VX-11e (C2)
55.9% | TWS-119 (C3)
63.1% | PHA-793887 (C1), CGP-60474 (C3)
71.8% | AT9283 (C2), Tozasertib
Expressed Epigenetic, Proteostasis & DDR (42–100% CD8+ T)
HDAC1, BRD4, DNMT1, SIRT1, HSP90AA1, NAE1, TOP1, MDM2, ATP2A2, NR3C1
92.0% | panobinostat (C1), Apicidin, belinostat
90.6% | I-BET 762 (C2)
97.4% | Decitabine (C3)
41.9% | EX-527 (C3), Resveratrol (C2)
100.0% | Geldanamycin (C2), NVP-AUY922 (C3)
91.2% | Pevonedistat (C2)
96.2% | camptothecin (C2)
86.4% | Serdemetan (C3)
80.8% | Thapsigargin (C2)
86.2% | Dexamethasone (C3)
Lineage-Absent "Missing Targets" (<0.08–10% CD8+ T)
IGF1R, ERBB2, INSR, FLT1, MET, KIT, SYK, PDGFRB, EPHB4, SMO, ALK, EGFR, KDR, FLT3, TEK, DRD2
10.2% | BMS-536924 (Non-shifting)
4.7% | Neratinib canonical RTK (rescued by ITK/LCK)
3.3% | BMS-536924 (Non-shifting)
2.2% | Sorafenib canonical VEGFR1 (Non-shifting)
1.8% | Crizotinib canonical c-Met (Non-shifting)
1.6% | Ponatinib, Sorafenib, Fostamatinib (Non-shifting)
1.0% | Entospletinib, Fostamatinib (Non-shifting)
0.8% | Sorafenib, Ponatinib (Non-shifting)
0.8% | NVP-BHG 712 canonical RTK (rescued by LCK)
0.1% | Purmorphamine (Non-shifting)
<0.08% | Crizotinib (Non-shifting)
<0.08% | Neratinib canonical (<0.08%, Absent)
<0.08% | Sorafenib, Ponatinib VEGFR2 (<0.08%, Absent)
<0.08% | Rebastinib, Ponatinib, TG-101348 (<0.08%, Absent)
<0.08% | Rebastinib TIE-2 (<0.08%, Absent)
<0.08% | Penfluridol Dopamine D2R (<0.08%, Absent)
Tn, Tn.prolif, T.eff.IL7R+, T.eff, T.eff.Term
Mean log1P Expression in DMSO CD8+ T Cells
% DMSO CD8+ T Cells (UMI > 0) & Perturbing Compounds
d
C1: Naive-Arrest (n=8)
C2: Effector-promoting (n=10)
C3: Memory-shifting (n=10)
Non-shifting (n=34)
Canonical RTK (<5%)
Missing-Target Inert Zone (<5% CD8+ T Cells)
0 / 8 (0.0%) True Missing-Target Drugs Shift
(Entospletinib, Fostamatinib, Crizotinib, Rebastinib, Sorafenib, Ponatinib, Penfluridol, Purmorphamine)
Statistical Validation of "Missing Target" Gate:
• Strict Missing Targets (<5% CD8+ T, n=8):
0 / 8 (0.0%) Shift vs. 28 / 54 (51.9%) Expressed
Boschloo p = 0.0025** (Fisher p = 0.0054**, OR = 18.3)
• Missing + Trace RTK Targets (<15% CD8+ T, n=9):
0 / 9 (0.0%) Shift vs. 28 / 53 (52.8%) Expressed
Boschloo p = 0.0012** (Fisher p = 0.0026**, OR = 21.2)
• Within Kinases + Ion + NR Classes (N=41):
0 / 9 (0.0%) Missing vs. 16 / 32 (50.0%) Expressed
Boschloo p = 0.0030** | Cat 1 vs RTK/SYK %: p = 0.00019***
Resveratrol (RRM2 84%, SIRT1 42%)
Thapsigargin (ATP2A2 80.8%)
ruxolitinib (JAK1 96.7%)
Serdemetan (MDM2 86.4%)
Decitabine (DNMT1 97.4%)
TWS-119 (GSK3A/B 66.6%)
CGP-60474 (CDK1/2/9 78.5%)
Entospletinib (SYK 0.99%)
Neratinib canonical (ERBB2 4.7%, EGFR <0.08%)
Temsirolimus (FKBP1A 96%, MTOR 53%)
Neratinib Rescued! (ITK 61.5%, MAP4K1 63.8%, LCK 95.2%)
Fostamatinib (SYK/KIT 1.6%)
Sapanisertib (MTOR/RICTOR 55%)
Crizotinib (MET 1.8%, ALK 0.06%)
BMS-536924 (IGF1R 10.2%)
NVP-BHG 712 Rescued! (LCK 95.2%, RAF1 68.4%)
Penfluridol (DRD2 <0.08%)
NVP-BHG 712 canonical (EPHB4 0.79%)
Purmorphamine (SMO 0.12%)
Ponatinib (RTKs <=1.6%)
Rebastinib (TEK/FLT3 <0.08%)
Sorafenib (VEGFR/PDGFR <=2.2%)
Max State Abundance Shift (max |log2FC| vs. DMSO)
Target Gene Detection Rate in Primary Human CD8+ T Cells (% DMSO Cells, log10 scale)
