## Extended Data Fig. 6 for "Disentangled generative modeling of single-cell perturbation responses across dynamic cellular states"

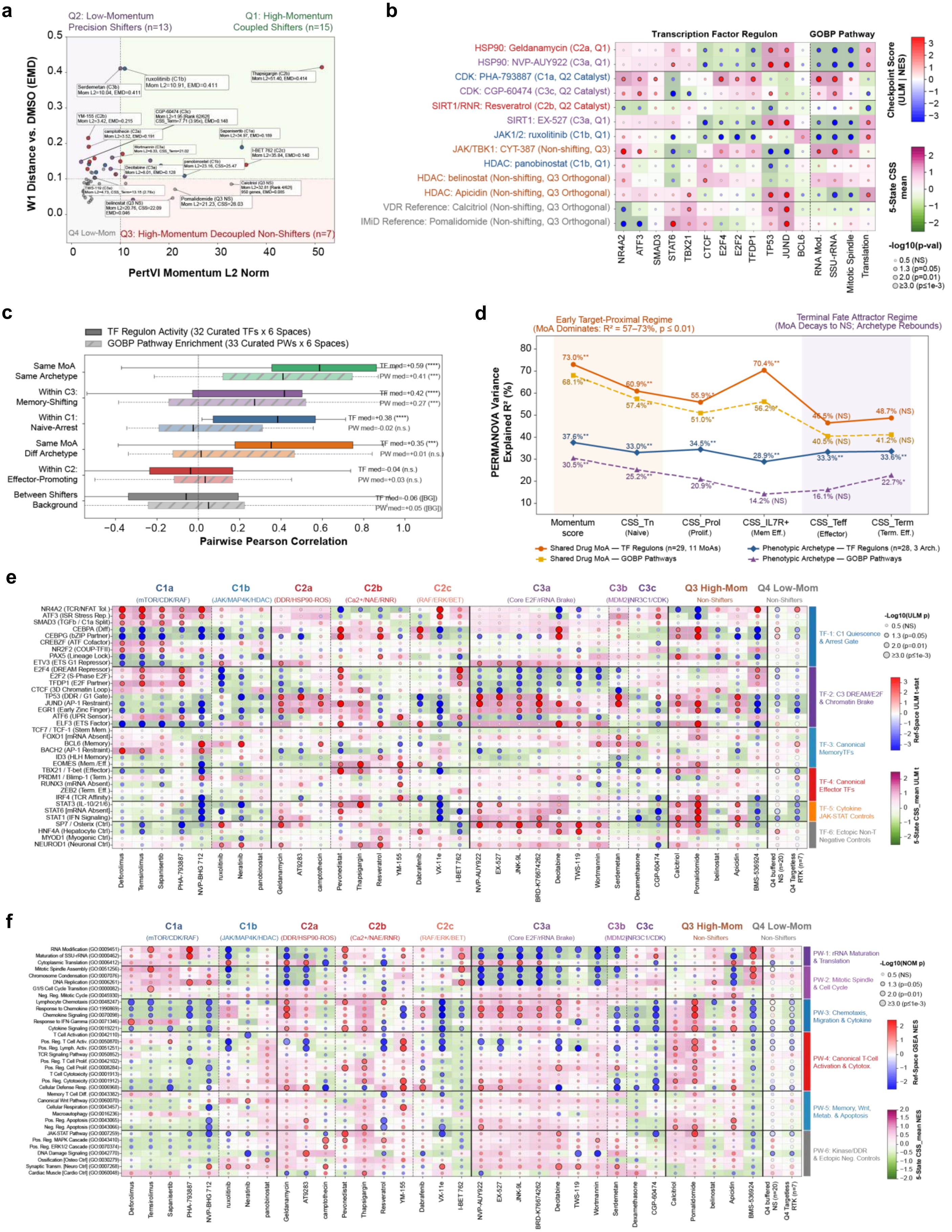

a
Q2: Low-Momentum Precision Shifters (n=13)
Q1: High-Momentum Coupled Shifters (n=15)
W1 Distance vs. DMSO (EMD)
PertVI Momentum L2 Norm
Q4 Low-Mom
Q3: High-Momentum Decoupled Non-Shifters (n=7)
Serdemetan (C3b) Mom L2=10.04, EMD=0.411
ruxolitinib (C1b) Mom L2=10.91, EMD=0.411
Thapsigargin (C2b) Mom L2=51.40, EMD=0.414
YM-155 (C2b) Mom L2=3.42, EMD=0.215
camptothecin (C2a) Mom L2=3.52, EMD=0.191
Sapanisertib (C1a) Mom L2=34.97, EMD=0.189
I-BET 762 (C2c) Mom L2=35.84, EMD=0.140
Decitabine (C3a) Mom L2=8.01, EMD=0.128
panobinostat (C1b) Mom L2=23.16, CSS=25.47
Calcitriol (Q3 NS)
Pomalidomide (Q3 NS) Mom L2=21.23, CSS=28.03
belinostat (Q3 NS) Mom L2=20.76, CSS=22.09 EMD=0.046
b
Transcription Factor Regulon
GOBP Pathway
HSP90: Geldanamycin (C2a, Q1)
HSP90: NVP-AUY922 (C3a, Q1)
CDK: PHA-793887 (C1a, Q2 Catalyst)
CDK: CGP-60474 (C3c, Q2 Catalyst)
SIRT1/RNR: Resveratrol (C2b, Q2 Catalyst)
SIRT1: EX-527 (C3a, Q1)
JAK1/2: ruxolitinib (C1b, Q1)
JAK/TBK1: CYT-387 (Non-shifting, Q3)
HDAC: panobinostat (C1b, Q1)
HDAC: belinostat (Non-shifting, Q3 Orthogonal)
HDAC: Apicidin (Non-shifting, Q3 Orthogonal)
VDR Reference: Calcitriol (Non-shifting, Q3 Orthogonal)
IMiD Reference: Pomalidomide (Non-shifting, Q3 Orthogonal)
NR4A2 ATF3 SMAD3 STAT6 TBX21 CTCF E2F4 E2F2 TFDP1 TP53 JUND BCL6 RNA Mod. SSU-rRNA Mitotic Spindle Translation
Checkpoint Score (ULM | NES)
5-State CSS mean
-log10(p-val)
0.5 (NS)
1.3 (p=0.05)
2.0 (p=0.01)
≥3.0 (p≤1e-3)
c
TF Regulon Activity (32 Curated TFs x 6 Spaces)
GOBP Pathway Enrichment (33 Curated PWs x 6 Spaces)
Same MoA Same Archetype
Within C3: Memory-Shifting
Within C1: Naive-Arrest
Same MoA Diff Archetype
Within C2: Effector-Promoting
Between Shifters Background
TF med=+0.59 (****)
PW med=+0.41 (***)
TF med=+0.42 (****)
PW med=+0.27 (***)
TF med=+0.38 (****)
PW med=-0.02 (n.s.)
TF med=+0.35 (***)
PW med=+0.01 (n.s.)
TF med=-0.04 (n.s.)
PW med=+0.03 (n.s.)
TF med=-0.06 ([BG])
PW med=+0.05 ([BG])
Pairwise Pearson Correlation
d
Early Target-Proximal Regime (MoA Dominates: R² = 57–73%, p ≤ 0.01)
Terminal Fate Attractor Regime (MoA Decays to NS; Archetype Rebounds)
PERMANOVA Variance Explained R² (%)
Momentum score
CSS_Tn (Naive)
CSS_Prol (Prolif.)
CSS_IL7R+ (Mem Eff.)
CSS_Teff (Effector)
CSS_Term (Term. Eff.)
Shared Drug MoA — TF Regulons (n=29, 11 MoAs)
Shared Drug MoA — GOBP Pathways
Phenotypic Archetype — TF Regulons (n=28, 3 Arch.)
Phenotypic Archetype — GOBP Pathways
e
C1a (mTOR/CDK/RAF)
C1b (JAK/MAP4K/HDAC)
C2a (DDR/HSP90-ROS)
C2b (Ca2+/NAE/RNR)
C2c (RAF/ERK/BET)
C3a (Core E2F/rRNA Brake)
C3b (MDM2)
C3c (NR3C1/CDK)
Q3 High-Mom Non-Shifters
Q4 Low-Mom Non-Shifters
TF-1: C1 Quiescence & Arrest Gate
TF-2: C3 DREAM/E2F & Chromatin Brake
TF-3: Canonical MemoryTFs
TF-4: Canonical Effector TFs
TF-5: Cytokine JAK-STAT Controls
TF-6: Ectopic Non-T Negative Controls
-Log10(ULM p)
Ref-Space ULM t-stat
5-State CSS_mean ULM t
Deforolimus Temsirolimus Sapanisertib PHA-793887 NVP-BHG 712 ruxolitinib Neratinib panobinostat Geldanamycin AT9283 camptothecin Pevonedistat Thapsigargin Resveratrol YM-155 Dabrafenib VX-11e I-BET 762 NVP-AUY922 EX-527 JNK-9L BRD-K76674262 Decitabine TWS-119 Wortmannin Serdemetan Dexamethasone CGP-60474 Calcitriol Pomalidomide belinostat Apicidin BMS-536924 Q4 buffered NS (n=20) Q4 Targetless RTK (n=7)
f
PW-1: rRNA Maturation & Translation
PW-2: Mitotic Spindle & Cell Cycle
PW-3: Chemotaxis, Migration & Cytokine
PW-4: Canonical T-Cell Activation & Cytotox.
PW-5: Memory, Wnt, Metab. & Apoptosis
PW-6: Kinase/DDR & Ectopic Neg. Controls
-Log10(NOM p)
Ref-Space GSEA NES
5-State CSS_mean NES
