## Extended Data Fig. 6 for "Disentangled generative modeling of single-cell perturbation responses across dynamic cellular states"

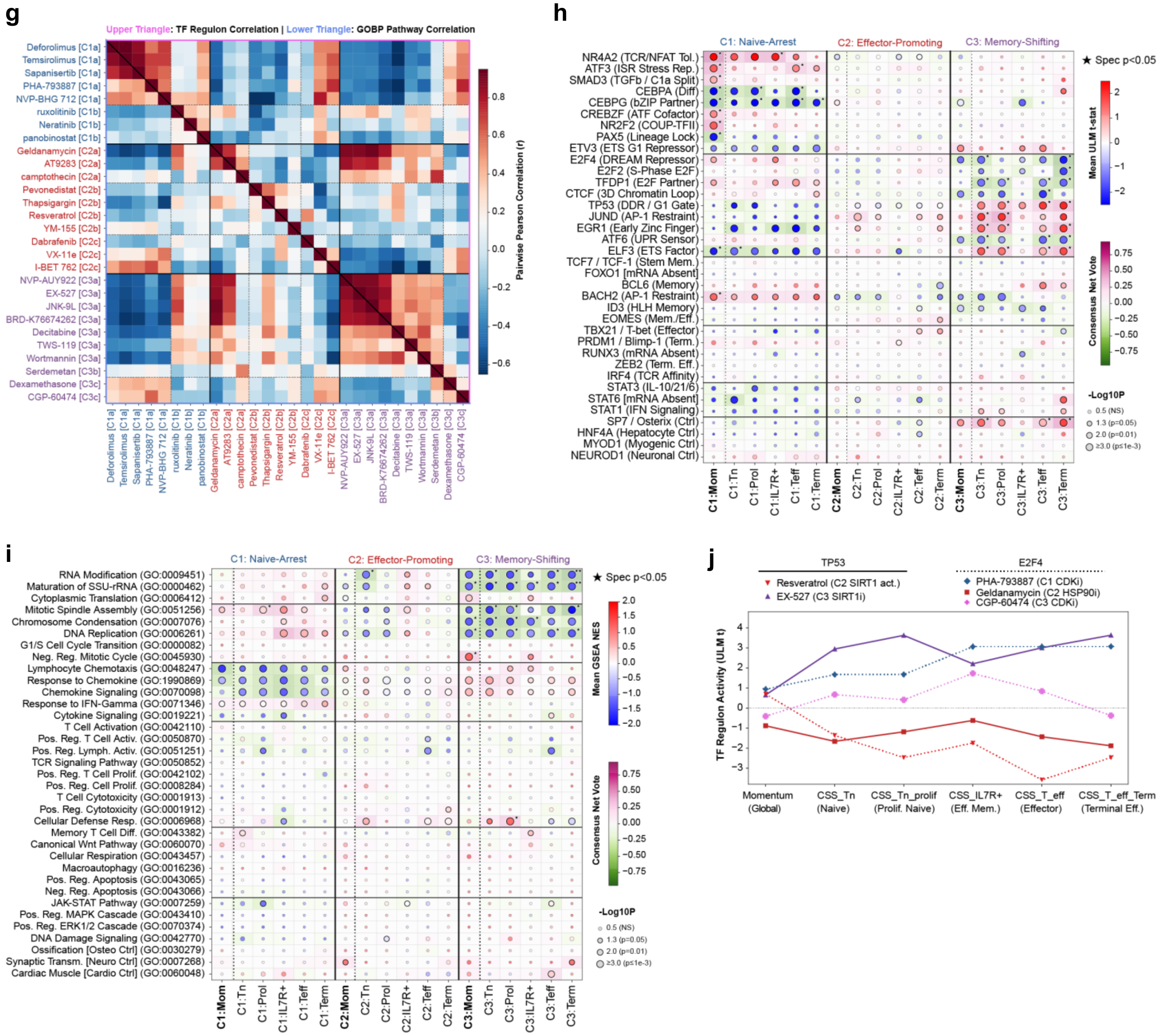
g
Upper Triangle: TF Regulon Correlation | Lower Triangle: GOBP Pathway Correlation
Deforolimus [C1a]
Temsirolimus [C1a]
Sapanisertib [C1a]
PHA-793887 [C1a]
NVP-BHG 712 [C1a]
ruxolitinib [C1b]
Neratinib [C1b]
panobinostat [C1b]
Geldanamycin [C2a]
AT9283 [C2a]
camptothecin [C2a]
Pevonedistat [C2b]
Thapsigargin [C2b]
Resveratrol [C2b]
YM-155 [C2b]
Dabrafenib [C2c]
VX-11e [C2c]
I-BET 762 [C2c]
NVP-AUY922 [C3a]
EX-527 [C3a]
JNK-9L [C3a]
BRD-K76674262 [C3a]
Decitabine [C3a]
TWS-119 [C3a]
Wortmannin [C3a]
Serdemetan [C3b]
Dexamethasone [C3c]
CGP-60474 [C3c]
Pairwise Pearson Correlation (r)
0.8
0.6
0.4
0.2
0.0
−0.2
−0.4
−0.6
h
C1: Naive-Arrest
C2: Effector-Promoting
C3: Memory-Shifting
★ Spec p<0.05
NR4A2 (TCR/NFAT Tol.)
ATF3 (ISR Stress Rep.)
SMAD3 (TGFb / C1a Split)
CEBPA (Diff)
CEBPG (bZIP Partner)
CREBZF (ATF Cofactor)
NR2F2 (COUP-TFII)
PAX5 (Lineage Lock)
ETV3 (ETS G1 Repressor)
E2F4 (DREAM Repressor)
E2F2 (S-Phase E2F)
TFDP1 (E2F Partner)
CTCF (3D Chromatin Loop)
TP53 (DDR / G1 Gate)
JUND (AP-1 Restraint)
EGR1 (Early Zinc Finger)
ATF6 (UPR Sensor)
ELF3 (ETS Factor)
TCF7 / TCF-1 (Stem Mem.)
FOXO1 [mRNA Absent]
BCL6 (Memory)
BACH2 (AP-1 Restraint)
ID3 (HLH Memory)
EOMES (Mem./Eff.)
TBX21 / T-bet (Effector)
PRDM1 / Blimp-1 (Term.)
RUNX3 (mRNA Absent)
ZEB2 (Term. Eff.)
IRF4 (TCR Affinity)
STAT3 (IL-10/21/6)
STAT6 [mRNA Absent]
STAT1 (IFN Signaling)
SP7 / Osterix (Ctrl)
HNF4A (Hepatocyte Ctrl)
MYOD1 (Myogenic Ctrl)
NEUROD1 (Neuronal Ctrl)
C1:Mom
C1:Tn
C1:Prol
C1:IL7R+
C1:Teff
C1:Term
C2:Mom
C2:Tn
C2:Prol
C2:IL7R+
C2:Teff
C2:Term
C3:Mom
C3:Tn
C3:Prol
C3:IL7R+
C3:Teff
C3:Term
Mean ULM t-stat
2
1
0
−1
−2
Consensus Net Vote
0.75
0.50
0.25
0.00
−0.25
−0.50
−0.75
-Log10P
0.5 (NS)
1.3 (p=0.05)
2.0 (p=0.01)
≥3.0 (p≤1e-3)
i
C1: Naive-Arrest
C2: Effector-Promoting
C3: Memory-Shifting
★ Spec p<0.05
RNA Modification (GO:0009451)
Maturation of SSU-rRNA (GO:0000462)
Cytoplasmic Translation (GO:0006412)
Mitotic Spindle Assembly (GO:0051256)
Chromosome Condensation (GO:0007076)
DNA Replication (GO:0006261)
G1/S Cell Cycle Transition (GO:0000082)
Neg. Reg. Mitotic Cycle (GO:0045930)
Lymphocyte Chemotaxis (GO:0048247)
Response to Chemokine (GO:1990869)
Chemokine Signaling (GO:0070098)
Response to IFN-Gamma (GO:0071346)
Cytokine Signaling (GO:0019221)
T Cell Activation (GO:0042110)
Pos. Reg. T Cell Activ. (GO:0050870)
Pos. Reg. Lymph. Activ. (GO:0051251)
TCR Signaling Pathway (GO:0050852)
Pos. Reg. T Cell Prolif. (GO:0042102)
Pos. Reg. Cell Prolif. (GO:0008284)
T Cell Cytotoxicity (GO:0001913)
Pos. Reg. Cytotoxicity (GO:0001912)
Cellular Defense Resp. (GO:0006968)
Memory T Cell Diff. (GO:0043382)
Canonical Wnt Pathway (GO:0060070)
Cellular Respiration (GO:0043457)
Macroautophagy (GO:0016236)
Pos. Reg. Apoptosis (GO:0043065)
Neg. Reg. Apoptosis (GO:0043066)
JAK-STAT Pathway (GO:0007259)
Pos. Reg. MAPK Cascade (GO:0043410)
Pos. Reg. ERK1/2 Cascade (GO:0070374)
DNA Damage Signaling (GO:0042770)
Ossification [Osteo Ctrl] (GO:0030279)
Synaptic Transm. [Neuro Ctrl] (GO:0007268)
Cardiac Muscle [Cardio Ctrl] (GO:0060048)
C1:Mom
C1:Tn
C1:Prol
C1:IL7R+
C1:Teff
C1:Term
C2:Mom
C2:Tn
C2:Prol
C2:IL7R+
C2:Teff
C2:Term
C3:Mom
C3:Tn
C3:Prol
C3:IL7R+
C3:Teff
C3:Term
Mean GSEA NES
2.0
1.5
1.0
0.5
0.0
−0.5
−1.0
−1.5
−2.0
Consensus Net Vote
0.75
0.50
0.25
0.00
−0.25
−0.50
−0.75
-Log10P
0.5 (NS)
1.3 (p=0.05)
2.0 (p=0.01)
≥3.0 (p≤1e-3)
j
TP53
Resveratrol (C2 SIRT1 act.)
EX-527 (C3 SIRT1i)
E2F4
PHA-793887 (C1 CDKi)
Geldanamycin (C2 HSP90i)
CGP-60474 (C3 CDKi)
TF Regulon Activity (ULM t)
4
3
2
1
0
−1
−2
−3
Momentum (Global)
CSS_Tn (Naive)
CSS_Tn_prolif (Prolif. Naive)
CSS_IL7R+ (Eff. Mem.)
CSS_T_eff (Effector)
CSS_T_eff_Term (Terminal Eff.)
