## Extended Data Fig. 7 for "Disentangled generative modeling of single-cell perturbation responses across dynamic cellular states"

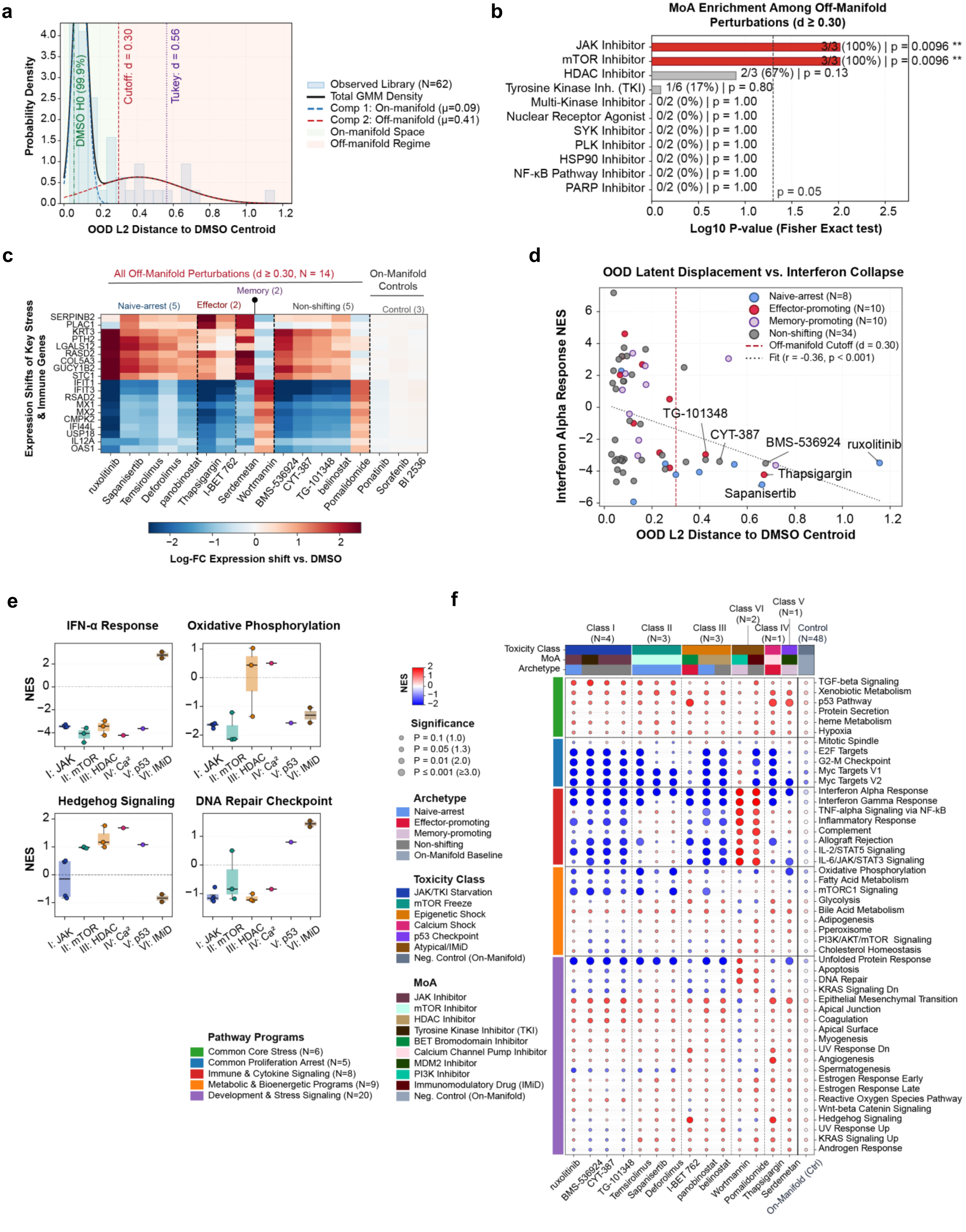
a
Probability Density
DMSO H0 (99.9%)
Cutoff: d = 0.30
Tukey: d = 0.56
Observed Library (N=62)
Total GMM Density
Comp 1: On-manifold (μ=0.09)
Comp 2: Off-manifold (μ=0.41)
On-manifold Space
Off-manifold Regime
OOD L2 Distance to DMSO Centroid
b
MoA Enrichment Among Off-Manifold Perturbations (d ≥ 0.30)
JAK Inhibitor 3/3 (100%) | p = 0.0096 **
mTOR Inhibitor 3/3 (100%) | p = 0.0096 **
HDAC Inhibitor 2/3 (67%) | p = 0.13
Tyrosine Kinase Inh. (TKI) 1/6 (17%) | p = 0.80
Multi-Kinase Inhibitor 0/2 (0%) | p = 1.00
Nuclear Receptor Agonist 0/2 (0%) | p = 1.00
SYK Inhibitor 0/2 (0%) | p = 1.00
PLK Inhibitor 0/2 (0%) | p = 1.00
HSP90 Inhibitor 0/2 (0%) | p = 1.00
NF-κB Pathway Inhibitor 0/2 (0%) | p = 1.00
PARP Inhibitor 0/2 (0%) | p = 1.00
p = 0.05
Log10 P-value (Fisher Exact test)
c
All Off-Manifold Perturbations (d ≥ 0.30, N = 14)
On-Manifold Controls
Naive-arrest (5)
Effector (2)
Memory (2)
Non-shifting (5)
Control (3)
Expression Shifts of Key Stress & Immune Genes
SERPINB2
PLAC1
KRT3
PTH2
LGALS12
RASD2
COL5A3
GUCY1B2
STC1
IFIT1
IFIT3
RSAD2
MX1
MX2
CMPK2
IFI44L
USP18
IL12A
OAS1
ruxolitinib
Sapanisertib
Temsirolimus
Deforolimus
panobinostat
Thapsigargin
I-BET 762
Serdemetan
Wortmannin
BMS-536924
CYT-387
TG-101348
belinostat
Pomalidomide
Ponatinib
Sorafenib
BI 2536
Log-FC Expression shift vs. DMSO
d
OOD Latent Displacement vs. Interferon Collapse
Interferon Alpha Response NES
Naive-arrest (N=8)
Effector-promoting (N=10)
Memory-promoting (N=10)
Non-shifting (N=34)
Off-manifold Cutoff (d = 0.30)
Fit (r = -0.36, p < 0.001)
TG-101348
CYT-387
BMS-536924
ruxolitinib
Thapsigargin
Sapanisertib
OOD L2 Distance to DMSO Centroid
e
IFN-α Response
Oxidative Phosphorylation
Hedgehog Signaling
DNA Repair Checkpoint
NES
I: JAK
II: mTOR
III: HDAC
IV: Ca²
V: p53
VI: IMiD
Pathway Programs
Common Core Stress (N=6)
Common Proliferation Arrest (N=5)
Immune & Cytokine Signaling (N=8)
Metabolic & Bioenergetic Programs (N=9)
Development & Stress Signaling (N=20)
NES
Significance
P = 0.1 (1.0)
P = 0.05 (1.3)
P = 0.01 (2.0)
P ≤ 0.001 (≥3.0)
Archetype
Naive-arrest
Effector-promoting
Memory-promoting
Non-shifting
On-Manifold Baseline
Toxicity Class
JAK/TKI Starvation
mTOR Freeze
Epigenetic Shock
Calcium Shock
p53 Checkpoint
Atypical/IMiD
Neg. Control (On-Manifold)
MoA
JAK Inhibitor
mTOR Inhibitor
HDAC Inhibitor
Tyrosine Kinase Inhibitor (TKI)
BET Bromodomain Inhibitor
Calcium Channel Pump Inhibitor
MDM2 Inhibitor
PI3K Inhibitor
Immunomodulatory Drug (IMiD)
Neg. Control (On-Manifold)
f
Class I (N=4)
Class II (N=3)
Class III (N=3)
Class VI (N=2)
Class IV (N=1)
Class V (N=1)
Control (N=48)
Toxicity Class
MoA
Archetype
TGF-beta Signaling
Xenobiotic Metabolism
p53 Pathway
Protein Secretion
heme Metabolism
Hypoxia
Mitotic Spindle
E2F Targets
G2-M Checkpoint
Myc Targets V1
Myc Targets V2
Interferon Alpha Response
Interferon Gamma Response
TNF-alpha Signaling via NF-kB
Inflammatory Response
Complement
Allograft Rejection
IL-2/STAT5 Signaling
IL-6/JAK/STAT3 Signaling
Oxidative Phosphorylation
Fatty Acid Metabolism
mTORC1 Signaling
Glycolysis
Bile Acid Metabolism
Adipogenesis
Pperoxisome
PI3K/AKT/mTOR Signaling
Cholesterol Homeostasis
Unfolded Protein Response
Apoptosis
DNA Repair
KRAS Signaling Dn
Epithelial Mesenchymal Transition
Apical Junction
Coagulation
Apical Surface
Myogenesis
UV Response Dn
Angiogenesis
Spermatogenesis
Estrogen Response Early
Estrogen Response Late
Reactive Oxygen Species Pathway
Wnt-beta Catenin Signaling
Hedgehog Signaling
UV Response Up
KRAS Signaling Up
Androgen Response
ruxolitinib
BMS-536924
CYT-387
TG-101348
Temsirolimus
Sapanisertib
Deforolimus
I-BET 762
panobinostat
belinostat
Wortmannin
Pomalidomide
Thapsigargin
Serdemetan
On-Manifold (Ctrl)
