## Supplementary figures and images for "Disentangled generative modeling of single-cell perturbation responses across dynamic cellular states"

### Extended Data Fig. 8

**a**

| Donor | Rest | Stim 8hr | Stim 48hr |
|---|---|---|---|
| D1 | 9,537 | 8,688 | 7,526 |
| D2 | 14,304 | 14,124 | 12,705 |
| D3 | 9,43 | 8,441 | 8,707 |
| D4 | 9,160 | 9,019 | 8,991 |

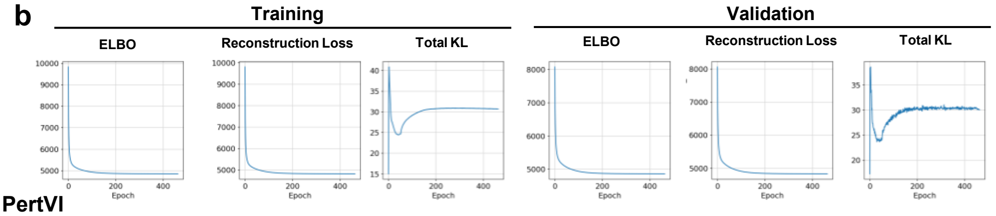


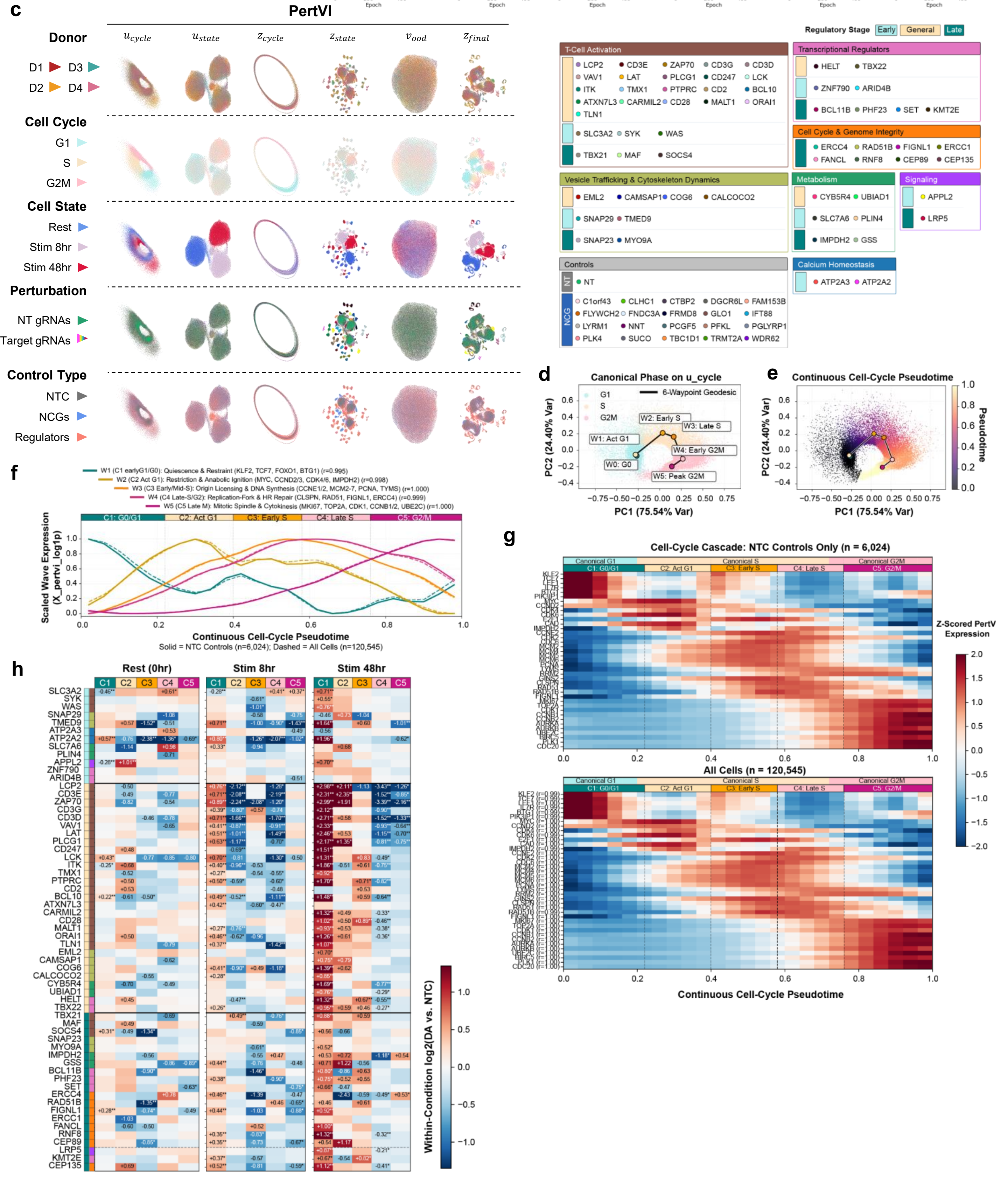
