## Extended Data Fig. 9 for "Disentangled generative modeling of single-cell perturbation responses across dynamic cellular states"

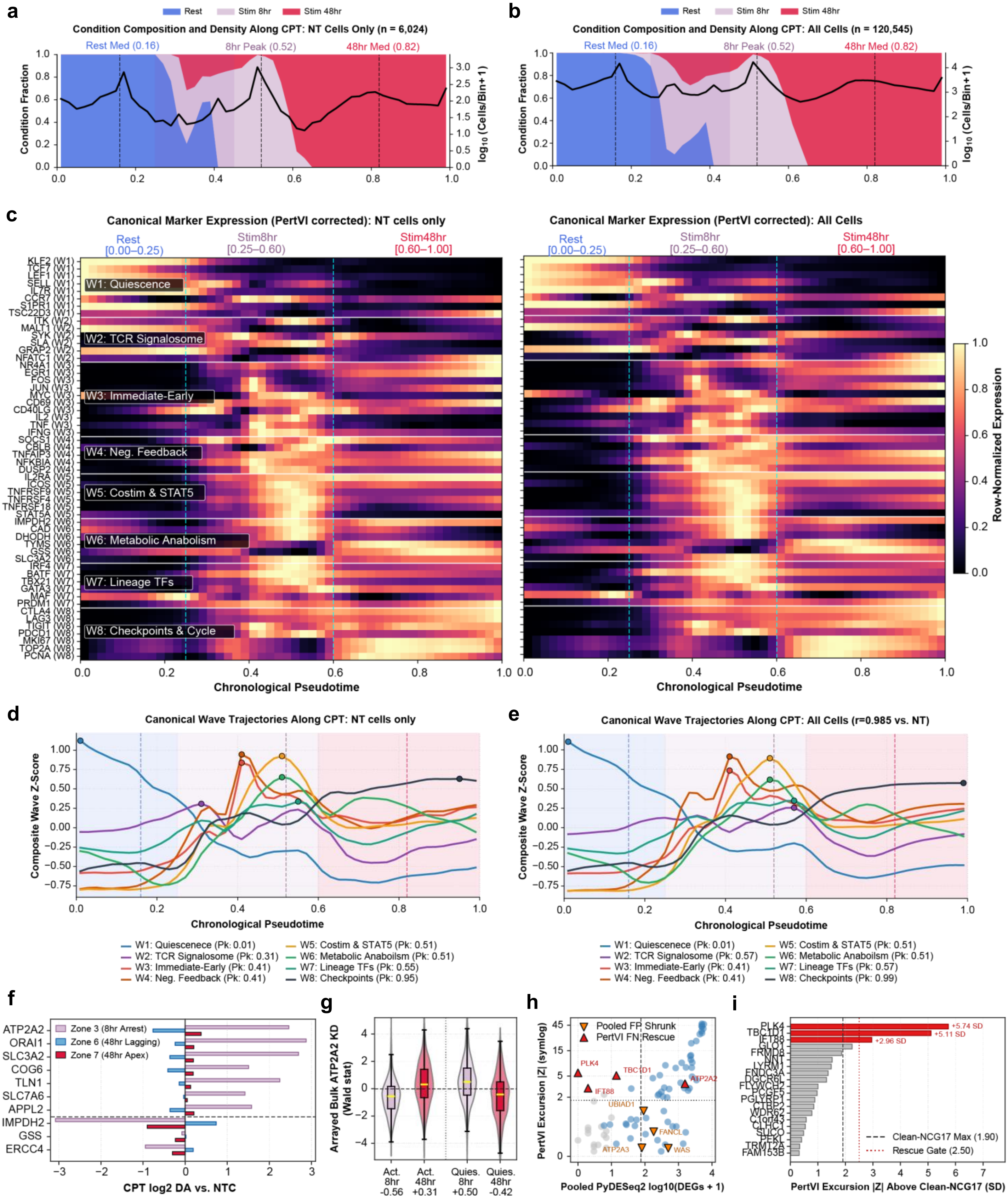
a
Rest
Stim 8hr
Stim 48hr
Condition Composition and Density Along CPT: NT Cells Only (n = 6,024)
Rest Med (0.16)
8hr Peak (0.52)
48hr Med (0.82)
Condition Fraction
log10 (Cells/Bin+1)
b
Condition Composition and Density Along CPT: All Cells (n = 120,545)
c
Canonical Marker Expression (PertVI corrected): NT cells only
Canonical Marker Expression (PertVI corrected): All Cells
Rest [0.00–0.25]
Stim8hr [0.25–0.60]
Stim48hr [0.60–1.00]
KLF2 (W1)
TCF7 (W1)
LEF1 (W1)
SELL (W1)
IL7R (W1)
CCR7 (W1)
S1PR1 (W1)
TSC22D3 (W1)
ITK (W2)
MALT1 (W2)
SYK (W2)
SLA (W2)
GRAP2 (W2)
NFATC1 (W2)
NR4A1 (W3)
EGR1 (W3)
FOS (W3)
JUN (W3)
MYC (W3)
CD69 (W3)
CD40LG (W3)
IL2 (W3)
TNF (W3)
IFNG (W3)
SOCS1 (W4)
CBLB (W4)
TNFAIP3 (W4)
NFKBIA (W4)
DUSP2 (W4)
IL2RA (W5)
ICOS (W5)
TNFRSF9 (W5)
TNFRSF4 (W5)
TNFRSF18 (W5)
STAT5A (W5)
IMPDH2 (W6)
CAD (W6)
DHODH (W6)
TYMS (W6)
GSS (W6)
SLC3A2 (W6)
IRF4 (W7)
BATF (W7)
TBX21 (W7)
GATA3 (W7)
MAF (W7)
PRDM1 (W7)
CTLA4 (W8)
LAG3 (W8)
TIGIT (W8)
PDCD1 (W8)
MKI67 (W8)
TOP2A (W8)
PCNA (W8)
W1: Quiescence
W2: TCR Signalosome
W3: Immediate-Early
W4: Neg. Feedback
W5: Costim & STAT5
W6: Metabolic Anabolism
W7: Lineage TFs
W8: Checkpoints & Cycle
Chronological Pseudotime
Row-Normalized Expression
d
Canonical Wave Trajectories Along CPT: NT cells only
Composite Wave Z-Score
W1: Quiescenece (Pk: 0.01)
W2: TCR Signalosome (Pk: 0.31)
W3: Immediate-Early (Pk: 0.41)
W4: Neg. Feedback (Pk: 0.41)
W5: Costim & STAT5 (Pk: 0.51)
W6: Metabolic Anaboilsm (Pk: 0.51)
W7: Lineage TFs (Pk: 0.55)
W8: Checkpoints (Pk: 0.95)
e
Canonical Wave Trajectories Along CPT: All Cells (r=0.985 vs. NT)
W1: Quiescenece (Pk: 0.01)
W2: TCR Signalosome (Pk: 0.57)
W3: Immediate-Early (Pk: 0.41)
W4: Neg. Feedback (Pk: 0.41)
W5: Costim & STAT5 (Pk: 0.51)
W6: Metabolic Anaboilsm (Pk: 0.51)
W7: Lineage TFs (Pk: 0.57)
W8: Checkpoints (Pk: 0.99)
f
Zone 3 (8hr Arrest)
Zone 6 (48hr Lagging)
Zone 7 (48hr Apex)
ATP2A2
ORAI1
SLC3A2
COG6
TLN1
SLC7A6
APPL2
IMPDH2
GSS
ERCC4
CPT log2 DA vs. NTC
g
Arrayed Bulk ATP2A2 KD (Wald stat)
Act. 8hr -0.56
Act. 48hr +0.31
Quies. 8hr +0.50
Quies. 48hr -0.42
h
Pooled FP Shrunk
PertVI FN Rescue
PLK4
TBC1D1
IFT88
ATP2A2
UBIAD1
FANCL
ATP2A3
WAS
PertVI Excursion |Z| (symlog)
Pooled PyDESeq2 log10(DEGs + 1)
i
PLK4
TBC1D1
IFT88
GLO1
FRMD8
NIN1
LYRM4
ENDOG
DGCR6L
FLYWCH2
PCGF5
PGLYRP1
CTBP2
WDR62
C1orf43
SLCO
CLHC1
PFKL
TRMT2A
FAM153B
+5.74 SD
+5.11 SD
+2.96 SD
Clean-NCG17 Max (1.90)
Rescue Gate (2.50)
PertVI Excursion |Z| Above Clean-NCG17 (SD)
