## Extended Data Fig. 10 for "Disentangled generative modeling of single-cell perturbation responses across dynamic cellular states"

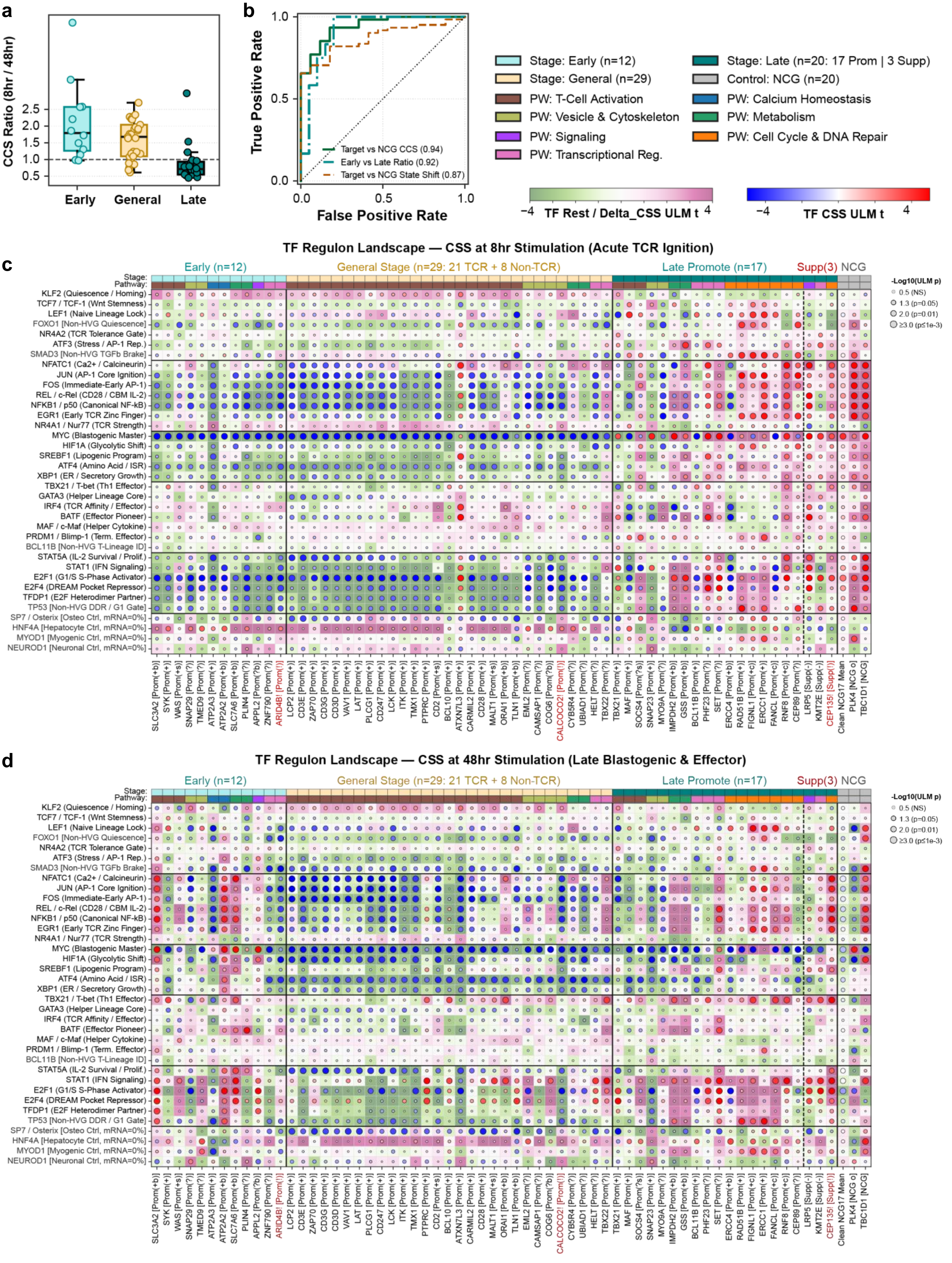
a
CCS Ratio (8hr / 48hr)
2.5
2.0
1.5
1.0
0.5
Early
General
Late
b
True Positive Rate
False Positive Rate
1.0
0.5
0.0
Target vs NCG CCS (0.94)
Early vs Late Ratio (0.92)
Target vs NCG State Shift (0.87)
Stage: Early (n=12)
Stage: General (n=29)
PW: T-Cell Activation
PW: Vesicle & Cytoskeleton
PW: Signaling
PW: Transcriptional Reg.
Stage: Late (n=20: 17 Prom | 3 Supp)
Control: NCG (n=20)
PW: Calcium Homeostasis
PW: Metabolism
PW: Cell Cycle & DNA Repair
−4 TF Rest / Delta_CSS ULM t 4
−4 TF CSS ULM t 4
TF Regulon Landscape — CSS at 8hr Stimulation (Acute TCR Ignition)
c
Early (n=12)
General Stage (n=29: 21 TCR + 8 Non-TCR)
Late Promote (n=17)
Supp(3)
NCG
Stage:
Pathway:
-Log10(ULM p)
0.5 (NS)
1.3 (p=0.05)
2.0 (p=0.01)
≥3.0 (p≤1e-3)
KLF2 (Quiescence / Homing)
TCF7 / TCF-1 (Wnt Stemness)
LEF1 (Naive Lineage Lock)
FOXO1 [Non-HVG Quiescence]
NR4A2 (TCR Tolerance Gate)
ATF3 (Stress / AP-1 Rep.)
SMAD3 [Non-HVG TGFb Brake]
NFATC1 (Ca2+ / Calcineurin)
JUN (AP-1 Core Ignition)
FOS (Immediate-Early AP-1)
REL / c-Rel (CD28 / CBM IL-2)
NFKB1 / p50 (Canonical NF-kB)
EGR1 (Early TCR Zinc Finger)
NR4A1 / Nur77 (TCR Strength)
MYC (Blastogenic Master)
HIF1A (Glycolytic Shift)
SREBF1 (Lipogenic Program)
ATF4 (Amino Acid / ISR)
XBP1 (ER / Secretory Growth)
TBX21 / T-bet (Th1 Effector)
GATA3 (Helper Lineage Core)
IRF4 (TCR Affinity / Effector)
BATF (Effector Pioneer)
MAF / c-Maf (Helper Cytokine)
PRDM1 / Blimp-1 (Term. Effector)
BCL11B [Non-HVG T-Lineage ID]
STAT5A (IL-2 Survival / Prolif.)
STAT1 (IFN Signaling)
E2F1 (G1/S S-Phase Activator)
E2F4 (DREAM Pocket Repressor)
TFDP1 (E2F Heterodimer Partner)
TP53 [Non-HVG DDR / G1 Gate]
SP7 / Osterix [Osteo Ctrl, mRNA=0%]
HNF4A [Hepatocyte Ctrl, mRNA=0%]
MYOD1 [Myogenic Ctrl, mRNA=0%]
NEUROD1 [Neuronal Ctrl, mRNA=0%]
SLC3A2 [Prom(+b)]
SYK [Prom(+b)]
WAS [Prom(+s)]
SNAP29 [Prom(?)]
TMED9 [Prom(?)]
ATP2A3 [Prom(+)]
ATP2A2 [Prom(+b)]
SLC7A6 [Prom(+b)]
PLIN4 [Prom(?)]
APPL2 [Prom(?b)]
ZNF790 [Prom(?)]
ARID4B! [Prom(!)]
LCP2 [Prom(+)]
CD3E [Prom(+)]
ZAP70 [Prom(+)]
CD3G [Prom(+)]
CD3D [Prom(+)]
VAV1 [Prom(+)]
LAT [Prom(+)]
PLCG1 [Prom(+)]
CD247 [Prom(+)]
LCK [Prom(+)]
ITK [Prom(+)]
TMX1 [Prom(+)]
PTPRC [Prom(+)]
CD2 [Prom(+s)]
BCL10 [Prom(+)]
ATXN7L3 [Prom(+)]
CARMIL2 [Prom(+)]
CD28 [Prom(+)]
MALT1 [Prom(+s)]
ORAI1 [Prom(+b)]
TLN1 [Prom(+b)]
EML2 [Prom(?)]
CAMSAP1 [Prom(?b)]
COG6 [Prom(?b)]
CALCOCO2! [Prom(!)]
CYB5R4 [Prom(?)]
UBIAD1 [Prom(?)]
HELT [Prom(?)]
TBX22 [Prom(?)]
TBX21 [Prom(+)]
MAF [Prom(+)]
SOCS4 [Prom(?s)]
SNAP23 [Prom(+)]
MYO9A [Prom(+)]
IMPDH2 [Prom(+b)]
GSS [Prom(+b)]
BCL11B [Prom(+)]
PHF23 [Prom(?)]
SET [Prom(?)]
ERCC4 [Prom(+b)]
RAD51B [Prom(+)]
FIGNL1 [Prom(+c)]
ERCC1 [Prom(+)]
FANCL [Prom(+c)]
RNF8 [Prom(+c)]
CEP89 [Prom(?)]
LRP5 [Supp(-)]
KMT2E [Supp(-)]
CEP135! [Supp(!)]
Clean NCG17 Mean
PLK4 [NCG o]
TBC1D1 [NCG]
TF Regulon Landscape — CSS at 48hr Stimulation (Late Blastogenic & Effector)
d
Early (n=12)
General Stage (n=29: 21 TCR + 8 Non-TCR)
Late Promote (n=17)
Supp(3)
NCG
Stage:
Pathway:
-Log10(ULM p)
0.5 (NS)
1.3 (p=0.05)
2.0 (p=0.01)
≥3.0 (p≤1e-3)
