## Extended Data Fig. 10 for "Disentangled generative modeling of single-cell perturbation responses across dynamic cellular states"

**e**

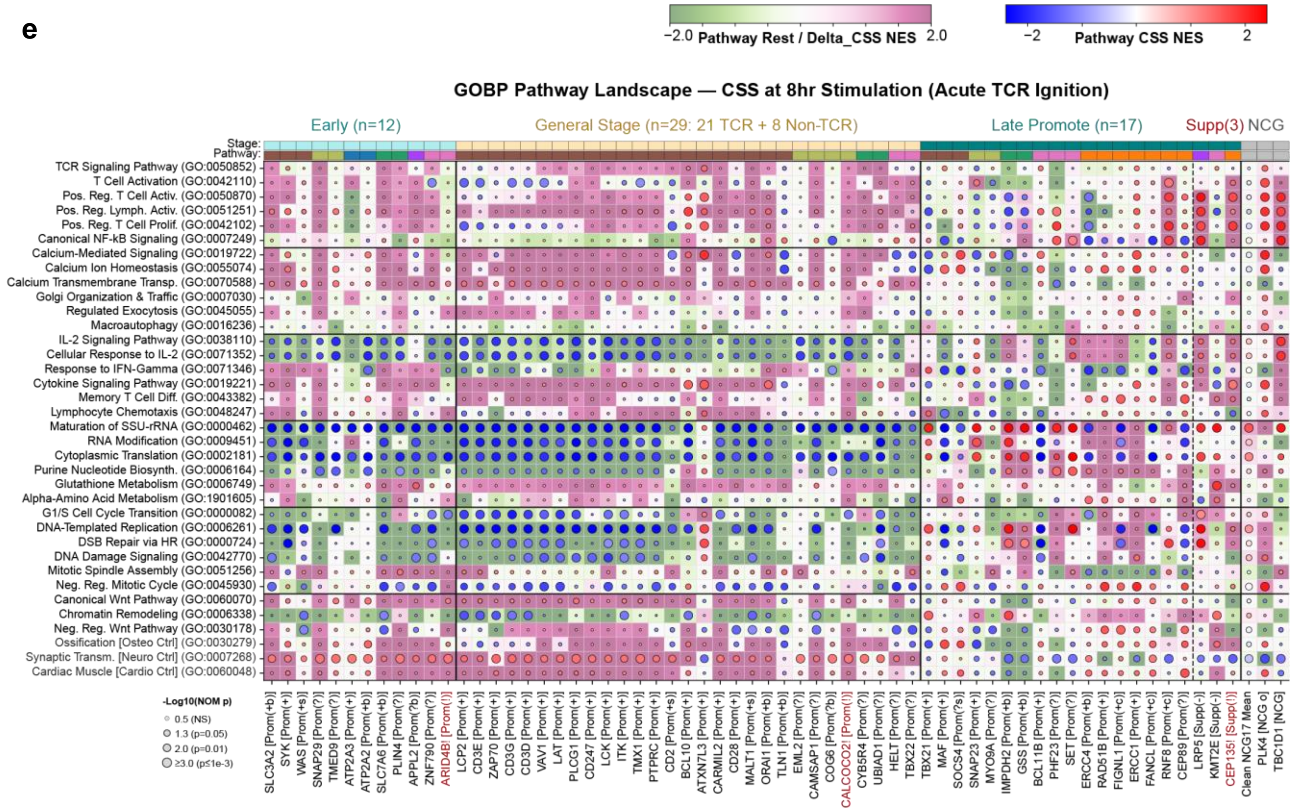
-2.0 Pathway Rest / Delta_CSS NES 2.0
-2 Pathway CSS NES 2
GOBP Pathway Landscape — CSS at 8hr Stimulation (Acute TCR Ignition)
Early (n=12)
General Stage (n=29: 21 TCR + 8 Non-TCR)
Late Promote (n=17)
Supp(3)
NCG
Stage:
Pathway:
TCR Signaling Pathway (GO:0050852)
T Cell Activation (GO:0042110)
Pos. Reg. T Cell Activ. (GO:0050870)
Pos. Reg. Lymph. Activ. (GO:0051251)
Pos. Reg. T Cell Prolif. (GO:0042102)
Canonical NF-kB Signaling (GO:0007249)
Calcium-Mediated Signaling (GO:0019722)
Calcium Ion Homeostasis (GO:0055074)
Calcium Transmembrane Transp. (GO:0070588)
Golgi Organization & Traffic (GO:0007030)
Regulated Exocytosis (GO:0045055)
Macroautophagy (GO:0016236)
IL-2 Signaling Pathway (GO:0038110)
Cellular Response to IL-2 (GO:0071352)
Response to IFN-Gamma (GO:0071346)
Cytokine Signaling Pathway (GO:0019221)
Memory T Cell Diff. (GO:0043382)
Lymphocyte Chemotaxis (GO:0048247)
Maturation of SSU-rRNA (GO:0000462)
RNA Modification (GO:0009451)
Cytoplasmic Translation (GO:0002181)
Purine Nucleotide Biosynth. (GO:0006164)
Glutathione Metabolism (GO:0006749)
Alpha-Amino Acid Metabolism (GO:1901605)
G1/S Cell Cycle Transition (GO:0000082)
DNA-Templated Replication (GO:0006261)
DSB Repair via HR (GO:0000724)
DNA Damage Signaling (GO:0042770)
Mitotic Spindle Assembly (GO:0051256)
Neg. Reg. Mitotic Cycle (GO:0045930)
Canonical Wnt Pathway (GO:0060070)
Chromatin Remodeling (GO:0006338)
Neg. Reg. Wnt Pathway (GO:0030178)
Ossification [Osteo Ctrl] (GO:0030279)
Synaptic Transm. [Neuro Ctrl] (GO:0007268)
Cardiac Muscle [Cardio Ctrl] (GO:0060048)
-Log10(NOM p)
0.5 (NS)
1.3 (p=0.05)
2.0 (p=0.01)
≥3.0 (p≤1e-3)
SLC3A2 [Prom(+b)]
SYK [Prom(+)]
WAS [Prom(+s)]
SNAP29 [Prom(?)]
TMED9 [Prom(?)]
ATP2A3 [Prom(+)]
ATP2A2 [Prom(+b)]
SLC7A6 [Prom(+b)]
PLIN4 [Prom(?)]
APPL2 [Prom(?b)]
ZNF790 [Prom(?)]
ARID4B! [Prom(!)]
LCP2 [Prom(+)]
CD3E [Prom(+)]
ZAP70 [Prom(+)]
CD3G [Prom(+)]
CD3D [Prom(+)]
VAV1 [Prom(+)]
LAT [Prom(+)]
PLCG1 [Prom(+)]
CD247 [Prom(+)]
LCK [Prom(+)]
ITK [Prom(+)]
TMX1 [Prom(+)]
PTPRC [Prom(+)]
CD2 [Prom(+s)]
BCL10 [Prom(+)]
ATXN7L3 [Prom(+)]
CARMIL2 [Prom(+)]
CD28 [Prom(+)]
MALT1 [Prom(+s)]
ORAI1 [Prom(+b)]
TLN1 [Prom(+b)]
EML2 [Prom(?)]
CAMSAP1 [Prom(?)]
COG6 [Prom(?b)]
CALCOCO2! [Prom(!)]
CYB5R4 [Prom(?)]
UBIAD1 [Prom(?)]
HELT [Prom(?)]
TBX22 [Prom(?)]
TBX21 [Prom(+)]
MAF [Prom(+)]
SOCS4 [Prom(?s)]
SNAP23 [Prom(+)]
MYO9A [Prom(?)]
IMPDH2 [Prom(+b)]
GSS [Prom(+b)]
BCL11B [Prom(+)]
PHF23 [Prom(?)]
SET [Prom(?)]
ERCC4 [Prom(+b)]
RAD51B [Prom(+)]
FIGNL1 [Prom(+c)]
ERCC1 [Prom(+)]
FANCL [Prom(+c)]
RNF8 [Prom(+c)]
CEP89 [Prom(?)]
LRP5 [Supp(-)]
KMT2E [Supp(-)]
CEP135! [Supp(!)]
Clean NCG17 Mean
PLK4 [NCG o]
TBC1D1 [NCG]

**f**

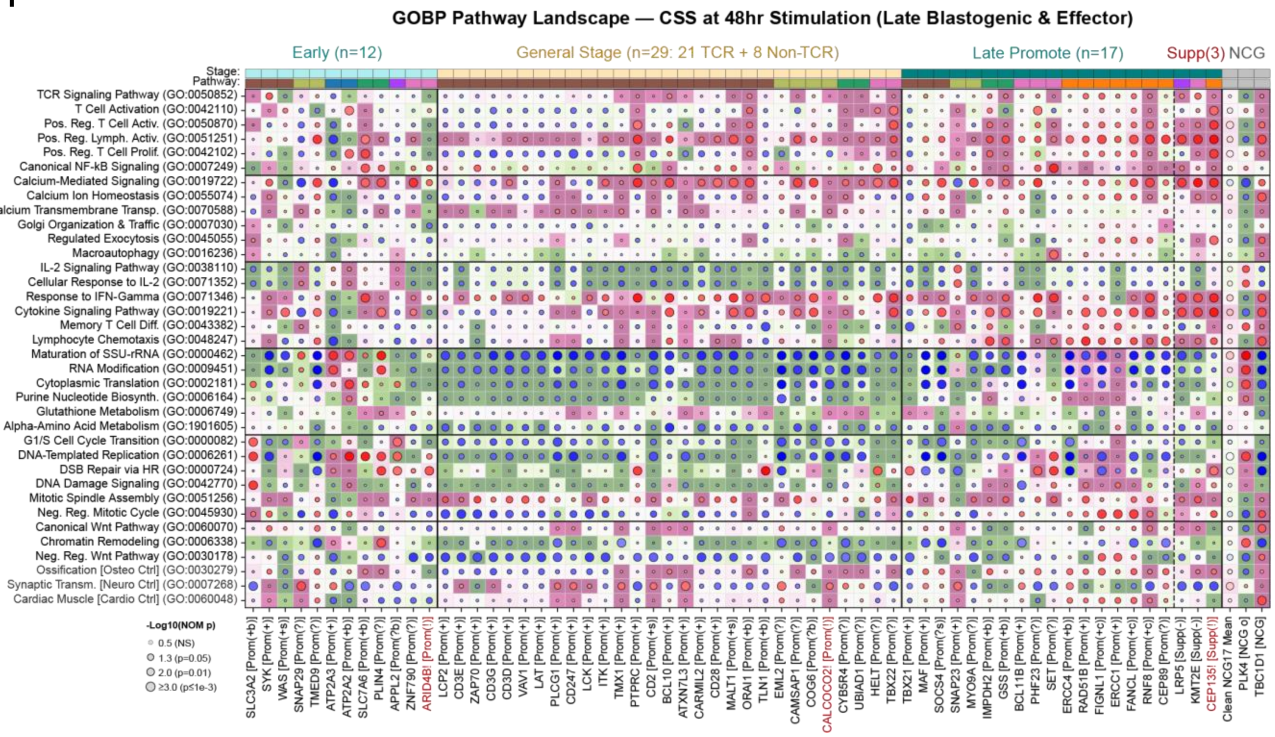
GOBP Pathway Landscape — CSS at 48hr Stimulation (Late Blastogenic & Effector)
Early (n=12)
General Stage (n=29: 21 TCR + 8 Non-TCR)
Late Promote (n=17)
Supp(3)
NCG
Stage:
Pathway:
TCR Signaling Pathway (GO:0050852)
T Cell Activation (GO:0042110)
Pos. Reg. T Cell Activ. (GO:0050870)
Pos. Reg. Lymph. Activ. (GO:0051251)
Pos. Reg. T Cell Prolif. (GO:0042102)
Canonical NF-kB Signaling (GO:0007249)
Calcium-Mediated Signaling (GO:0019722)
Calcium Ion Homeostasis (GO:0055074)
Calcium Transmembrane Transp. (GO:0070588)
Golgi Organization & Traffic (GO:0007030)
Regulated Exocytosis (GO:0045055)
Macroautophagy (GO:0016236)
IL-2 Signaling Pathway (GO:0038110)
Cellular Response to IL-2 (GO:0071352)
Response to IFN-Gamma (GO:0071346)
Cytokine Signaling Pathway (GO:0019221)
Memory T Cell Diff. (GO:0043382)
Lymphocyte Chemotaxis (GO:0048247)
Maturation of SSU-rRNA (GO:0000462)
RNA Modification (GO:0009451)
Cytoplasmic Translation (GO:0002181)
Purine Nucleotide Biosynth. (GO:0006164)
Glutathione Metabolism (GO:0006749)
Alpha-Amino Acid Metabolism (GO:1901605)
G1/S Cell Cycle Transition (GO:0000082)
DNA-Templated Replication (GO:0006261)
DSB Repair via HR (GO:0000724)
DNA Damage Signaling (GO:0042770)
Mitotic Spindle Assembly (GO:0051256)
Neg. Reg. Mitotic Cycle (GO:0045930)
Canonical Wnt Pathway (GO:0060070)
Chromatin Remodeling (GO:0006338)
Neg. Reg. Wnt Pathway (GO:0030178)
Ossification [Osteo Ctrl] (GO:0030279)
Synaptic Transm. [Neuro Ctrl] (GO:0007268)
Cardiac Muscle [Cardio Ctrl] (GO:0060048)
-Log10(NOM p)
0.5 (NS)
1.3 (p=0.05)
2.0 (p=0.01)
≥3.0 (p≤1e-3)
SLC3A2 [Prom(+b)]
SYK [Prom(+)]
WAS [Prom(+s)]
SNAP29 [Prom(?)]
TMED9 [Prom(?)]
ATP2A3 [Prom(+)]
ATP2A2 [Prom(+b)]
SLC7A6 [Prom(+b)]
PLIN4 [Prom(?)]
APPL2 [Prom(?b)]
ZNF790 [Prom(?)]
ARID4B! [Prom(!)]
LCP2 [Prom(+)]
CD3E [Prom(+)]
ZAP70 [Prom(+)]
CD3G [Prom(+)]
CD3D [Prom(+)]
VAV1 [Prom(+)]
LAT [Prom(+)]
PLCG1 [Prom(+)]
CD247 [Prom(+)]
LCK [Prom(+)]
ITK [Prom(+)]
TMX1 [Prom(+)]
PTPRC [Prom(+)]
CD2 [Prom(+s)]
BCL10 [Prom(+)]
ATXN7L3 [Prom(+)]
CARMIL2 [Prom(+)]
CD28 [Prom(+)]
MALT1 [Prom(+s)]
ORAI1 [Prom(+b)]
TLN1 [Prom(+b)]
EML2 [Prom(?)]
CAMSAP1 [Prom(?)]
COG6 [Prom(?b)]
CALCOCO2! [Prom(!)]
CYB5R4 [Prom(?)]
UBIAD1 [Prom(?)]
HELT [Prom(?)]
TBX22 [Prom(?)]
TBX21 [Prom(+)]
MAF [Prom(+)]
SOCS4 [Prom(?s)]
SNAP23 [Prom(+)]
MYO9A [Prom(?)]
IMPDH2 [Prom(+b)]
GSS [Prom(+b)]
BCL11B [Prom(+)]
PHF23 [Prom(?)]
SET [Prom(?)]
ERCC4 [Prom(+b)]
RAD51B [Prom(+)]
FIGNL1 [Prom(+c)]
ERCC1 [Prom(+)]
FANCL [Prom(+c)]
RNF8 [Prom(+c)]
CEP89 [Prom(?)]
LRP5 [Supp(-)]
KMT2E [Supp(-)]
CEP135! [Supp(!)]
Clean NCG17 Mean
PLK4 [NCG o]
TBC1D1 [NCG]
