## Extended Data Fig. 11 for "Disentangled generative modeling of single-cell perturbation responses across dynamic cellular states"

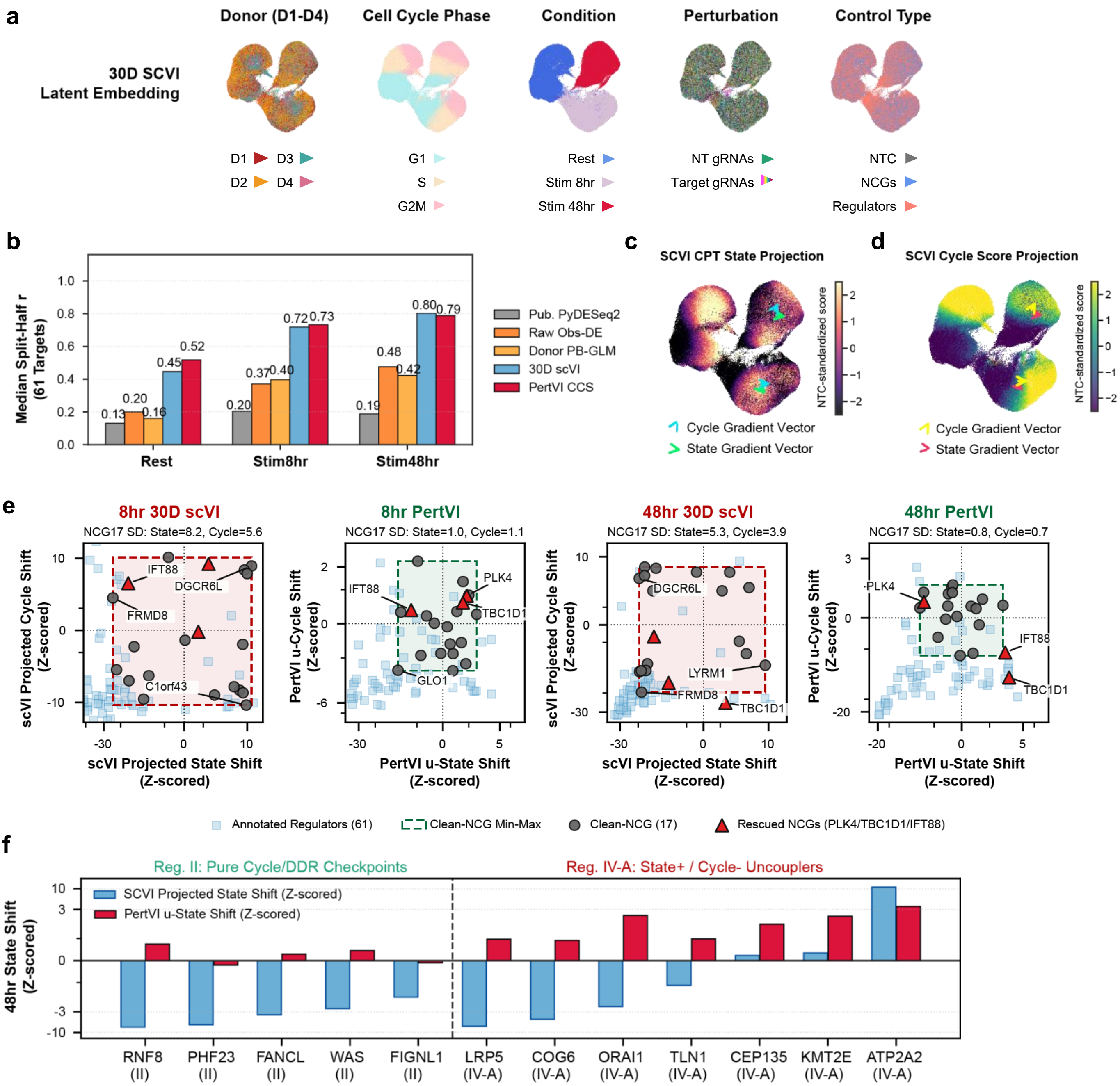
a
Donor (D1-D4)
Cell Cycle Phase
Condition
Perturbation
Control Type
30D SCVI
Latent Embedding
D1
D3
D2
D4
G1
S
G2M
Rest
Stim 8hr
Stim 48hr
NT gRNAs
Target gRNAs
NTC
NCGs
Regulators
b
Median Split-Half r
(61 Targets)
Pub. PyDESeq2
Raw Obs-DE
Donor PB-GLM
30D scVI
PertVI CCS
Rest
Stim8hr
Stim48hr
c
SCVI CPT State Projection
NTC-standardized score
Cycle Gradient Vector
State Gradient Vector
d
SCVI Cycle Score Projection
NTC-standardized score
Cycle Gradient Vector
State Gradient Vector
e
8hr 30D scVI
NCG17 SD: State=8.2, Cycle=5.6
8hr PertVI
NCG17 SD: State=1.0, Cycle=1.1
48hr 30D scVI
NCG17 SD: State=5.3, Cycle=3.9
48hr PertVI
NCG17 SD: State=0.8, Cycle=0.7
scVI Projected Cycle Shift
(Z-scored)
PertVI u-Cycle Shift
(Z-scored)
scVI Projected State Shift
(Z-scored)
PertVI u-State Shift
(Z-scored)
IFT88
DGCR6L
FRMD8
C1orf43
PLK4
TBC1D1
GLO1
LYRM1
Annotated Regulators (61)
Clean-NCG Min-Max
Clean-NCG (17)
Rescued NCGs (PLK4/TBC1D1/IFT88)
f
Reg. II: Pure Cycle/DDR Checkpoints
Reg. IV-A: State+ / Cycle- Uncouplers
SCVI Projected State Shift (Z-scored)
PertVI u-State Shift (Z-scored)
48hr State Shift
(Z-scored)
RNF8 (II)
PHF23 (II)
FANCL (II)
WAS (II)
FIGNL1 (II)
LRP5 (IV-A)
COG6 (IV-A)
ORAI1 (IV-A)
TLN1 (IV-A)
CEP135 (IV-A)
KMT2E (IV-A)
ATP2A2 (IV-A)
